# Exploratory Data Analysis in a Six-Year Longitudinal Study in Healthy Brain Aging

**DOI:** 10.1101/674853

**Authors:** Jaime Gómez-Ramírez, Marina Ávila Villanueva, Belén Frades Payo, Teodoro del Ser Quijano, Meritxell Valentí Soler, María Ascensión Zea Sevilla, Miguel Ángel Fernández-Blázquez

## Abstract

Alzheimer’s Disease (AD) is a complex, multifactorial and comorbid condition. The asymptomatic behavior in early stages of the disease is a paramount obstacle to formulate a preclinical and predictive model of AD. Not surprisingly, the AD drug approval rate is one of the lowest in the industry, an exiguous 0.4%. The identification of risk factors, preferably obtained by the subject herself, is sorely needed given that the incidence of Alzheimer’s disease grows exponentially with age [Ferri et al., 2005], [Ganguli and Rodriguez, 2011].

During the last 7 years, researchers at *Proyecto Vallecas* have collected information about the project’s volunteers, aged 70 or more. The *Proyecto Vallecas* dataset includes information about a wide range of factors including magnetic resonance imaging, genetic, demographic, socioeconomic, cognitive performance, subjective memory complaints, neuropsychiatric disorders, cardiovascular, sleep, diet, physical exercise and self assessed quality of life. The subjects in each visit were diagnosed as healthy, mild cognitive impairment (MCI) or dementia.

In this study we perform Exploratory Data Analysis to summarize the main characteristics of this unique longitudinal dataset. The objective is to characterize the evolution of the collected features over time and most importantly, how their dynamics are related to cognitive decline. We show that the longitudinal dataset of *Proyecto Vallecas*, if conveniently exploited, holds promise to identifying either factors promoting healthy aging and risk factors related to cognitive decline.

## 1 Introduction

The Vallecas Project^1^ for early detection of AD is the most ambitious population-based study for healthy aging in Spain. The project is carried out at the Queen Sofia Foundation Alzheimer Center by a multidisciplinary team of researchers from the CIEN Foundation. The main objective of the Vallecas Project is to elucidate, through tracking of progression of the cohort, the best combination of features, clinical and others that are informative about developing cognitive impairment in the future.

The dataset at its inception in the first year contained 1,213 subjects who were studied using a large range of features. It is worth noting that the data set contains two types of observations: variables and time series. The former refers to variables that are measured only once, typically in the first year’s visit (e.g. APOE genotyping or educational attainment) and the latter are variables with a time index attached (time series) measured at every subject’s visit (e.g. results of cognitive performance tests, hippocampal volumetry, subjective memory complaints etc.). Importantly, the dimensionality of the dataset grows every year (new measurements of features are added) as it does the number of samples as the subjects keep coming for their yearly visits, however the number of samples collected per year decreases (some volunteers drop the study). Table 1 shows the features types collected in The Vallecas Project. Each item in the table refers to a cluster of features containing related features. For example, the weight and height of the participants are included under the term Anthropometric. Note that some features are collected only during the first year for example APOE (*Genetics*) or the education level (*Demographics*), but for the most part, features are assessed at each year’s visit.

**Table 1:** The table depicts a classification of the features of the *Vallecas* dataset. Each item in the table refers to a cluster of features containing related features.

|  |  |  |
| --- | --- | --- |
| MRI | Genetics(APOE) | Demographics |
| Anthropometric | Sleep | Quality of life |
| Cognitive performance | Subjective Cognitive Decline | Diet |
| Physical exercise | Diagnoses | Neuropsychiatric |
| Traumatic brain injury | Engagement external world |  |

Figure 1 depicts the number of volunteers for the seven years life of the project. The total number of visits collected so far is 5203. Note that the data collection for year 6 and year 7 are still ongoing.

**Figure 1:**
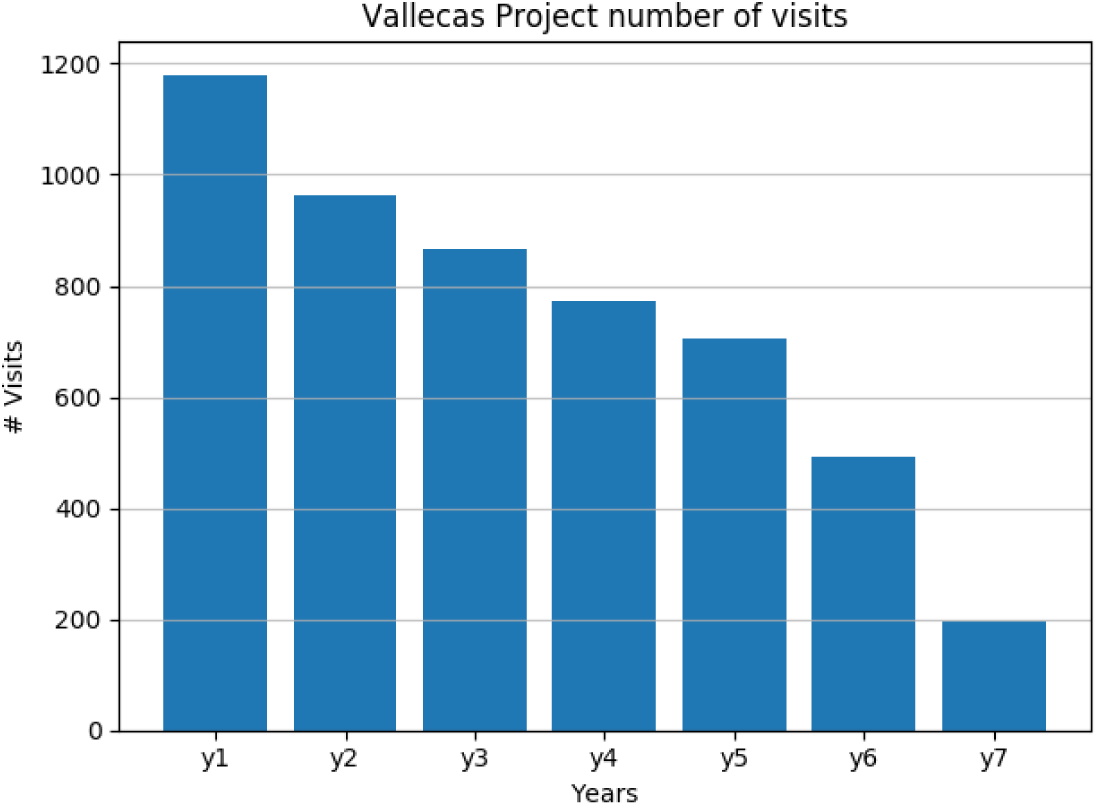
Number of volunteers across seven years of *Proyecto Vallecas*. As expected, the number of subjects decreases across time. The number of subjects in the year 1 was 1213 of which 33 were removed because were diagnosed with AD resulting in 1180 subjects. 965 of the initial 1180 came for a second visit (18% drop rate), 865 in year 3 (27% drop rate from year 1), 770 in year 4 (35% drop rate from year 1), 704 in year 5 (40% drop rate from year 1), 491 in year 6(58% drop rate from year 1) and 195 in year 7 which at this moment still ongoing. The total number of visits in seven years amounts to 5203.

## 2 Exploratory Data Analysis of cross sectional variables

In this section we perform Exploratory Data Analysis (EDA) for cross-sectional or short variables, that is, variables that are measured only once in the first year of the project. The longitudinal variables, one measurement for each project’s visit, are studied in the Section 3.6.

The objective of EDA is to build graphical representations that help us make sense of the data. EDA gives us a bird’s eye of the dataset aiding at identifying patterns and trends and most importantly, assisting in framing the type of questions we want to ask. EDA relies upon a careful pre-processing of the dataset. We perform EDA in a succession of steps. We will start with a preliminary data analysis plotting basic information of the dataset such as the age distribution and other demographics of the participants. The distribution of the most significant features of all type of features indicated in Table 1 is also studied. Furthermore, we plot the distribution of cognitive decline conditional to other features. This will set the stage for the correlation analysis described in Section 6 for both static (one observation) and dynamic variables (time series).

### 2.1 Genetic, Anthropometric and Demographic

The major risk of developing AD is age. Figure 2 shows the age of the participants at the inception of the project and the age of the participants that came to their sixth visit.

**Figure 2:**
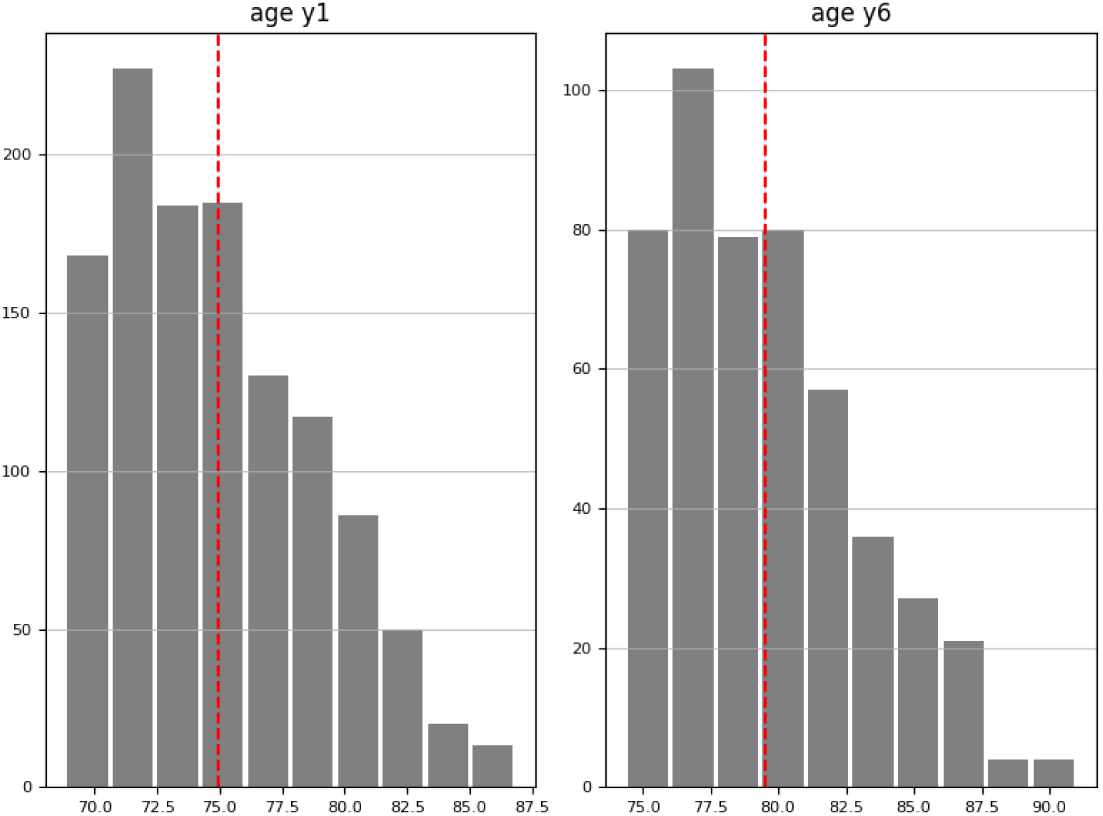
Histogram of the age of the participants in the first and last years of the *Proyecto Vallecas* dataset. The youngest and the oldest participant in year 1 were 69 and 86 years old respectively. In year 6, the minimum and maximum ages were 76 and 89 respectively. The discontinuous red line indicates the mean of the distribution.

Human apolipoprotein *ϵ* (APO*ϵ*) is a cholesterol carrier that supports lipid transport protein coded by the polymorphic APOE gene. The APOE gene has three major alleles: *ϵ*2, *ϵ*3 and *ϵ*4 [Mahley and Rall Jr, 2000], [Hauser et al., 2011]. Carriers of one copy of the *ϵ*4 allele (heterozygotes) have an increased risk of 3-4 times of developing sporadic AD, and carriers of two copies of the *ϵ*4 allele (homozygotes) have an increased risk of 8-12 times of developing the disease [Heffernan et al., 2016].

The APOE *ϵ*4 variant is the largest known genetic risk factor for late-onset sporadic Alzheimer’s disease (AD). The risk of developing the disease increases 8-12 times, furthermore APOE *ϵ*4 carriers tend to progress to dementia at younger age. This risk, however, varies by demographic factors such as sex and ethnicity [Sepehrnia et al., 1989], [Mehta et al., 2007], [Helzner et al., 2008].

The estimated worldwide human allele frequencies of the APOE*ϵ*4 allele in the normal population is between 10 *−* 15%[Myers et al., 1996] (13.7% for “Caucasian”^2^ populations [Corder et al., 1994]). On the other hand, 40–65% of subjects diagnosed with AD have one or two copies of the *APOE£*4 allele [Saunders et al., 1993].

Figure 3 shows the APOE Genotyping in *The Vallecas Project*, classifying the subjects in three groups depending on whether they lack a copy of the APOE*ϵ*4 allele, have one copy of APOE*ϵ*4 (heterozygotes) or have two copies of APOE*ϵ*4 (homozygotes) [Farrer et al., 1997]. In this cohort, 82% were APOE*ϵ*4 negative, 17% APOE*ϵ*4 heterozygotes and 1% APOE*ϵ*4 homozygotes.

**Figure 3:**
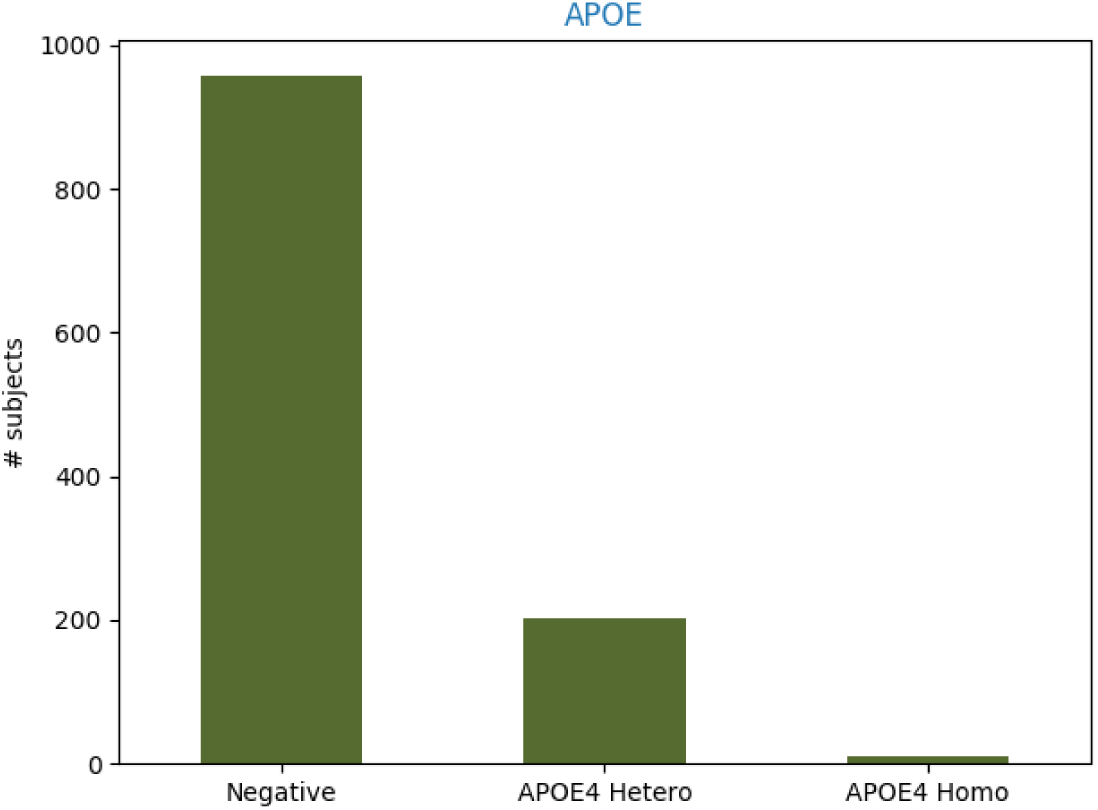
APOE Genotyping testing in *Proyecto Vallecas* dataset. 17% of subjects carry at least one copy of the allele *ϵ*4, slightly higher than the worldwide estimates in Caucasian population (13.7%) [Farrer et al., 1997]

People with a higher body mass index (BMI) in midlife are more likely to develop dementia. This conclusion was achieved in a study that pooled BMI data from a total of 1.3 million subjects in a period of up to 38 years from the original measurement of their BMI [Kivimäki et al., 2018]. The relationship between BMI and developing dementia is however complicated, the study found that people who developed dementia tend to have a lower than average BMI in the years leading up to the diagnosis. Nevertheless, the causal link between BMI and dementia is debatable, the reported relationship between BMI and dementia could be a product of the early stages of the disease rather than a risk factor.

Any causal statement relies upon the occurrence of events in time and since the onset of dementia is unknown, we must be very cautious about establishing causal links between, in this case, BMI and the diagnosis of AD. We must thus keep in mind the caveat that while higher BMI in midlife (our sample is composed of elder subjects) could be a risk factor for dementia, preclinical (undetected) stages of the disease could also produce weight loss and mask this effect.

According to the BMI metrics (25.0—29.9 overweight, *>* 30.0 obese) our population is overweight (right tail is longer than left tail in the top left histogram). Obesity (Higher midlife BMI) is related to higher risk of dementia and AD, independently of obesity-related risk factors and co-morbidities [Tolppanen et al., 2014], [Nepal et al., 2014]. However, a recent meta-analysis [Winter et al., 2014] found not association between overweight and an increased risk of mortality, on the other hand, the study found an increased risk for those at the lower end (BMI *<* 23.0). If we pay attention to these results, weight loss is a useful feature to monitor in older age.

Figure 4 shows the distribution of anthropometric variables: BMI, abdominal perimeter, weight and height, measured in the first visit. The mean and standard deviation of the BMI in our population is *µ_BMI_ =* 27.3 and *σ_BMI_ =* 3.6. 26.8% of the population have healthy weight (*BMI ∈* [18.5, 25]), 51.8% have over weight (*BMI ∈* [25, 30]) and 21.4% are obese (*BMI >* 30). The skewness, a measure of the asymmetry of the probability distribution, is positive in BMI (0.37), weight (0.49) and height (0.29).

**Figure 4:**
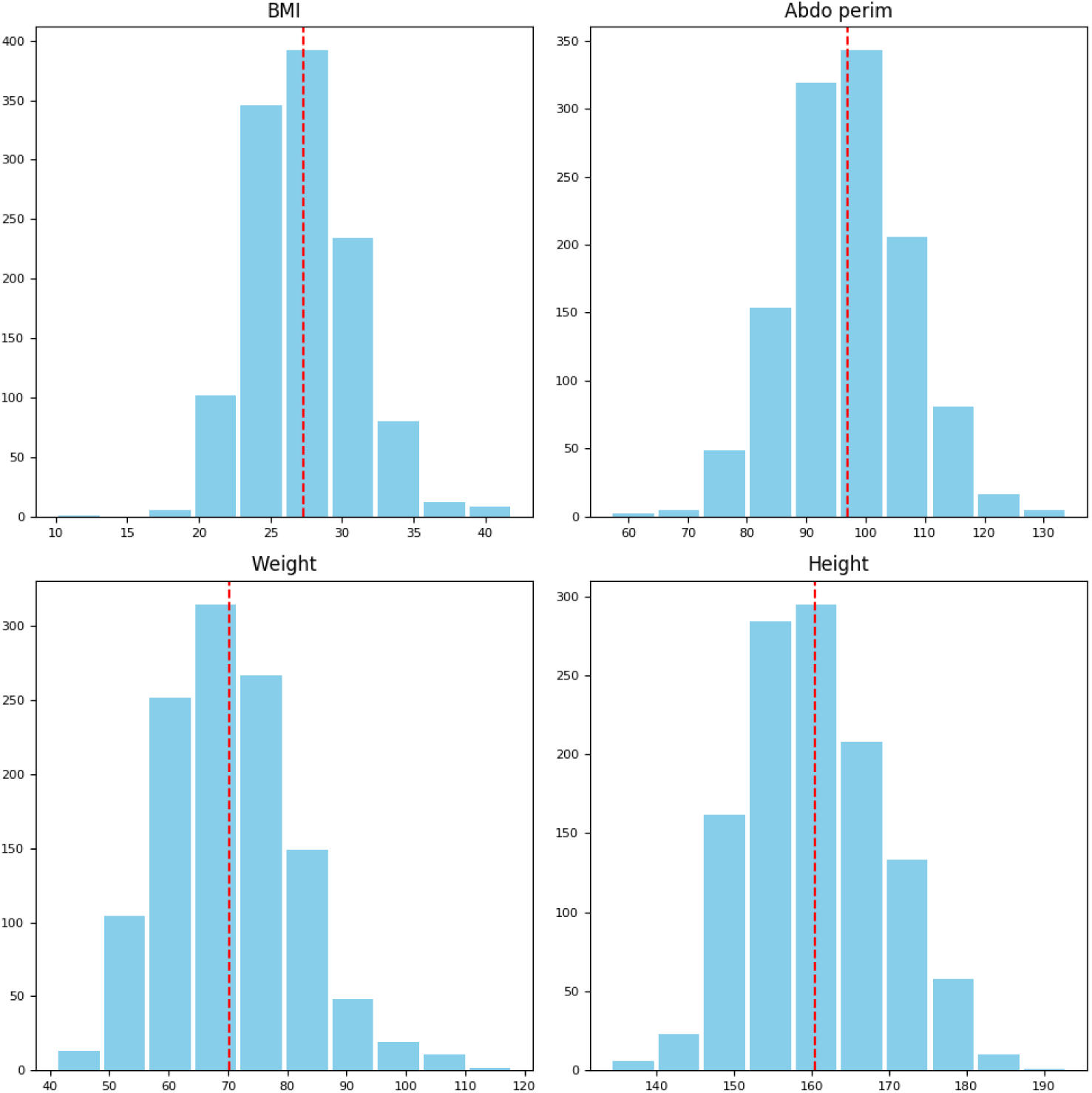
Histogram of anthropometric variables measured in year 1 of *Proyecto Vallecas* dataset. Clockwise, Body Mass Index (min=16.95, max=41.93), abdominal perimeter (min=57cm, max=134cm), weight (min=41kg, max=118kg) and height (min=134cm, max=193cm).

Although there are gender-specific risk factors for AD, a convincing account of how brain structure and function may vary by sex is missing [Mielke et al., 2014]. Epidemiological studies show that after age, the main risk factor for developing AD is sex [Vina and Lloret, 2010], [Mazure and Swendsen, 2016]. While it is well-known that the incidence of the disease is higher in women, it is far from clear that this can be entirely attributed to the higher longevity of women versus men. Socioeconomic factors such as the limited access to high education and the labor market of women born before the 1960s could play a role.

From a physiological perspective there are gender differences worth mentioning. Mitochondria of younger women are better protected against amyloid-beta toxicity, but this advantage lessens with age. Estrogenic therapies to deal with mitochondrial damage in AD have failed so far. A 2014 cohort study reports that women with the APO*ϵ*4 allele are at greater risk of developing AD than are men with this allele [Altmann et al., 2014]. The same study argues that the inconsistency of previous findings about this allele could be motivated by researchers overlooking the APOE*ϵ*4-sex interaction.

Women and men seem to have different clinical presentations of the Alzheimer’s disease. According to [Sinforiani et al., 2010] men tend to show more aggressive behaviors and more co-morbidity, while women would present more affective symptoms and disability but longer survival times. These behavioral differences would indicate sex-based neuropathologies in need of clarification. A report from the MIRIAD dataset [Malone et al., 2013]-a database of volumetric MRI brain-scans of Alzheimer’s sufferers and healthy elderly people-suggest that hippocampal atrophy in subjects with probable AD has a faster rate in women as compared to men, making female sex a risk factor for faster descent into AD [Ardekani et al., 2016].

Figure 5 shows the histogram for sex in the *Vallecas* cohort.

**Figure 5:**
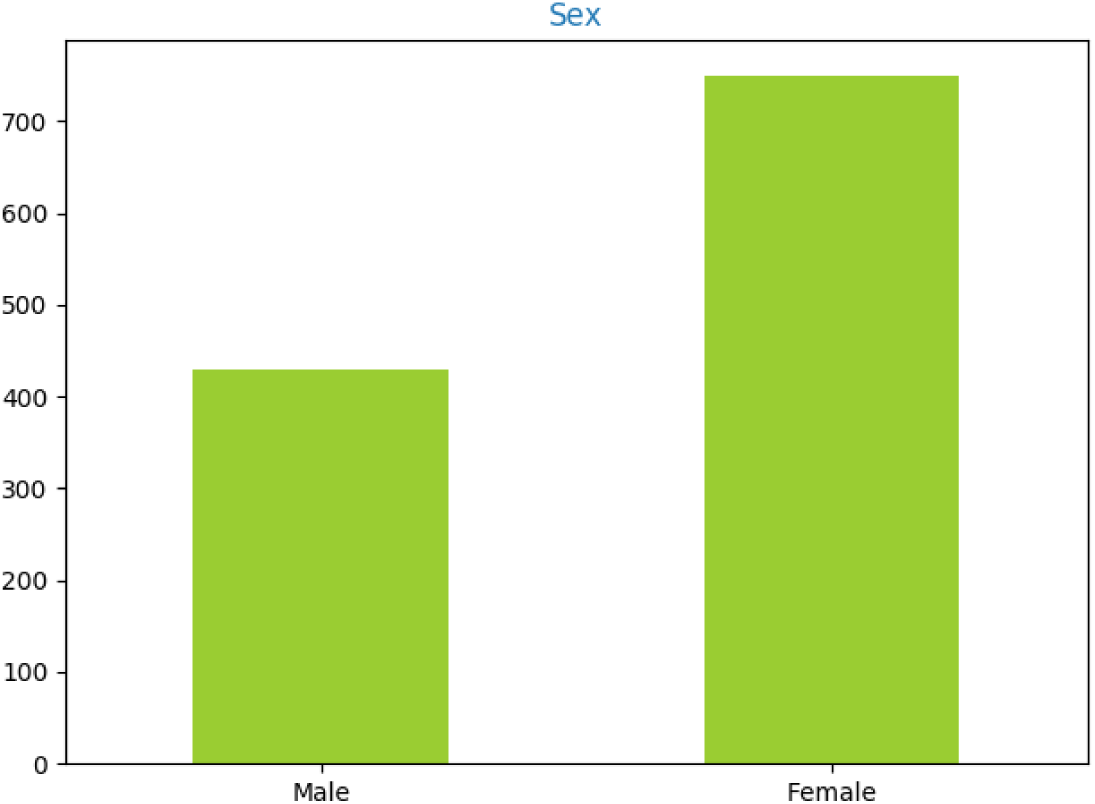
Histogram of sex distribution. There is majority of females, 753 versus 427 males.

A 1986 study reported that compared to right-handers, left-handers are less vulnerable to the cognitive problems associated with AD [De Leon et al., 1986]. However, hand laterality seems play a very marginal role, if any, in AD. In fact, research relating handedness preference and memory function have been abandoned since the early pre-MRI years.

Figure 6 shows the histogram for hand laterality in the *Vallecas* cohort.

**Figure 6:**
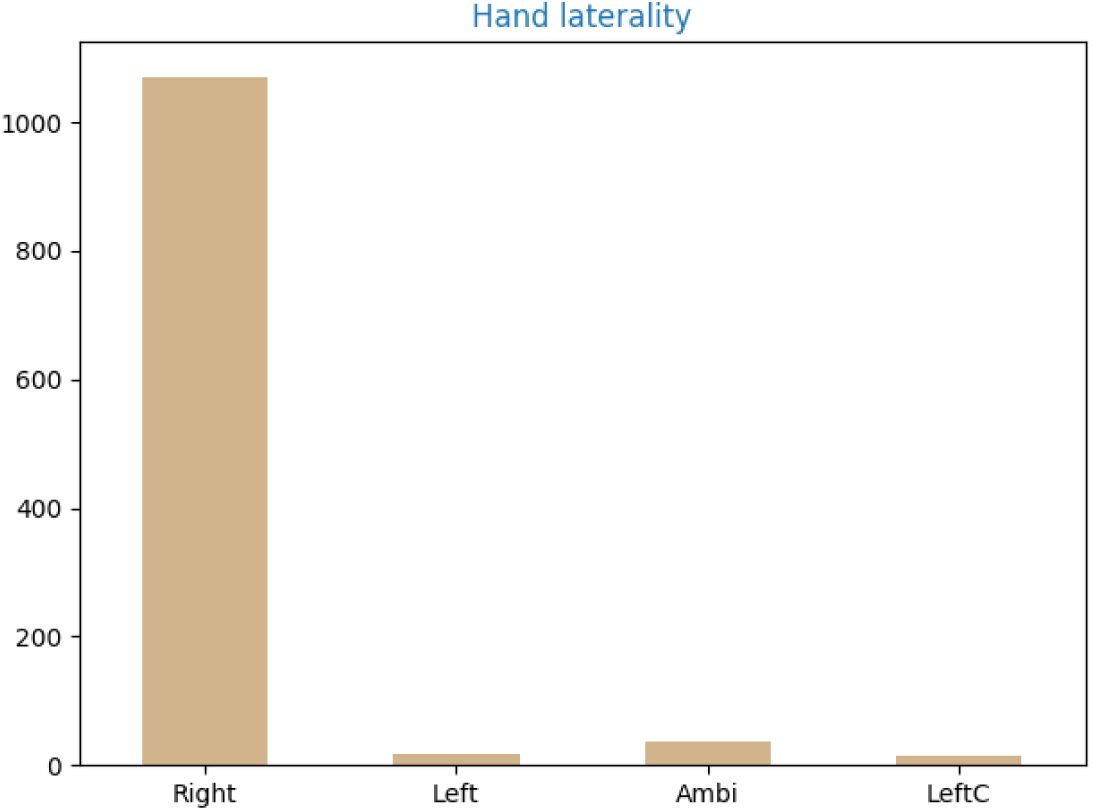
Histogram of hand laterality. There are 1106 right-hander (93.8%), 19 left-hander, 39 ambidextrous and 16 were forced right-hander about born left-hander (6.2% non right-hander).

A 2018 study using data on cognition from the 2000 and 2010 Health and Retirement Study [Crimmins et al., 2018], found that people with more education have lower prevalence of dementia and more years of cognitively healthy life. In a postmortem study of 86 brains, researchers found an association between number of children and neuropathology of AD for women only [Beeri et al., 2009]. Researchers argue that this difference may be due to sex-specific mechanisms (e.g. estrogen) rather socio-economic differences present in both men and women. However, in addition to the limitations of the study (women were older than men) the ways in which motherhood affect AD risk are more complex, even producing inconsistent results. A 3.5 thousands women study [Jang et al., 2018] found that women who give birth to five or more children may be more likely to develop Alzheimer’s disease than women who have fewer births, furthermore, incomplete pregnancy was associated with lower risk of AD in late life. But in a study with 14.5 thousands women presented at the 2018 Alzheimer’s Association International Conference in Chicago, researchers reported that women with three or more children had a 12 percent lower risk of dementia compared to women with one child [AAI, 2018].

Social isolation and loneliness is believed to be a risk factor in AD [Holmén et al., 1992], [Fratiglioni et al., 2000], [Shankar et al., 2013], [Holwerda et al., 2014], [Evans et al., 2018]. A 2007 longitudinal study with up to 4 years of in-home follow-up, reported a positive association between loneliness and an increased risk of late-life dementia [Wilson et al., 2007].

Figure 7 depicts demographic variables including: educational attainment, number of sons, years of work as an employee, self-assessed economic status, marital status and the number of people living at home.

**Figure 7:**
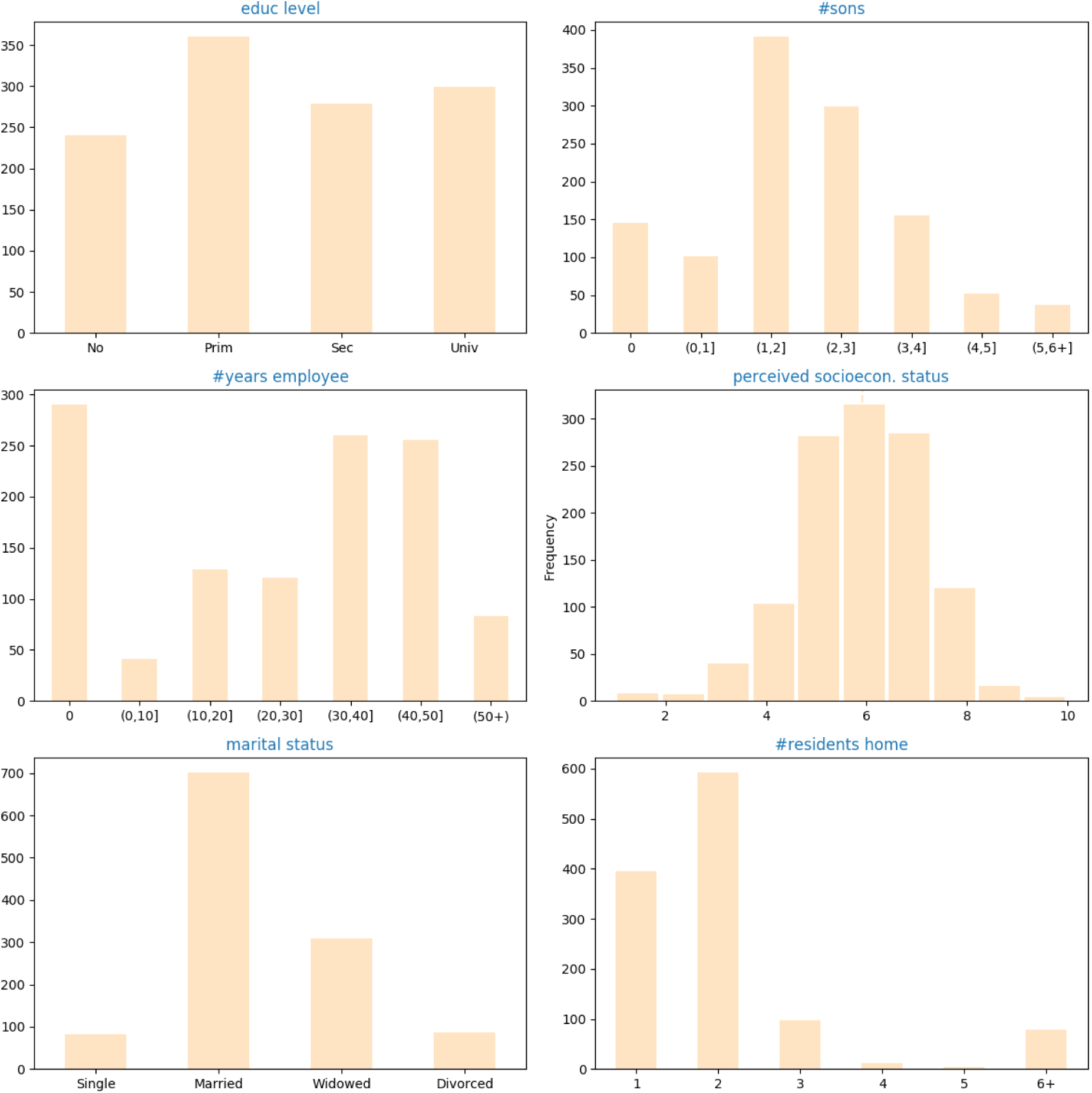
Histogram of demographic variables in the *Proyecto Vallecas* dataset measured in the first year. Clockwise, educational attainment classified as *no formal education, primary school, secondary school and university degree*, number of sons, number of years of work as an employee, self-assessed economic status (1 the lowest, 10 the highest, marital status (*single, married, widows, divorced*) and the number of people living at home.

We close the section of demographic variables with census income estimate based on the ZIP code. Lifestyle factors affect the susceptibility to developing AD and these factors are in many ways related to the level of personal income. *Ceteris paribus*, higher income subjects have better opportunities for education, rich and varied diet, leisure and health-care facilities than residents of lower income areas. The influence of the income level on AD mortality is, however, uncertain. Nevertheless, the variation of income-related modifiable lifestyle risk factors (diet, physical exercise, social life, job) across life span, makes it very difficult if not impossible to suggest a clear association between income level and risk of developing dementia [Stępkowski et al., 2015], [Ferri and Jacob, 2017].

Stress is a coping mechanism for taxing changes impinged upon us in our environment. In particular, allostasis is the adaptive mechanism put in place by the body in anticipation for departures from homeostatic equilibrium triggered by stressful situations [Ellis and Del Giudice, 2014]. However, not all the stressors are detrimental, stressors can also have beneficial effects. The duration and cyclicity of the stress is particularly important and chronic stress is unequivocally detrimental. The accumulative impact of neuroendocrine responses to chronic stressors might impair homeoastic mechanisms, setting the basis for depression, neurodegenerative disorders and other pathogenic conditions [Bisht et al., 2018]. Financially stressed individuals with recurrent difficulties to making ends meet, suffer from psychological stress that in turn may lead to cellular damage due to inflammation and oxidative damage [Hayashi, 2015].

Figure 8 shows the income distribution of the subjects estimated based on their home residency. The average annual net income was classified into 3 categories: (1) Low: up to 24,999€ (2) Medium: 25,000-49,999€ (3) 50,000€ and over. The GDP per capita in Madrid metropolitan area is 37,758€.

**Figure 8:**
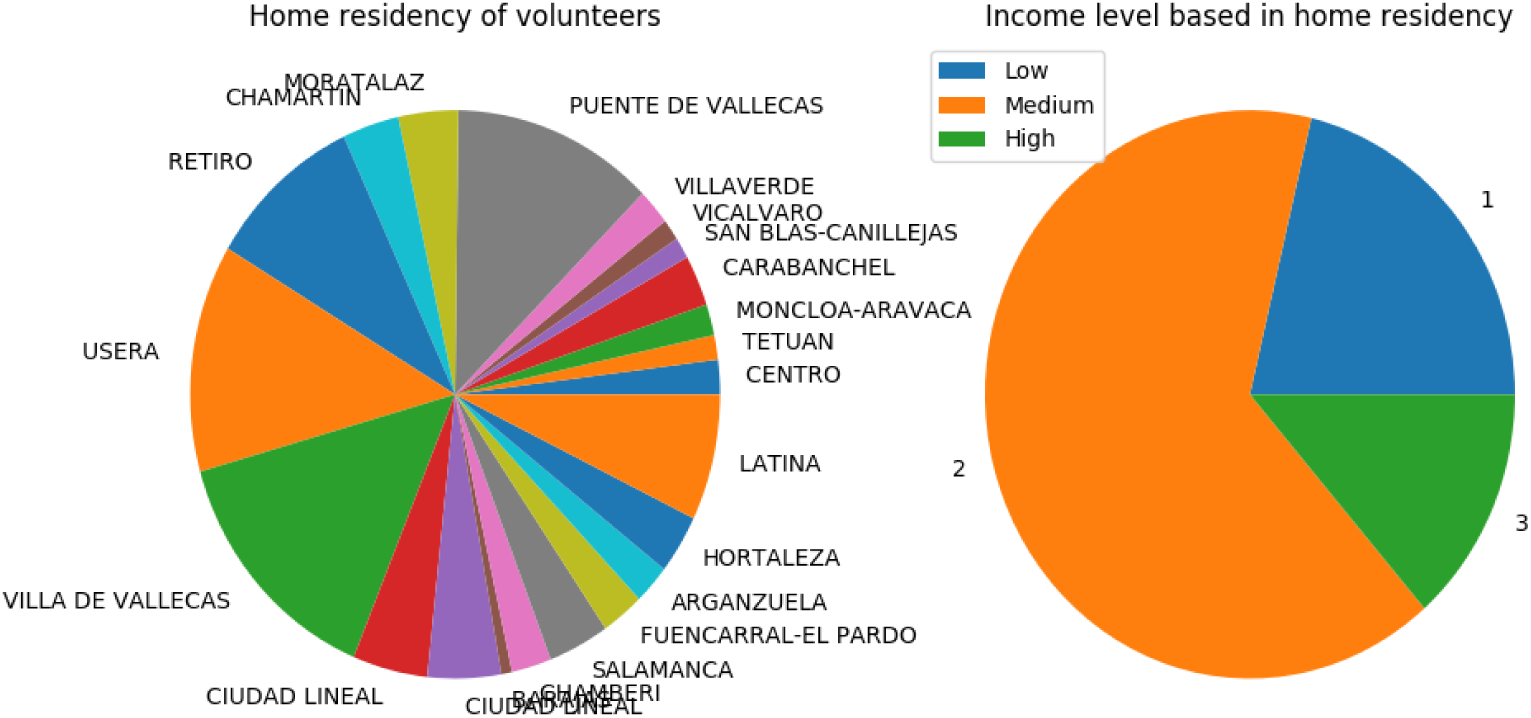
On the left, distribution of home residency by districts of *Proyecto Vallecas* volunteers. All districts of the city of Madrid are represented. Note that there are 156 subjects of a total of 1180 that are not residents in the city of Madrid and are not included here. On the right, we plot the income distribution as estimated based on the home residency, 251 subjects live in low income areas, 769 in medium income areas and 160 in high income neighborhoods of Madrid metropolitan area.

### 2.2 Sleep

Animal studies show that the concentration of Amyloid beta decreases drastically during sleep, suggesting that the removal of metabolic debris could take place during sleep [Kang et al., 2009], [Xie et al., 2013]. Poor sleep is linked to AD, in particular, the sleep–wake cycle has an effect on levels of amyloid-beta in the brain [Ju et al., 2014]. In a study that measured the association between amyloid-beta burden, relative to baseline, and sleep deprivation in 20 healthy controls [Shokri-Kojori et al., 2018], researchers reported that one night of sleep deprivation is sufficient for a significant increase in amyloid-beta in the right hippocampus and thalamus. This is in line with previous mouse studies that showed that brain levels of beta-amyloid decrease during sleep [Xie et al., 2013]. Tau protein is also affected in altered sleep cycles, a recent study [Holth et al., 2019] shows that excessive amounts of tau in the cerebral spinal fluid and spinal cord in extremely sleep-deprived adults. The increase in tau surpassed a 30% increase in the Amyloid beta peptide.

The two most important hallmarks of AD-amyloid plaques and tau aggregates-are thus, affected by sleep. However, the causal link sleep-AD is unclear; Does lack of sleep causes AD or is the pathology responsible for disrupting the sleeping cycle? This chicken-and-egg problem can not be solved with correlational neither with meta-analysis studies. Nevertheless, meta analysis [Bubu et al., 2016] have confirmed the association between sleep deprivation and cognitive impairment or AD, and can furthermore approximate an “average” magnitude of effect. More importantly, sleep-wake cycles can be approached as a modifiable risk factor of interest in the prevention of AD.

Figure 9 depicts features related to sleep patterns in the *Vallecas* cohort. Daily naps are less common than what it could be expected from a Spaniard population aged 70 or more; 37% of subjects do not take any nap at all, 24% of subjects sleep daily between 15 and 30 minutes and the 17% report taking naps of at least one hour a day. The majority of subjects report remembering their dreams 67.7% versus 32% that do not. The majority of subjects (60%) report to snore during sleep.

**Figure 9:**
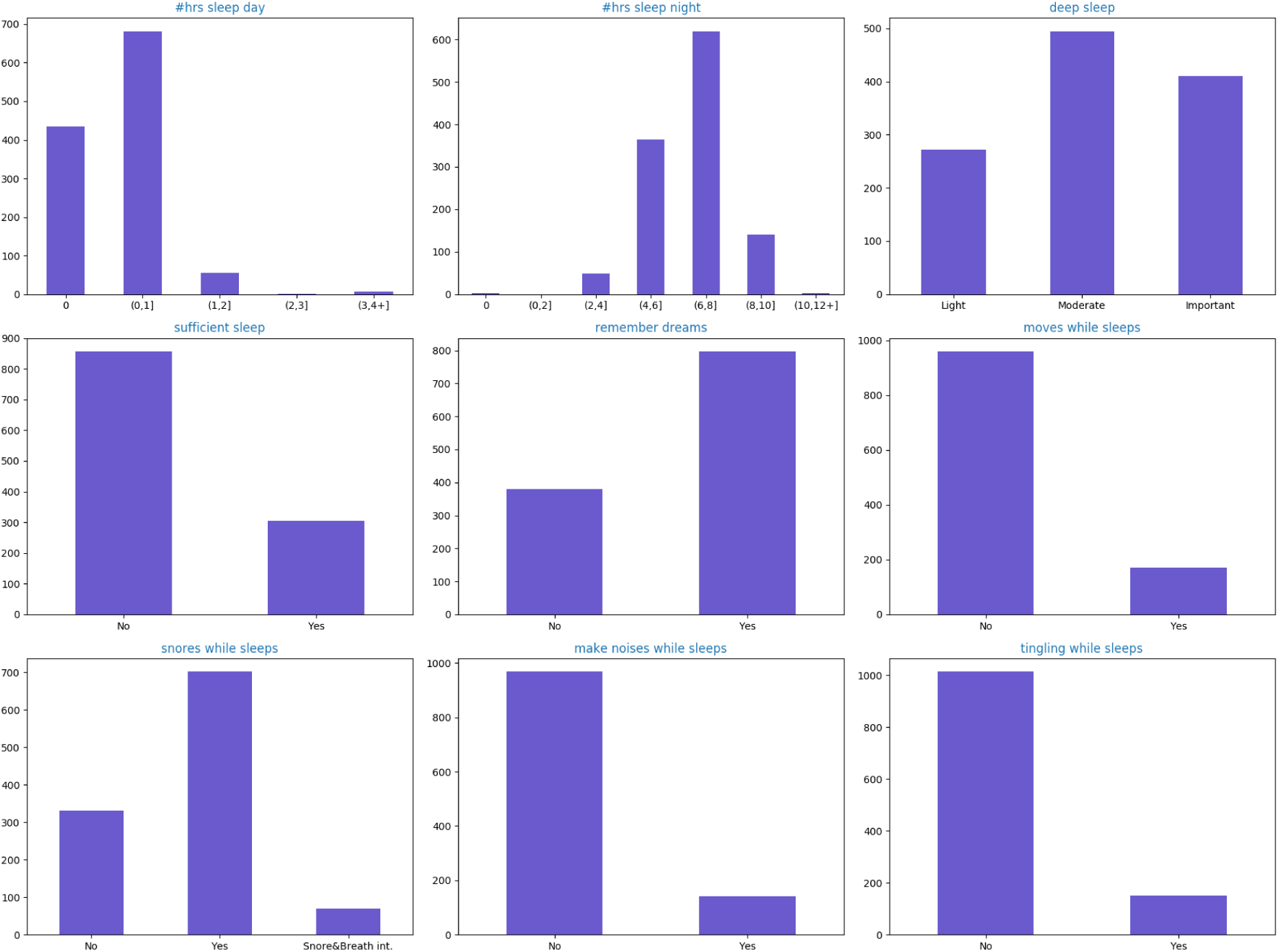
Histogram of sleep variables in the *Proyecto Vallecas* dataset in the first year. From left to right and up to the bottom, number of hours of sleep during the day (*µ =* 0.45, *σ =* 1), number of hours of night sleep (*µ =* 6.8, *σ =* 1.3), deep sleep (1 Light, 2 Moderate, 3 Deep), sufficient sleep (0 No, 1 Yes), remember dreams (0 No, 1 Yes), movements during sleep (0 No, 1 Yes), snoring (0 No, 1 Yes, 2 snore and difficult breathing), noises during sleep (0 No, 1 Yes), tingling during sleep (0 No, 1 Yes).

### 2.3 Life style: food and diet, physical exercise and social bonding

In a recent study performing PET imaging repeated in three years time, volunteers aged 30-60 with no symptoms of dementia when the study began were divided into two groups: 34 people ate a Mediterranean diet (high fruits, vegetables, lean protein) and 36 people ate a Western diet (high in saturated fats and refined sugar). Researchers reported that at the onset of the study, people that ate Western diet already had more beta-amyloid deposits than those who ate a Mediterranean diet. The follow-up scans showed an even greater difference in beta-amyloid deposits in the two groups, controlling for sex, age and genotype factors [Berti et al., 2018].

AD leads to metabolic impairment which is visible in reductions in cerebral glucose uptake. The early detection of decline in brain glucose metabolism could be of interest as a potential therapeutic intervention of AD. Since hypometabolism precedes clinical symptoms of AD, enhancing glucose uptake [Duran-Aniotz and Hetz, 2016], for example with ketone bodies [Gasior et al., 2006], could have a neuroprotective effect. The ketogenic diet (high in animal fat and low in carbohydrates) is a clinically established non-pharmacological treatment for epilepsy [Vining et al., 1998]. It is plausible that neuroprotective effects (enhanced cell energetics) observed in epilepsy could be of use in the treatment of neurodegenerative diseases, protecting against neuronal death through antioxidant and anti-inflammatory actions. From a biochemical standpoint, replacing glucose by ketone bodies can be seen as a sensible choice and there is research reporting an improvement in cognitive functioning in older adults with memory disorders followed by increases of plasma ketone body via the administration of medium chain triglycerides (MCTs) [Reger et al., 2004]. However, the potential downstream costs associated with the adherence to strict ketogenic nutritional recommendations are not fully understood.

Figure 10 depicts the type of food consumption as reported by the subjects in a questionnaire.

**Figure 10:**
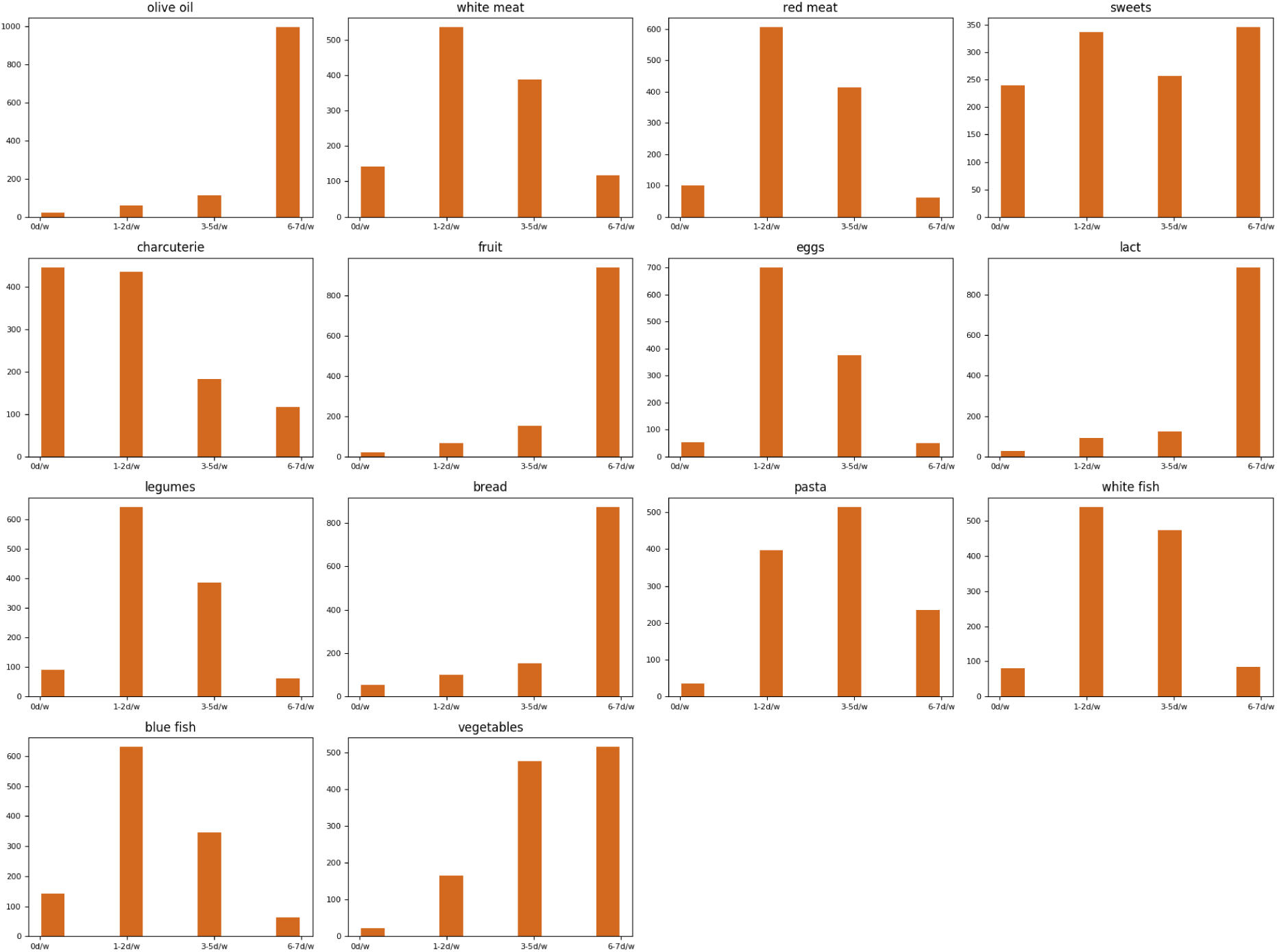
Histogram of type of weekly food consumption reported by the subjects in the first visit. The x-axis of each chart represents how many days a week they consume which type of food. The large majority of subjects report a daily consumption of bread, milk and olive oil. 84% of subjects consume olive oil 6/7 days a week, 74% bread, 44% vegetables and 29% report consuming sweets 6/7 days a week.

Figure 11 depicts the type diet based on the reported weekly food consumption. We distinguish between three different diets: diet with a strong glucemic component(carbs, sweets), diet with a strong proteic component or ketogenic (red meat, eggs, fish) and Mediterranean diet (fruit, fish, vegetables).

Studies have shown a beneficial effect of physical exercise in the aging brain, reporting increasing executive functioning and reduction in the expected with age decline of white and gray tissue density with increased fitness [Colcombe et al., 2003]. The effects of physical exercise in white matter of the aging brain are fascinating. Overall, white matter volume variations seems to be more contained than gray matter volume variation during healthy aging [Bartzokis et al., 2003]. However, white matter decline is better observed with DTI-based measures rather than conventional T1-weighted imaging [Giorgio et al., 2010].

**Figure 11:**
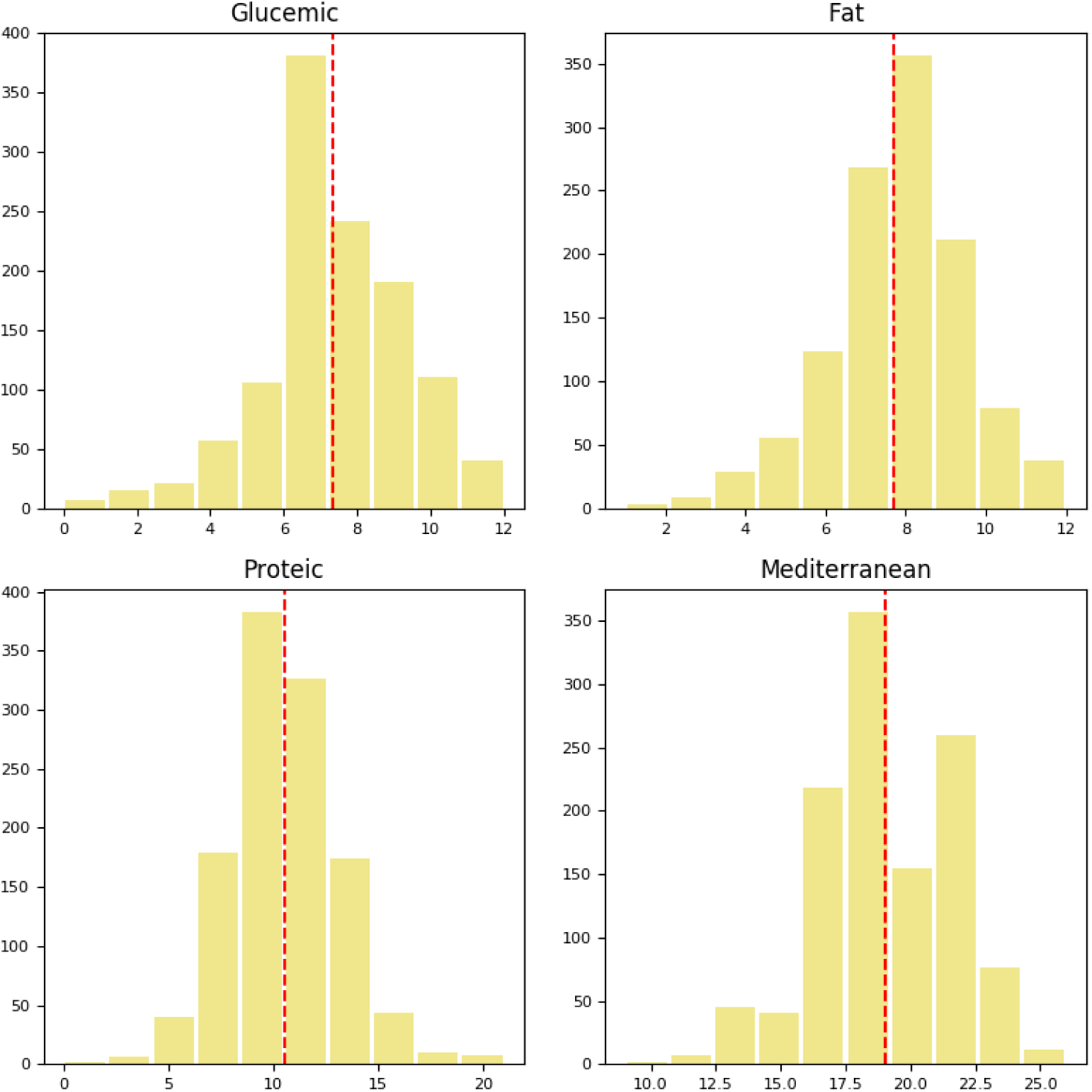
Histogram of type of diet based on weekly food consumption as reported by the subjects in the first visit. The x-axis of each chart represents the score for each type of diet, the larger the score the most prevalent the main component of the diet is in the subject’s diet. For example, a subject with a score of 12 in the glucemic diet will likely consume more sweets than a subject a score of 6, by the same token a subject with a large score in the Mediterranean diet will likely consume more fruit and vegetables than a subject with a lower score.

Figure 12 depicts features related to physical exercise. The subjects were asked the frequency in days per week of their physical exercise and the duration of the sessions.

**Figure 12:**
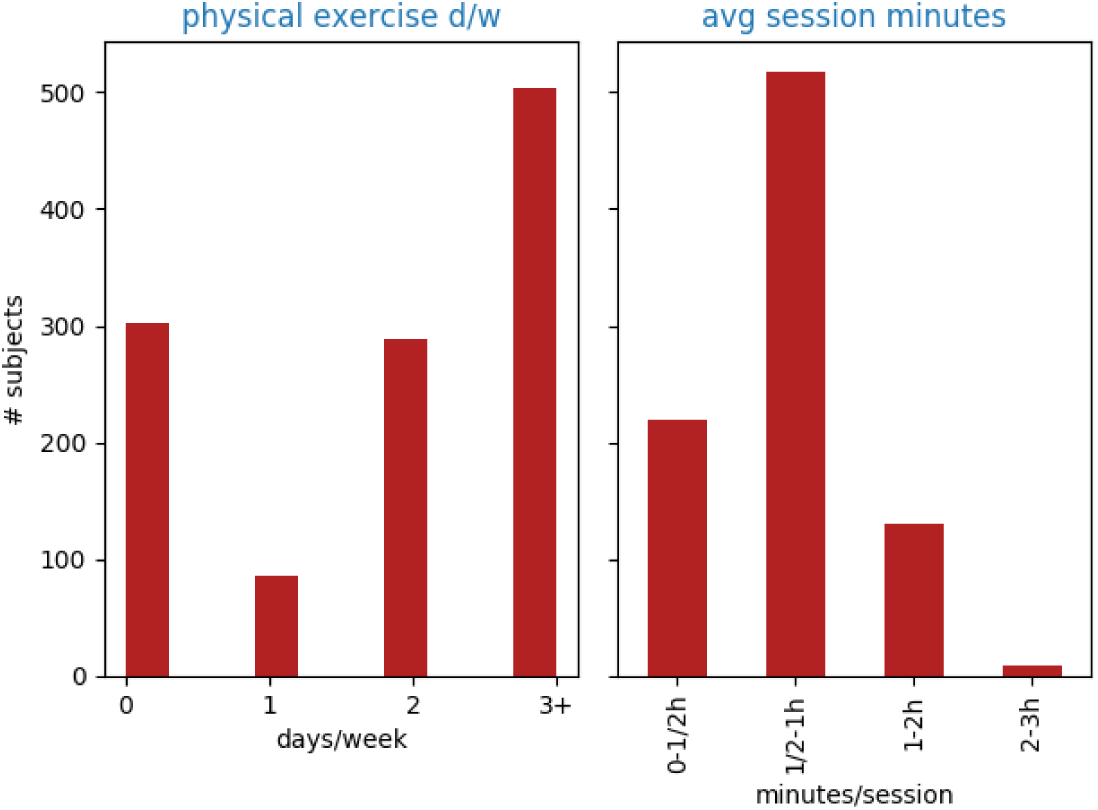
Histogram of features related to physical exercise. On the left days/week that subjects report to practice physical exercise (0,1,2,3 or more days a week), and on the right the average duration of the session (*less than 30 minutes, less than one hour, between one and two hours and more than two hours*)

Psycho social factors such as a rich social network, social engagement and mentally stimulating activities are commonly held as protective factors against dementia AD. Loneliness, depression, living alone are psycho social stressors that can increase the risk of developing AD [Johansson et al., 2013], [Sindi et al., 2015].

Figure 13 depicts features related to engagement with the external world. In particular, doing creative activities, going out with friends, travel and tourism, community activities (e.g. cultural associations and NGO), going to church, visit social club, go to the movie theater or art shows, go to sport events, how often listens to music, how often watches/listens TV/radio, reading habits (books, newspaper, magazines), and use of the Internet and information technologies. Of interest, church goers are a minority in our population, 75% never go to church, 17% that go a few times and 8% go often. 86% habitually watches the TV or listens to the radio, 68% read books or magazines often and only 29% use the Internet often.

**Figure 13:**
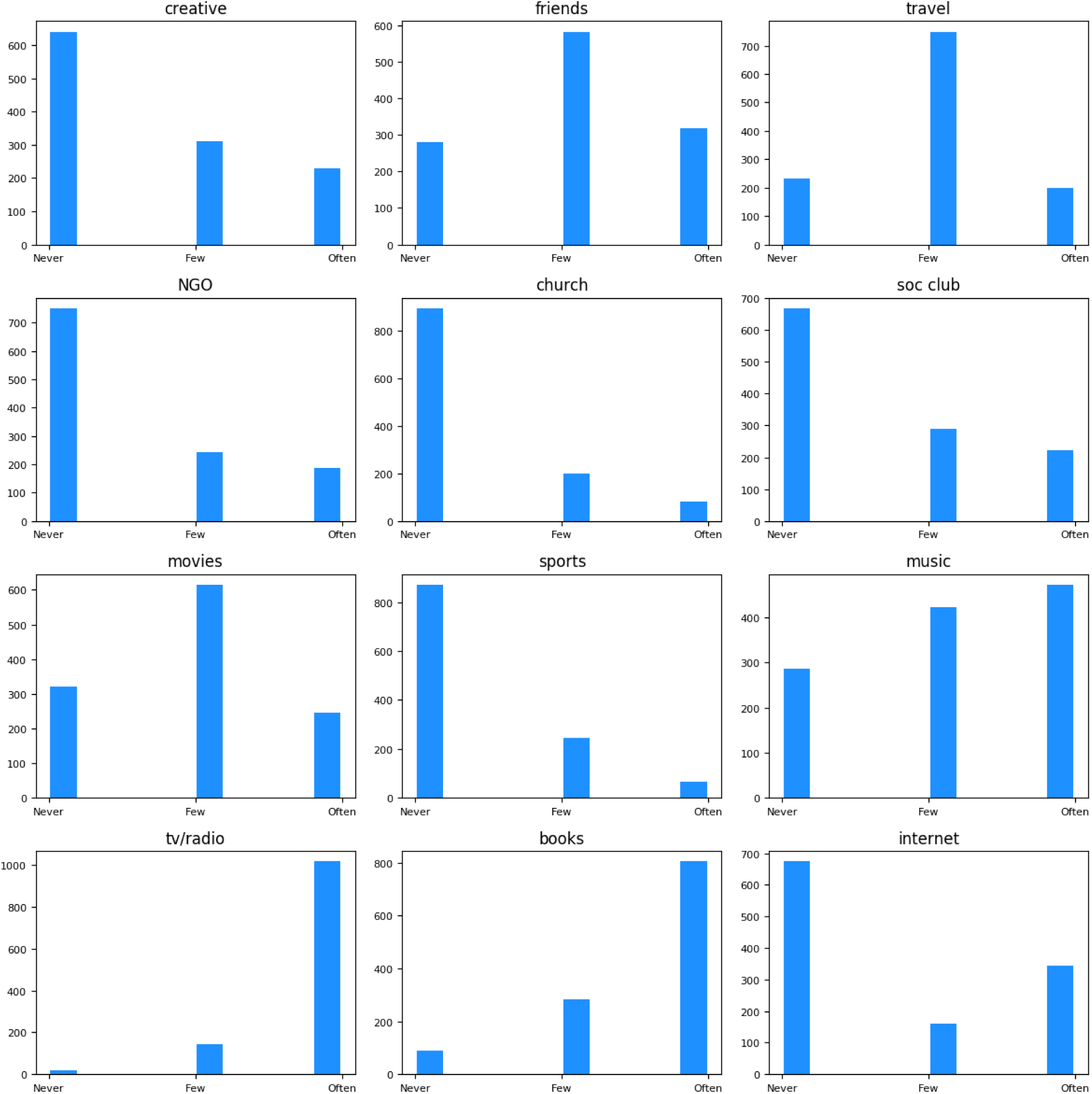
Histogram of features that reflect engagement with the external world from the part of the subject. From top down and left to right the charts depict the number of subjects that get involve in creative activities, going out with friends, travel and tourism, community activities (e.g. cultural associations and NGO), going to church, visit social club, go to the movie theater or art shows, go to sport events, how often listens to music, how often watches/listens TV/radio, how often reads (books, newspaper, magazines), how often uses the Internet.

### 2.4 Cardiovascular and cerebrovascular pathologies

A recent genome-wide study has been able to identify genetic variants that confer a risk of both cardiovascular disease and Alzheimer’s disease [Broce et al., 2019]. Vascular risk factors for AD such as diabetes, heart failure, stroke, hypertension and smoking among others could be involved in cerebrovascular dysfunction and AD pathology. Cardiovascular risk factors and their associated brain mechanisms are believed to play a role in the onset of AD. A better understanding of the interplay between cardiovascular disorders is not only beneficial from a clinical perspective but also from a prevention point of view, suggesting behavioral interventions (smoking cessation, healthy diet etc.) in early adult life [de Toledo Ferraz Alves et al., 2010].

Thyroid hormones have a significant impact on the cardiac system (excess thyroid hormone affects cardiovascular function [Klein and Danzi, 2007]). Stroke is a risk factor for coronary heart disease and subjects were asked if they had in the past hemorrhagic or ischemic cerebral ictus. Ischemic stroke and AD share pathophysiological mechanisms such as inflammation, immune exhaustion and neurovascular damage [Lucke-Wold et al., 2015]. The causality narrative between AD and hemorrhagic stroke seems to be reversed (AD causes X, rather than the usual search of factor X that causes AD), patients who had Alzheimer’s disease are at the highest risk of hemorrhagic stroke [Wang et al., 2014].

Figure 14 depicts features related to the cardiovascular health self reported in the *Vallecas* cohort. The subjects were asked whether they suffer hypertension, angina and heart attack, glucose metabolism disorders, dyslipidemia (an abnormal amount of lipids in the blood), arrhythmia and smoking habits. The majority of subjects report to have hypertension 53% vs 47%, this is in line with larger studies, for example in [Grau et al., 2011] 28,887 participants in Spain aged 35-74, high blood pressure was present in 47% of men and 39% of women, in [Lacruz et al., 2015]. 27% of the participants declared to be smokers and 33% ex-smokers.

**Figure 14:**
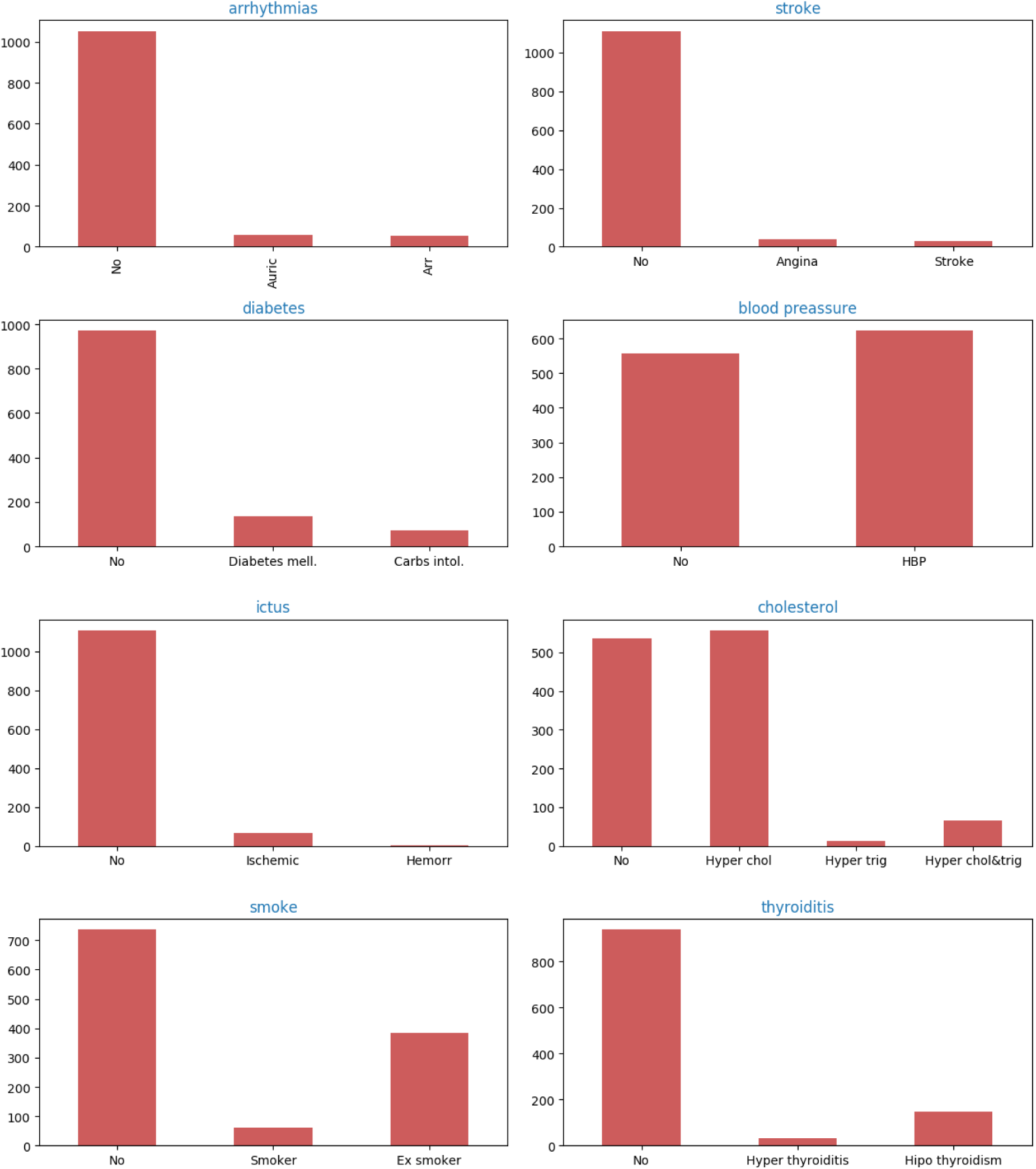
Histogram of features related to cardiovascular health. From top down and left to right: arrythmias (*No Arrhythmia, Atrial fibrillation, Arrhythmia*), heart stroke (*No past strokes, Angina, Stroke*), diabetes (*No diabetic, diabetes mellitus, intolerance to carbs*), hypertension, ictus history (*No, Ischemic, Hemorrhagic*), cholesterol (*No cholesterol, hyper cholesterol, Hyper triglycerides, both hyper cholesterol and triglycerides*), Smoke (*Non smoker, Smoker, Ex-smoker*), thyroid problems (*No thyroiditis, Hyper thyroiditis, hipo thyroiditis*).

Traumatic brain injury (TBI) is associated with long-term and acute disorders whose onset and ethiology are insufficiently understood. Violent head displacements in vulnerable brains is a precursor of dementia. A plausible mechanism of transmission is the propagation of abnormal proteins along damaged white matter pathways. Furthermore, TBI can initiate cerebrovascular pathology, which in turn could mediate in neurodegeneration including AD-like dementia [Ramos-Cejudo et al., 2018]. See [Mendez, 2017] for a summary of recent research on the TBI-dementia link.

Figure 15 shows the distribution of episodes (at least one) of traumatic brain injury in the *Vallecas* cohort.

## 3 Exploratory Data Analysis longitudinal variables

In this section we study the evolution of the variables measured in the *Vallecas* study. We dedicate the first section 3.1 to comment on the method employed to deal with data missing. We will study the progression of the longitudinal variables for those subjects that did not miss any visit of their 6 visits (complete-case-analysis). Thus, from an initial number of 1180 subjects that came to their first visit, we will study the time series for those subjects that came to all their visits from year 1 to year 6, which makes a total of 457 subjects spanned over a 6-year period. See Figure 16 for a quick representation of the procedure. The adjustment methods for missing data used here is complete-case analysis, that is, participants with missing data are excluded from the analysis. It ought to be noted that this is one methodological choice among other, see [Council et al., 2010] for a report with recommendations for missing data in clinical trials published by the National Research Council.

**Figure 15:**
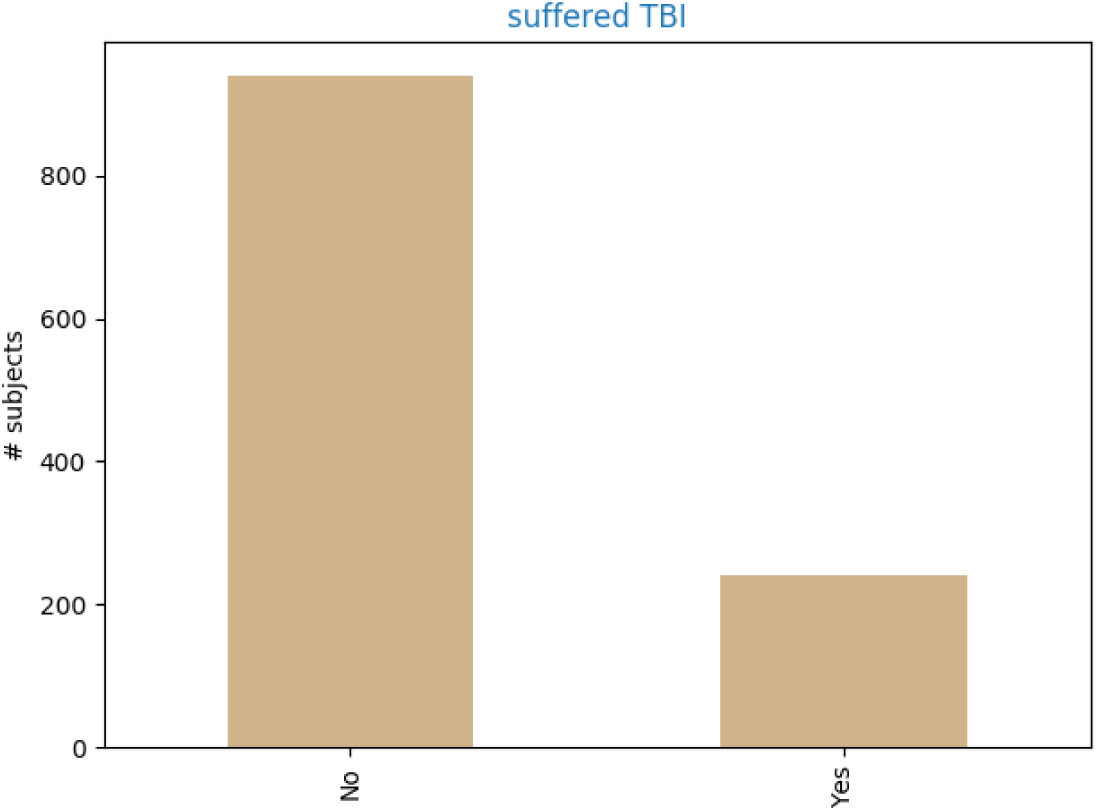
Histogram of traumatic brain injury. 20% of subjects declared to have suffered in the past an episode of traumatic brain injury of unspecified seriousness.

**Figure 16:**
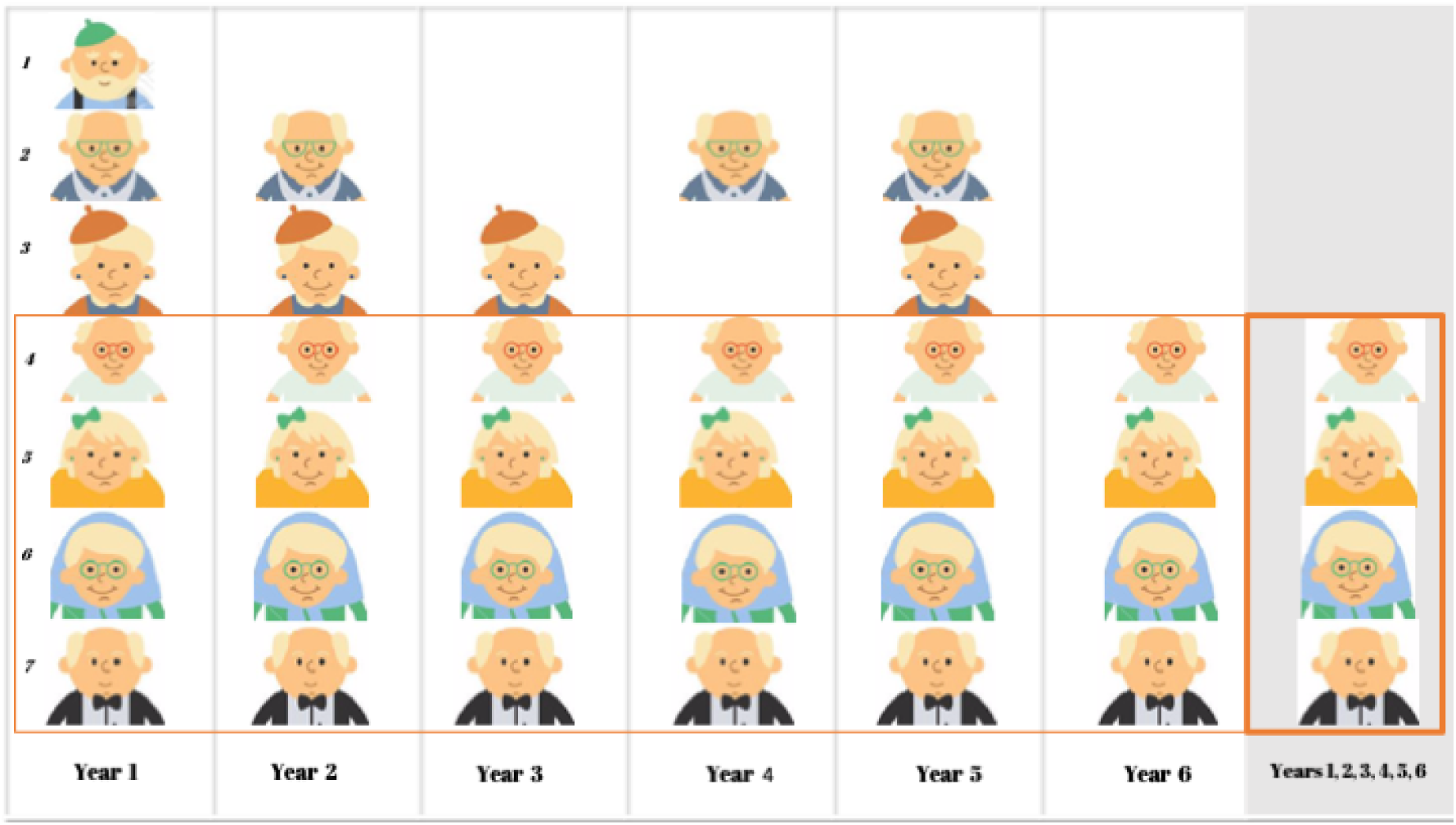
Sketch of the drop out handling in the *Vallecas* longitudinal study. Only the subjects that did not miss any of their six visits are considered for the longitudinal analysis described in this section.

The longitudinal variables are studied as time series of length 6 (number of yearly visits) with 457 observations for each time point. Neuropsychiatric features in section 3.3, quality of life in section 3.4, cognitive performance tests are shown in section 3.2 and the evolution of the diagnosis in Section 3.6 and MRI data including a description of the segmentation procedure is shown in section 3.7.

### 3.1 Treatment of missing imputation

Just like in any other longitudinal study, drop out is an important concern. The causes for missing visits can be related to first and foremost the voluntary and non remunerated design of the project and the age of the subjects, 70 at the inception. The discontinued participation can also be caused by the diagnosis of dementia, possibly in the clinical assessment performed in the study.

Missing data compromises statistical inference and represents a serious limitation for drawing conclusions [Little et al., 2012]. Although there are different methods for completion of missing data e.g. Maximum likelihood, Bayesian and multiple imputation [Hogan et al., 2004], it is worth reminding that the effectiveness of any statistical method relies upon the plausibility of its assumptions. Thus, it is desirable to have evidence of the mechanisms leading to missing data. There are two major possible scenarios: data missing are completely random and data missing are not at random. The first case is less stringent: the outcomes of participants that dropped out are not significantly different from those that did not drop out In the second case, data are missing not at random, outcomes are likely to be different from those of similar participants who did not drop out. In the latter case, the statistical inferential model needs to incorporate additional assumptions.

For our dataset, since the attrition rate is severe (1180 to 491 in year 6) we opt first to investigate whether the sample changes according to specific patterns, in which case we would not attempt to complete the missing data, and we would rather select the subset of subjects that did not miss any visit (471 subjects did not miss any visit from year 1 to year 6). Table 2 indicates that this is the case, *remainers* and the leavers are different (*p <* 0.05) for some variables.

**Table 2:** T-test for the means of two independent samples scores: group of subjects that were present at year 1 and year k (*k =* 2..6) versus subjects present that missed their *k-th* visit. The drop out of subjects in the study is not random because the groups are different for at least one variable (e.g. Education).

| Feature | Y1 – Y2<br>N=964,216 | Y1 – Y3<br>N=865,315 | Y1 – Y4<br>N=773,407 | Y1 – Y5<br>N=704,476 | Y1 – Y6<br>N=491,689 |
| --- | --- | --- | --- | --- | --- |
| APOE | > .05 | > .05 | > .05 | > .05 | > .05 |
| Familial AD | ** | ** | ** | ** | ** |
| Education | ** | ** | ** | ** | ** |
| Wealth | > .05 | * | * | * | > .05 |
| Internet | ** | ** | ** | ** | ** |
| Church | > .05 | > .05 | > .05 | > .05 | > .05 |
| Diagnosis (last) | > .05 | > .05 | > .05 | > .05 | * |
| Thyroid problems | > .05 | > .05 | > .05 | > .05 | > .05 |

Figure 16 depicts the drop-out handling the *Vallecas* longitudinal study. For example, to test the drop out in year 2 we compare the means between the two groups: (1) and (2, 3, 4, 5, 6, 7) because subject 1 drop the study in year 2, to test the drop out in year 4 referred to year 1 we compare between the groups: (1, 3) and (2, 4, 5, 6, 7) because subjects 1 and 3 leave the study in year 3. Thus, we compare *remainers* versus *leavers* for each year. In case the two groups are different for any year it would suggest that the data missing is not random.

Table 2 shows the statistical tests to study whether the subjects that drop out are significantly different from those that remain in the study. Note that we perform a two samples t-test comparing the subjects were present at year 1 and year k (*k =* 2..6) versus subjects present at year 1 but missed year k. The actual size of the groups is indicated in the second row, for example 689 missed their 6th visit and 491 came to their 6th visit. Since there are differences for at least one variable, it follows that the subjects drop out is not random and it is therefore advisable to pursue a complete-case longitudinal analysis.

To conclude this section, the complete-case analysis followed here to handle missing longitudinal data is justified on the basis of the non randomness of the missing data. As table 2 tries to convey, outcomes are likely to be different because the pattern of drop out of subjects is not random, that is, the hypothesis that those that remain and those that leave the study are the same is false in at least the variables shown in Table 2 (e.g. educational level, familial AD, internet use).

Note that we opt for complete-case analysis for longitudinal data, that is, an entire missing visits, but we used the much less stringent Multiple Imputation (MI) [Buuren and Groothuis-Oudshoorn, 2010] approach for dealing with missing data within a subject’s visit.

The complete-case analysis i.e. drop-out removal previously discussed will be used in Section 4 when we study the correlation analysis. The rest of the section deals with the evolution of the longitudinal variables.

### 3.2 Cognitive Performance

Figure 17 plots the evolution of four cognitive performance tests. Clockwise and starting on the top left chart: number of words that start by the letter “p” in one minute, number of animals recalled in one minute, score of speed processing of symbols and the score in the Mini-Mental Examination test (MMSE).

**Figure 17:**
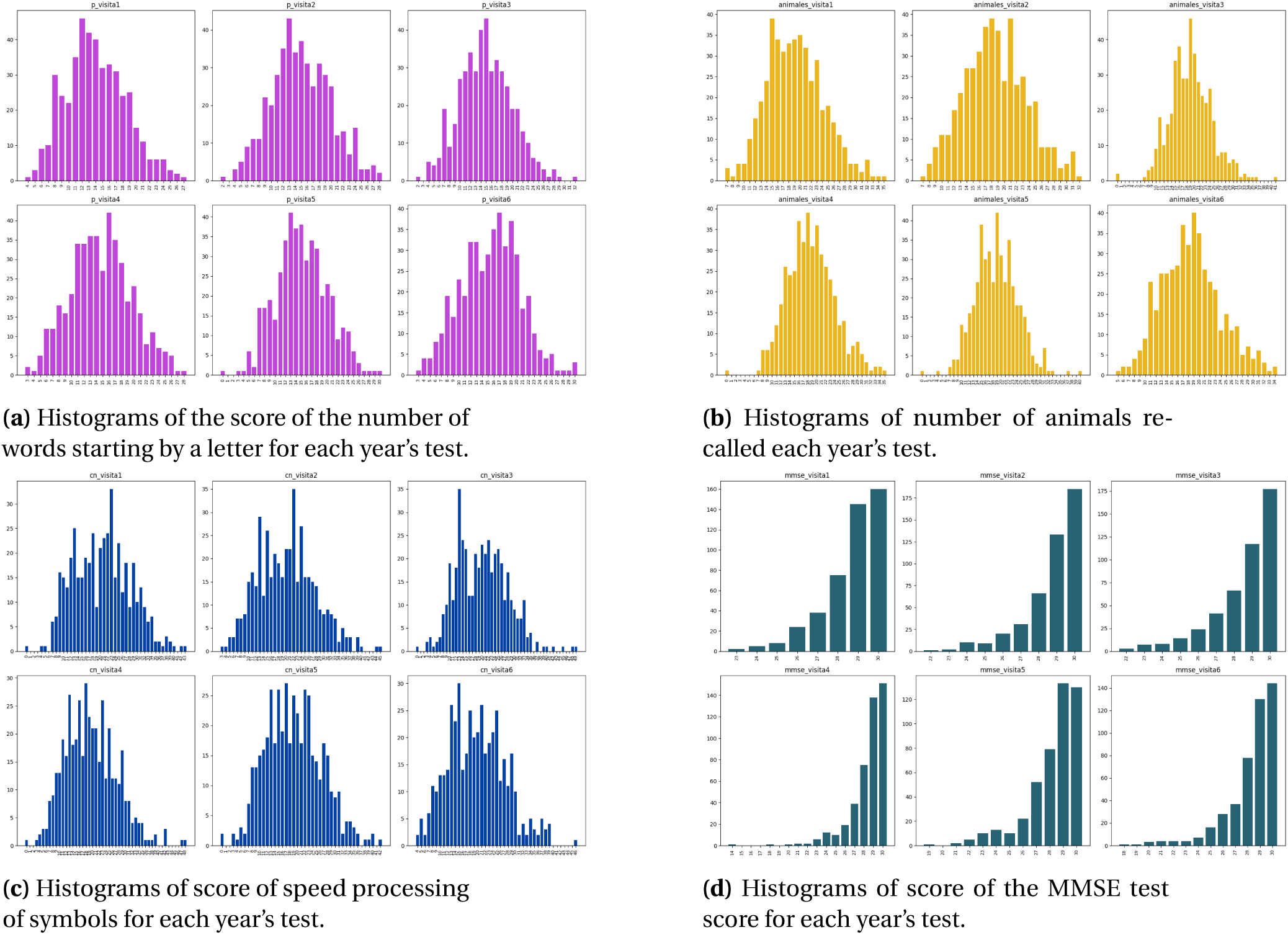
Histograms showing the evolution of features related to the cognitive performance of the subjects across the six-year visits (no missing visits). The scores from left to right and top to bottom refer to: number of words that start by letter p, number of animals, speed processing of symbols and MMSE.

Figure 18 plots the time series for four measures related to cognitive performance. Clockwise starting top left chart: score of speed processing of symbols, score for the number of animals that the subject can recall in one minute, score for the number of words that start by the letter p in one minute and the score in the mini-mental examination (MMSE).

**Figure 18:**
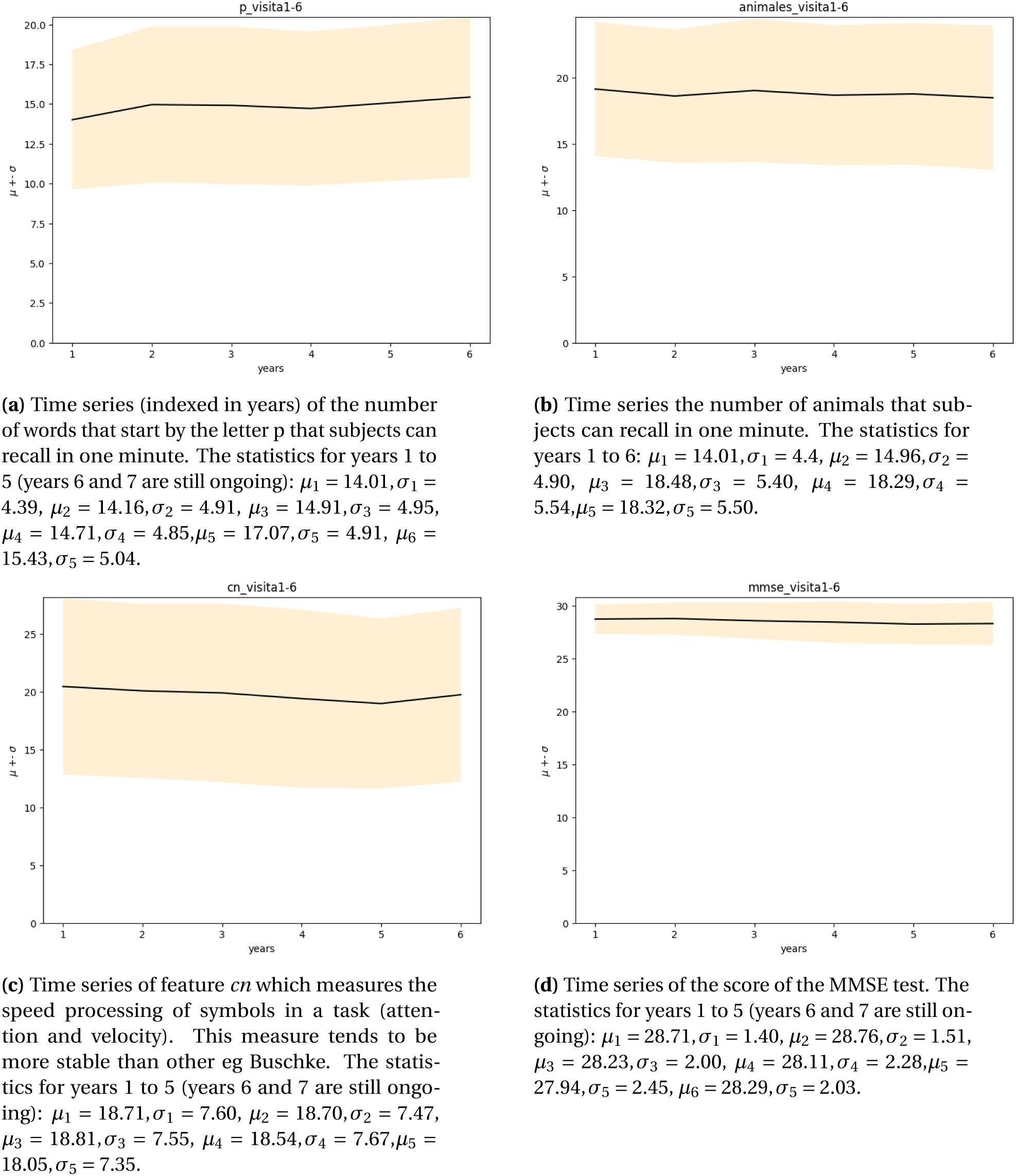
Time series of cognitive performance across the seven years of *Vallecas Project* of features related to cognitive performance. The x-axis are years from year 1 to year 6 and the y-axis the score. Each plot represents the mean (bold line) and the dispersion or standard deviation (shaded region) across the seven years of *Vallecas Project*.

Figure 19, shows the Buschke memory test which consists in making teach a subject a list of words to ask her later to remember of these words. This test is divided into 3 parts: main trial, interference test of attention and delayed test. Clockwise and starting in the top left chart: Free and Cued Selective Reminding Test (FCSRT) (a), (b) the arithmetic mean (three terms) of the free delayed recall test, (c) the total immediate recall and (d) the Buschke Integral score which is explained in a forthcoming paper. (We address the reader to the Appendix section for more details).

**Figure 19:**
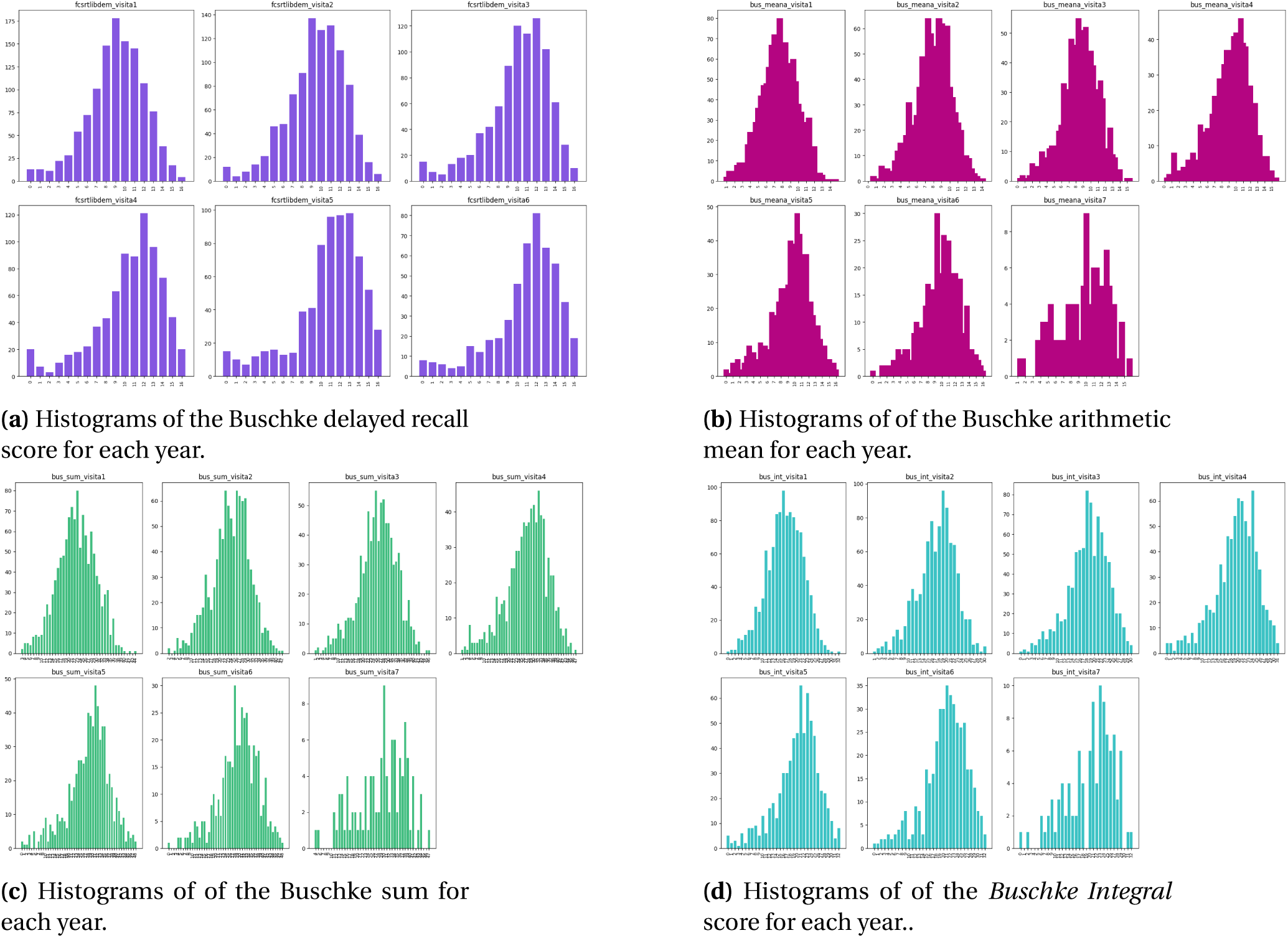
Histograms showing the evolution of features related to the cognitive performance of the subjects across the seven years of *Vallecas Project*. From left to right and top to bottom: Free and Cued Selective Reminding Test (FCSRT), the arithmetic mean (three terms), the sum and the *Buschke Integral*.

Figure 20 plots the time series for four measures related to cognitive performance measured using the Buschke test. Clockwise starting top left chart: delayed recalled score, mean, sum and the Buschke Integral score which is explained in a forthcoming paper. We address the reader to the Appendix section for more details).

**Figure 20:**
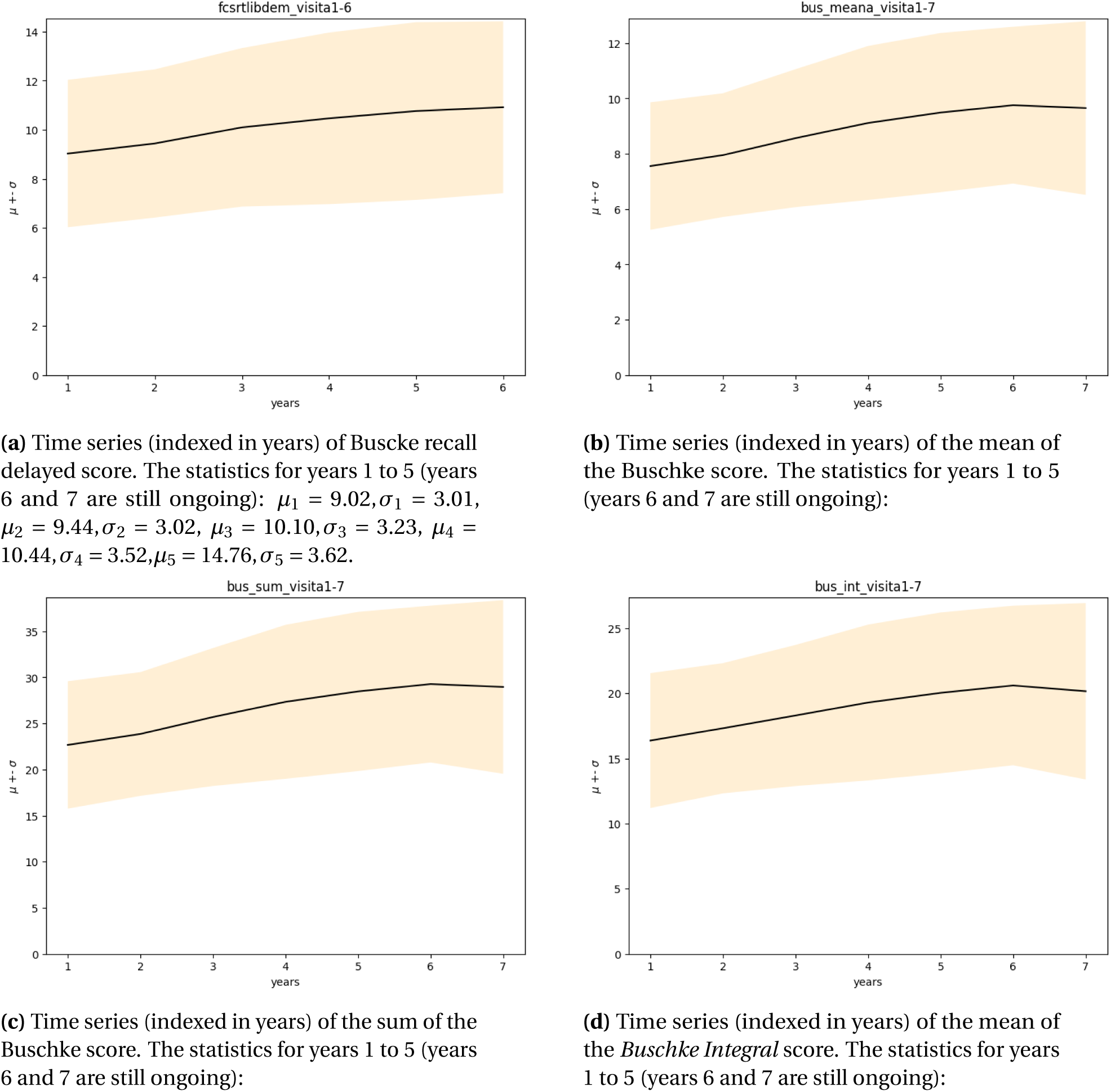
Time series of Buschke scores (delayed recall, mean of scores, sum of scores and the *Buschke Integral* score) across the seven years of *Vallecas Project* of features related to cognitive performance. The x-axis are years from year 1 to year 7 and the y-axis the score. Each plot represents the mean (bold line) and the dispersion or standard deviation (shaded region) across the seven years of *Vallecas Project*

### 3.3 Neuropsychiatric

Figure 21 plots the evolution of features related to neuropsychiatric disorders, specifically the Geriatric Depression Scale (GDS) [Yesavage, 1988] and the State and Trait Anxiety Scores (STAI) [Spielberger, 1985], [Julian, 2011].

**Figure 21:**
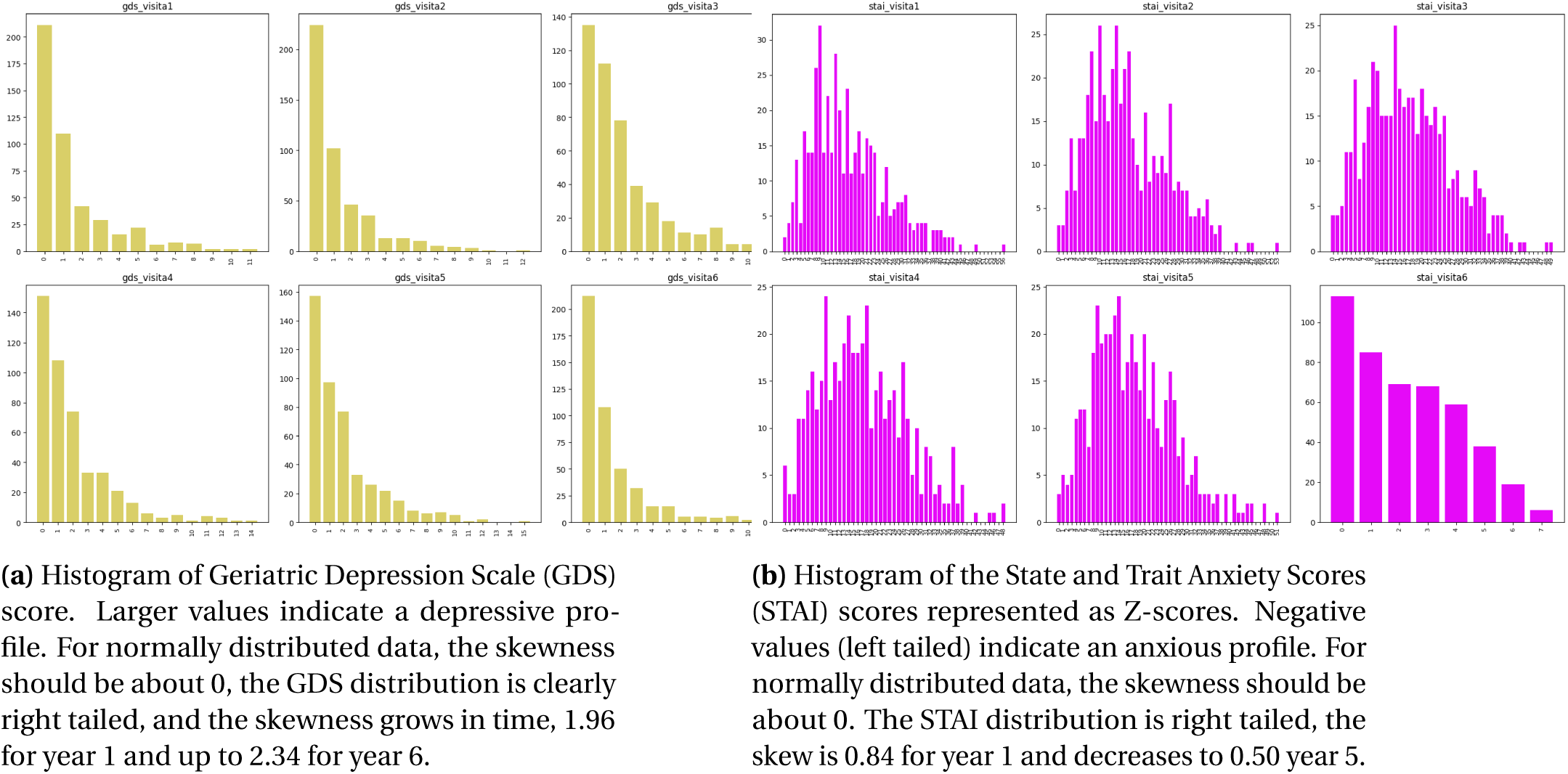
Histograms of Geriatric Depression Scale (GDS) and the State and Trait Anxiety Scores (STAI) z-score across the seven years of *Vallecas Project*.

### 3.4 Quality of Life self assessment

Figure 22 plots the evolution of features related to the quality of life, self-assessed by the participants in the *Vallecas Project*.

**Figure 22:**
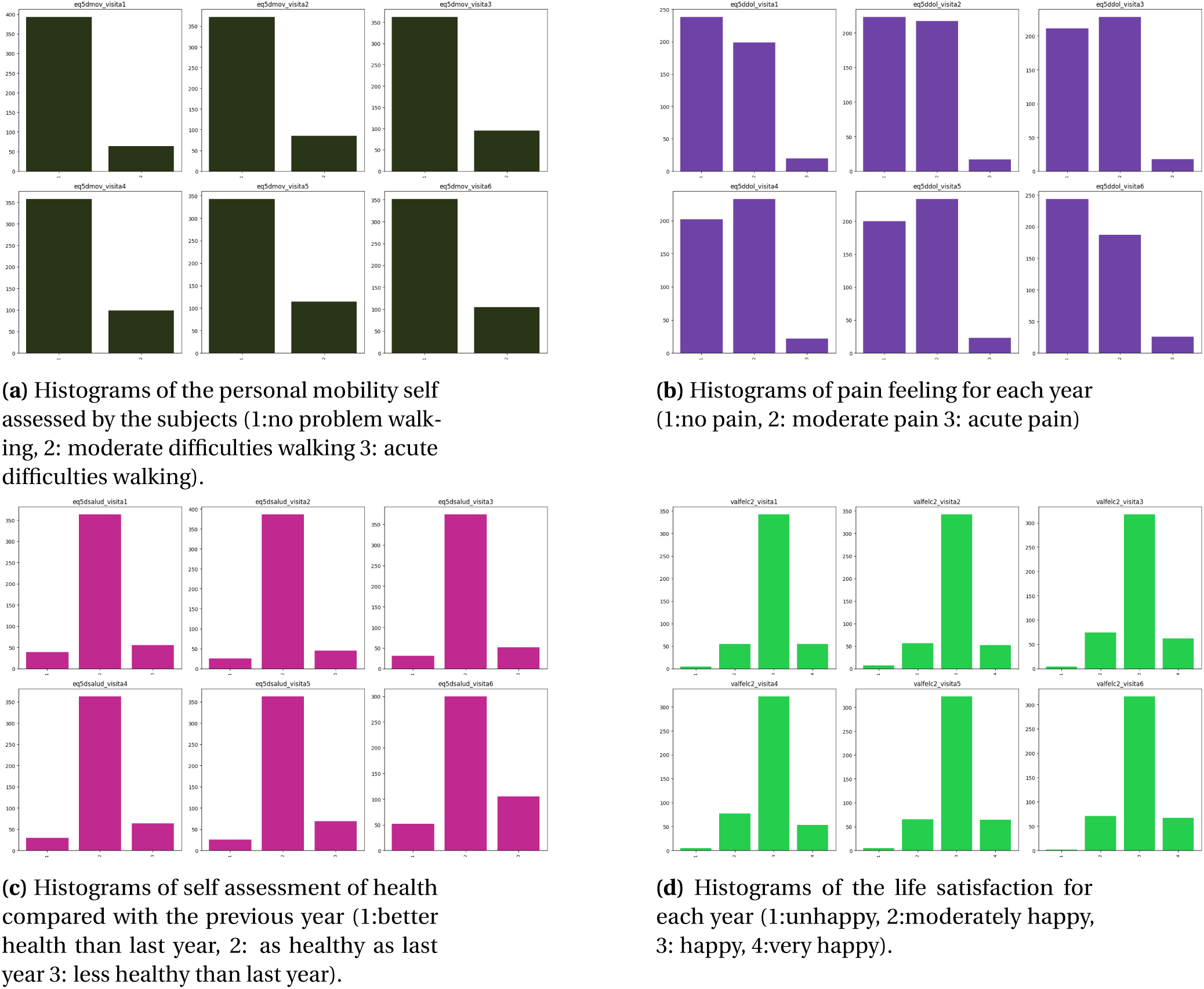
Histograms showing the evolution of features related to the quality of life of the subjects across the six years of *Vallecas Project* (no missing visits).

Figure 23 plots the time series for four measures related to quality of life self-assessed by the participants in *Vallecas Project*. Clockwise starting top left chart: score of personal mobility, feeling pain, self assessment of health compared with the previous year and life satisfaction score.

**Figure 23:**
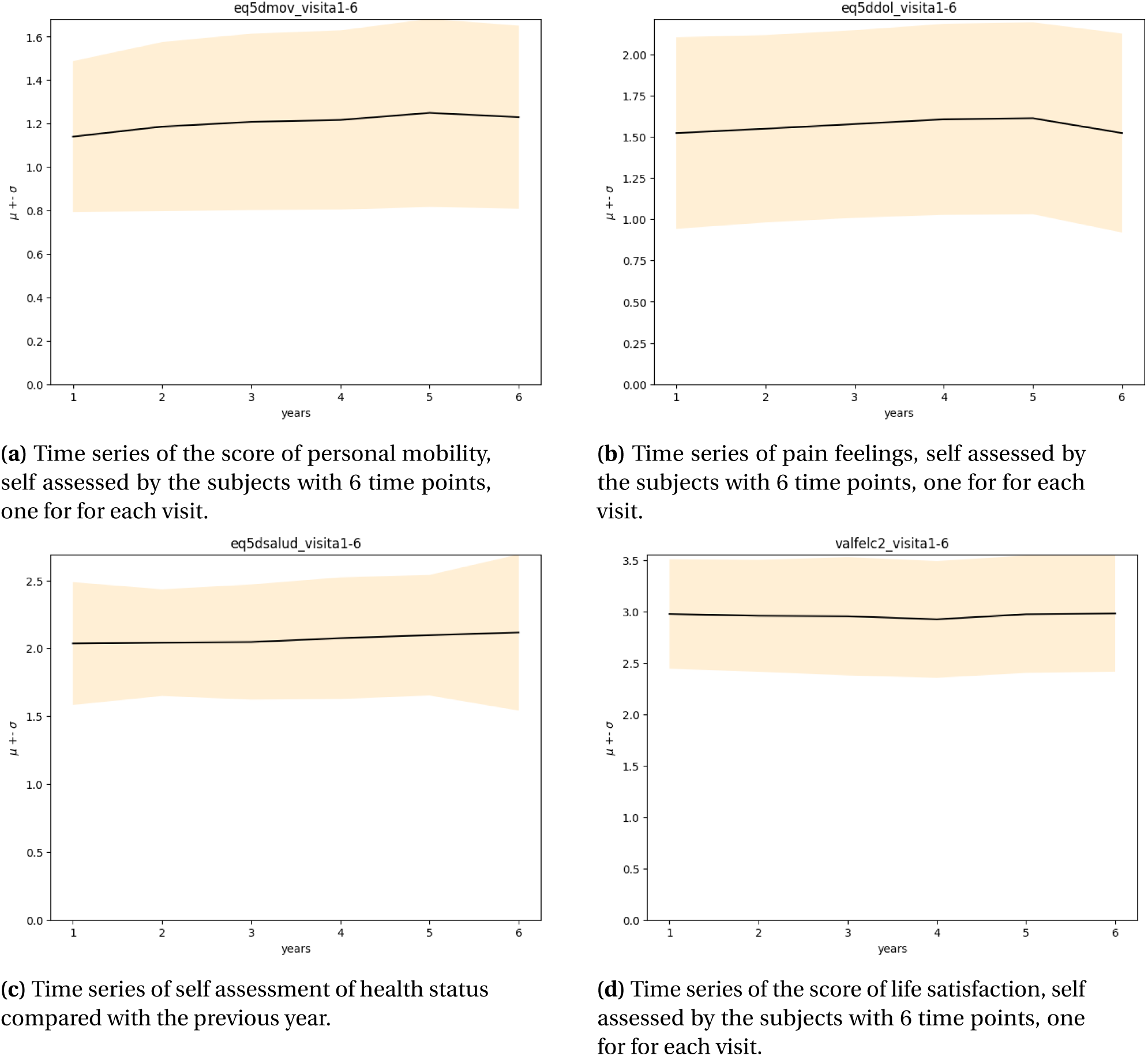
Time series of auto-informed scores of quality of life across the six years of *Vallecas Project* (no missing visits). The x-axis are time points, from year 1 to year 6 and the y-axis the score. Each plot represents the mean (bold line) and the dispersion or standard deviation (shaded region) across six years.

**Figure 24:**
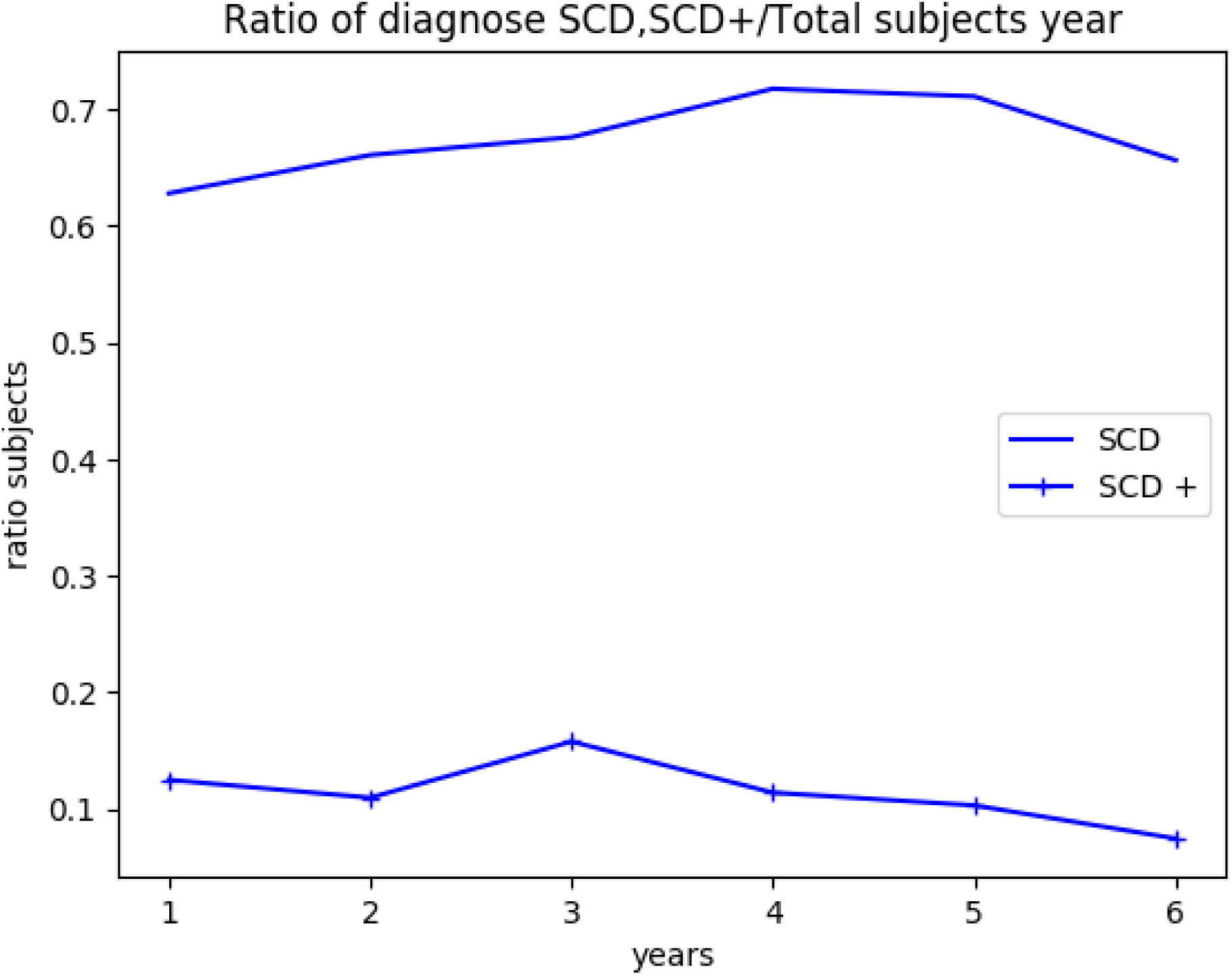
Ratio of subjects with Subjective Cognitive Decline (SCD) and Subjective Cognitive Decline Plus (SCD +) across the seven years of *Vallecas Project*.

### 3.5 Subjective Cognitive Decline

Figure 25 plots the time series for the Subjective Cognitive Decline (SCD). Subjective Cognitive Decline (SCD) is a term coined by [Jessen et al., 2014] and that tries to convey the decline in cognitive abilities experienced by the subject in comparison with prior self-assessed normal status by the same subject. SCD is a conceptual framework that has been used in the study of preclinical AD and independent of objective cognitive impairment [Ávila-Villanueva and Fernández-Blázquez, 2017].

**Figure 25:**
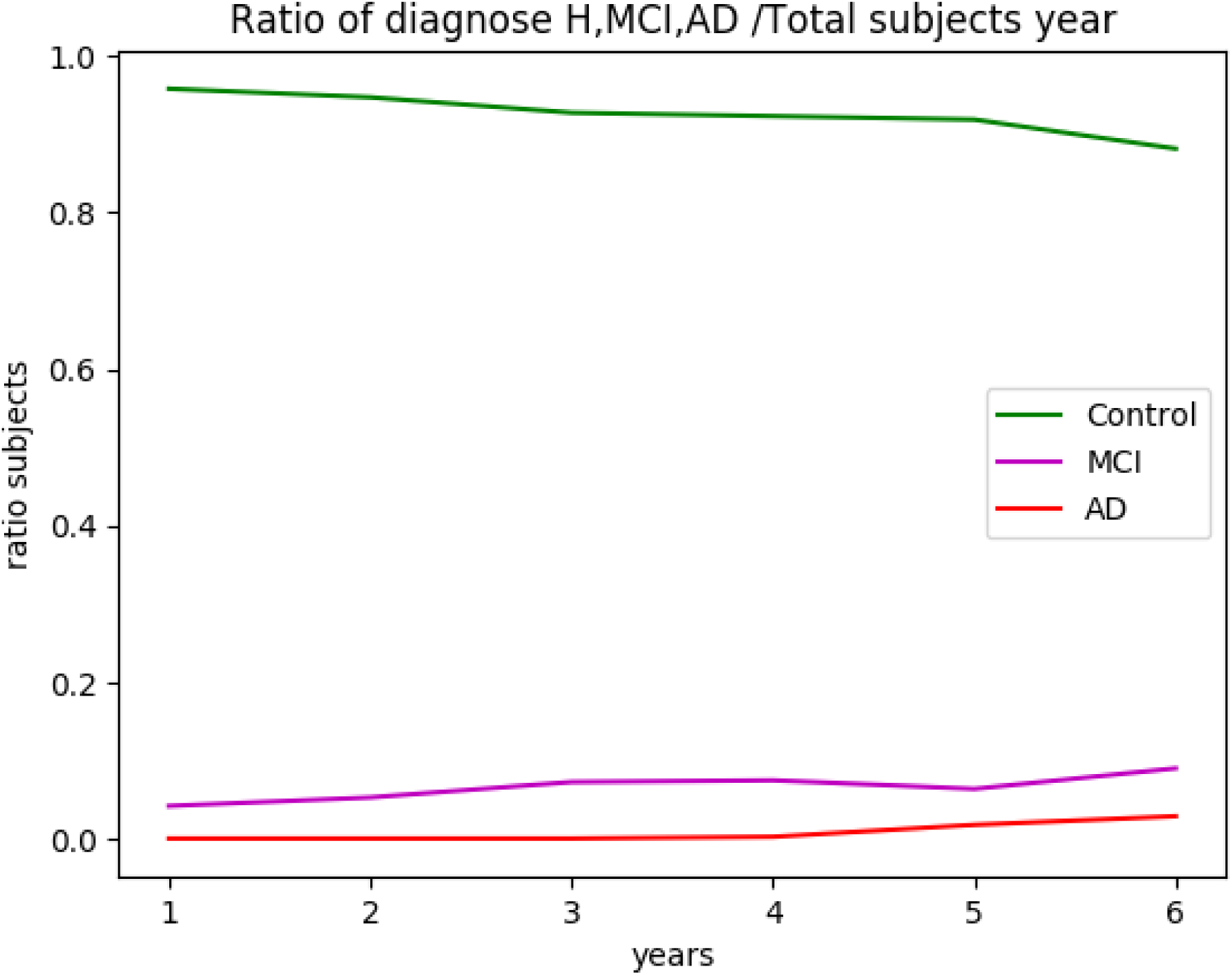
Ratio of subjects diagnosed as Healthy, Mild Cognitive Impairment (MCI) and Alzheimer’s disease (AD) across the seven years of *Vallecas Project*.

### 3.6 Diagnoses

Figure 25 plots the time series for the diagnoses administered to subjects for each year’s visit. The diagnosis includes 5 categories: Healthy, Subjective Cognitive Decline (SCD), Subjective Cognitive Decline Plus (SCD +), Mild Cognitive Impairment (MCI) and Alzheimer’s disease (AD).

### 3.7 Automatic Segmentation of Brain Structures

The segmentation pipeline used the FMRIB Software Library [fsl, 2019], [Jenkinson et al., 2012] and goes through the following stages: [Hannoun et al., 2018]

1. **Reorientation** to the standard (MNI) orientation
2. **Cropping** to remove the neck, this is a necessary step prior to brain extraction.
3. **Bias field correction**
4. **Registration** to standard space computing both linear registration (FLIRT algorithm) and non-linear registration (FNIRT algorithm)
5. **Brain Extraction** in two steps. First, the BET algorithm obtains a brain mask which can be coarse to then perform the brain extraction transforming standard-space mask to the input image using the FNIRT (non-linear) registration.
6. **Tissue Segmentation** (FAST algorithm)
7. **Subcortical Segmentation** (FIRST algorithm)

## 4 Bivariate analysis

In the previous sections we studied the dataset unidimensionally, that is, we plot distributions and compute statistics one variable at a time. Here we perform correlational analysis, that is, two or more dimensions, in order to study the effect of some variables upon others. Thus, the aim here is to try to come to grips with the mechanisms that could have generated the data, this is the realms of inferential statistics and is different from descriptive statistics and data visualization.

In what follows we go through each category of (cross-sectional) features previously described in Section 2 to via a visual inspection, investigate whether the features affect the rate of conversion to MCI. The contingency table [Everitt, 1992] will give us the (multivariate) frequency distribution of two categorical variables e.g. familial AD and conversion to MCI. Then, we quantify the putative effect indicated in the contingency table, using statistical inference tests which will gives us statistics e.g. chi-square or Fisher’s statistic from which a p-value is computed. We compute the Fisher’s exact test when the contingency table is 2 *×* 2 and the 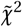 test otherwise [Jones et al., 01]. If the independent variable is not categorical (e.g. weight, defined in **ℝ**) we perform a one way ANOVA test. If the p-value is small enough to be considered relevant then we study whether the feature (e.g. familial AD) has an effect in brain capacity and cognitive reserve measured with brain structure volumes and the FCSRT score test scores respectively. This discovery process is described in Figure 26.

**Figure 26:**
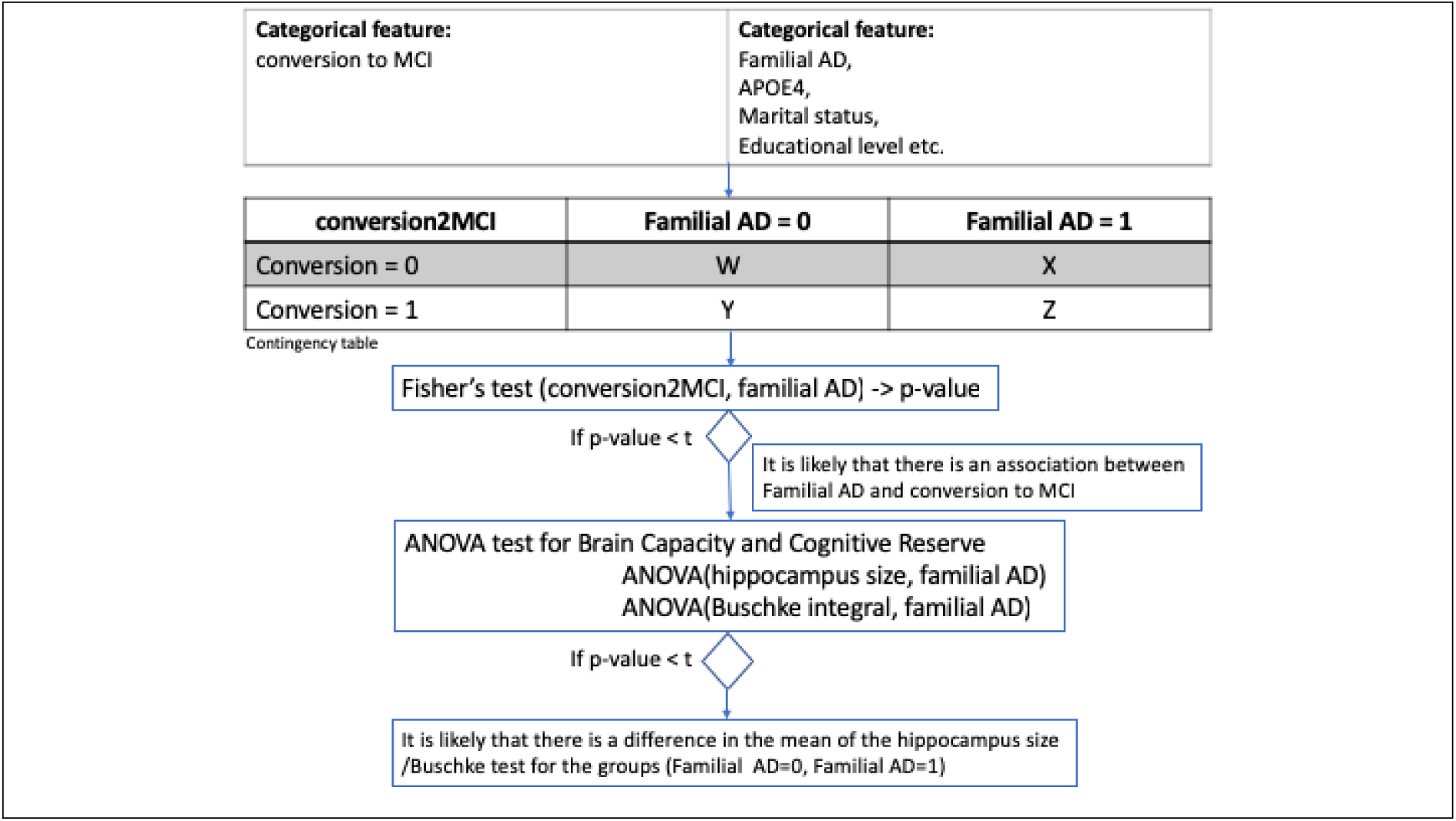
The figure shows the process followed to study whether there is an effect on conversion to MCI for the different categorical features studied in *Proyecto Vallecas*. A contingency table is a tool to quantify relationships via frequency counts between two categorical variables, e.g. conversion to MCI and familial AD. Significance tests (e.g. Fisher’s exact test [Fisher, 1937]or 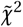) perform hypotheses testing about distributions of categorical data. For example, Is the distribution of converters different for familial AD versus non familial AD? Finally, for the categorical data with small p-values we analyze the differences among group means for variables related to *brain capacity* and *cognitive reserve*. For example, Is the mean of the hippocampus size Buschke Integral score different for familial AD?

The reader can go straight to the results of the statistical tests. Table 12 shows the p-values to asses the effect of categorical variables on conversion to MCI, using Fisher’s and 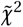 tests (Fisher’s test if the contingency matrix is 2*x*2 and 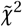 test otherwise). Table 13 asses the effect of non-categorical variables using one-way ANOVA, providing the p-values, the OLS estimator for *β* and the effect size.

### 4.1 Genetic, Anthropometric, Demographic

Figure 27 shows the results of grouping by *Conversion to MCI* relative to the *sex* and *hand laterality*. The contingency tables are shown in 3 and 4, respectively. The likelihood, via p-values, that a relationship between these two variables and conversion to MCI is caused by something other than chance are shown 12. The p-values are computed for two different scenarios: for subjects that have at least 2 visits (920 subjects) and for subjects that came to all their 6 visits (471 subjects).

**Figure 27:**
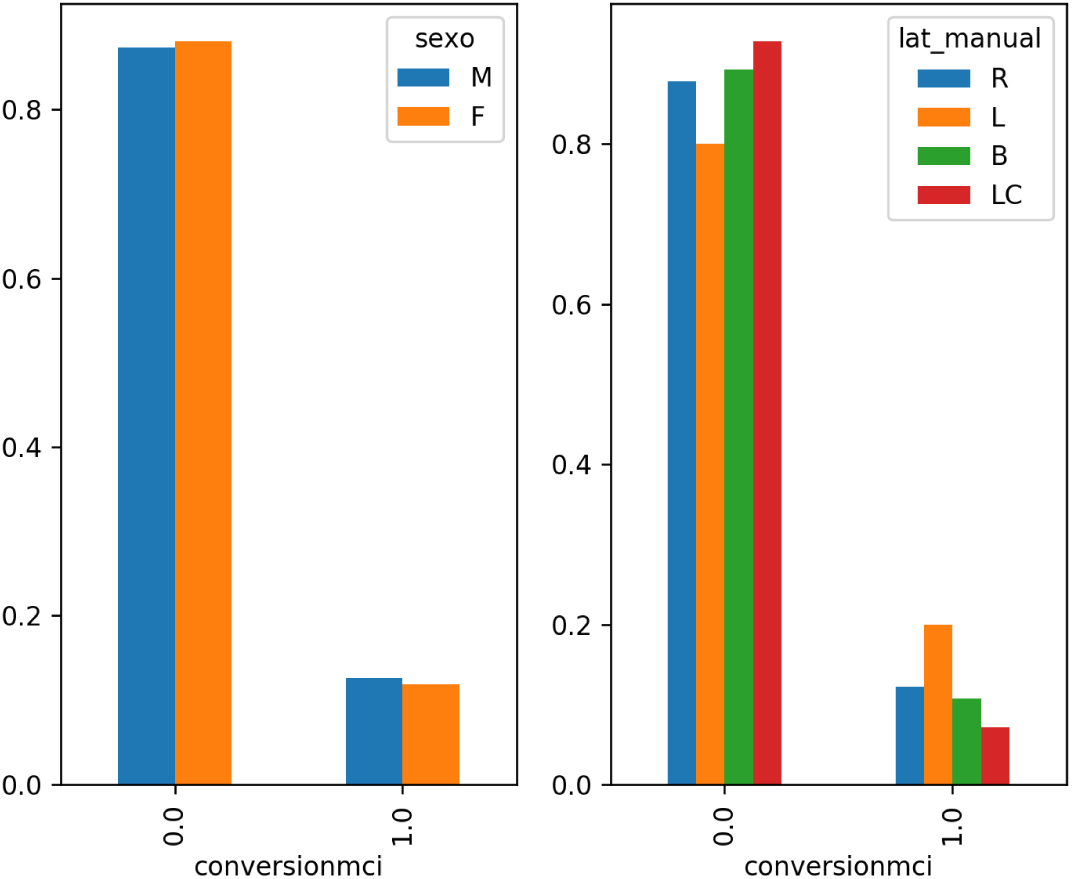
Frequency of sex and hand laterality related conversion to MCI in the future(for subjects with at least 2 visits). 0:non conversion to MCI, 1:conversion to MCI.

#### Statistical tests for sex and hand laterality

The contingency tables displaying the frequency distribution of *sex* and *MCI* and *hand laterality* and *MCI* are shown in 3 and 4 respectively. The significance of the difference between the proportion levels of sex is assessed using the Fisher’s exact test and 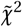 for hand laterality.

**Table 3:** Contingency table proportion of conversion to MCI conditioned and sex.

| Conversion | Male | Female |
| --- | --- | --- |
| No MCI | 165 | 265 |
| MCI | 15 | 28 |

**Table 4:** Contingency table conversion to MCI conditioned to hand laterality.

| Conversion | Right | Left | Ambi | LeftC |
| --- | --- | --- | --- | --- |
| No MCI | 365 | 7 | 14 | 7 |
| MCI | 38 | 1 | 1 | 1 |

No statistical significance is observed between these variables and conversion to MCI. See 12.

Figure 28 shows the results of grouping by *conversion to MCI* relative to the genetics profile. In essence, what we do is to split the data into groups depending on the variable APO*ϵ*4 (0:lack a copy of the APO*ϵ*4 allele, 1: one copy APO*ϵ*4 (heterozygotes), 2:two copies of APO*ϵ*4 (homozygotes)).

**Figure 28:**
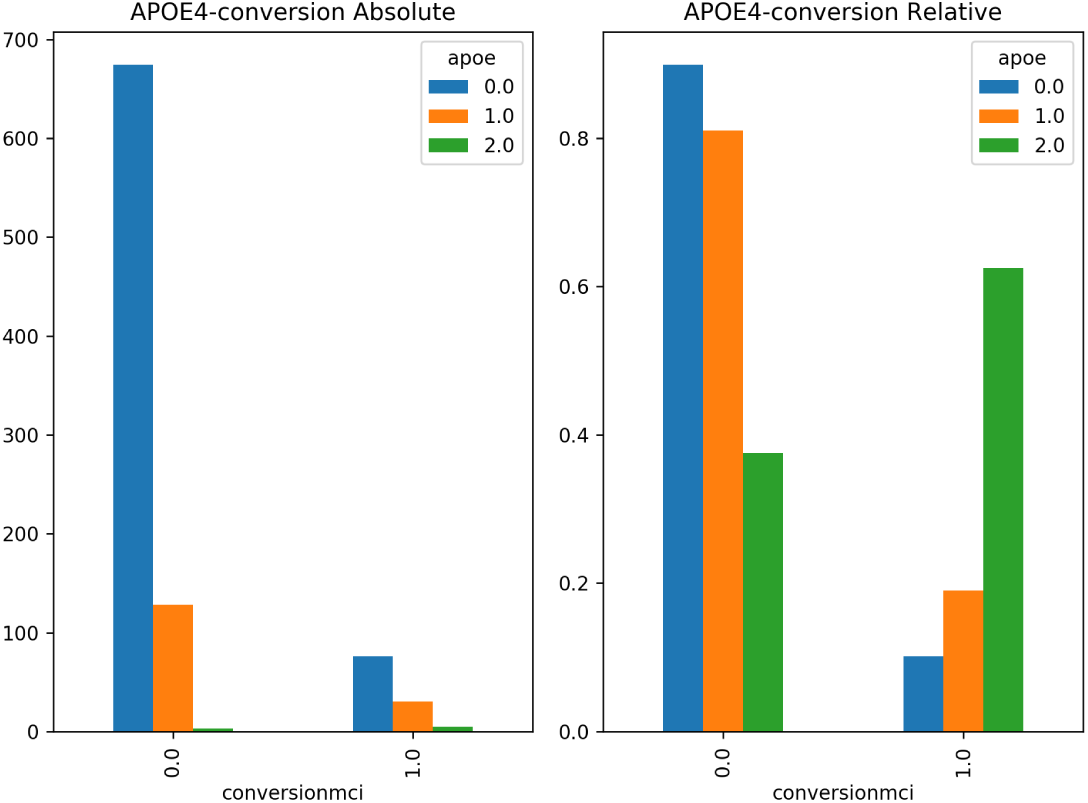
APO*ϵ*4 and conversion to MCI in the future(for subjects with at least 2 visits), represented as the genetic profile (APO*ϵ*4) conditioned to conversion.

Figure 29 shows the results of grouping by *conversion to MCI* relative to *familial AD* (0: non familial AD and 1: familial AD).

**Figure 29:**
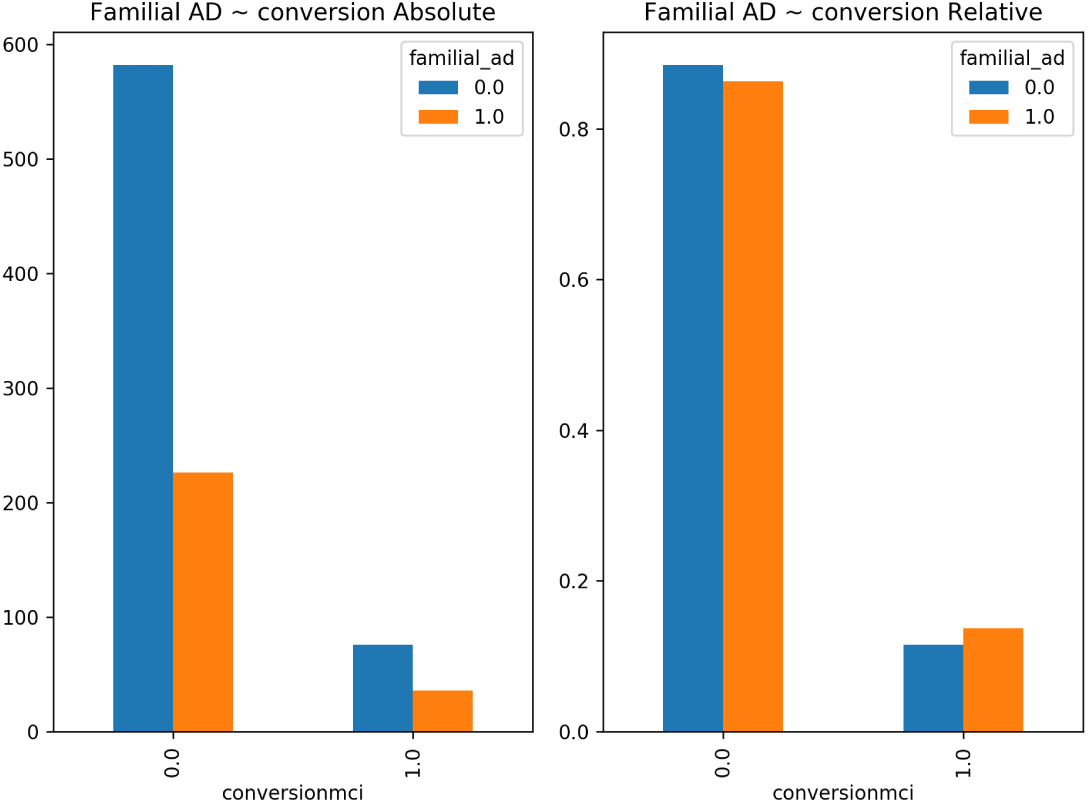
Grouping of conversion to MCI conditioned by familial AD.

Note that for each chart in Figures 28 and 29, on the left side the grouping is in absolute numbers and on the right side in relative numbers, the latter is a preferable representation since the dataset is unbalanced (many more non converters to MCI than converters). Note that the colored bars on the right sided charts must sum 1 (a frequentist could interpret this as a probability function) [Gomez-Ramirez and Sanz, 2013].

#### Statistical tests for genetic profile and MCI

The contingency table displaying the frequency distribution of APOE and MCI is shown in Table 5. The significance of the difference between the proportion levels is assessed using the 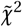 test. The imbalance observed in Table 5 is thus statistically significant and homozygotic APOE subjects have a larger conversion rate to MCI. Note that the frequency count for the homozygote cells (last column in Table 5) are very small (3 and 2).

**Table 5:** Contingency table showing the proportion of conversion to MCI conditioned to APOE*ϵ* (lack of allele, heterozygote, homozygote) for subjects with all their 6 visits.

| Conversion | APOEε = 0 | APOEε = 1 | APOEε = 2 |
| --- | --- | --- | --- |
| No MCI | 334 | 51 | 3 |
| MCI | 28 | 13 | 2 |

The contingency table displaying the frequency distribution of familial AD and MCI is shown in Table 6. The significance of the difference between the proportion levels is assessed using the Fisher’s exact test, *F =* 0.01882 which means that the probability that we would observe this or an even more imbalanced ratio in familial AD related to MCI by chance is about 2%. Nevertheless, when the test if performed for a longer N (subjects that came to at least two visits) we find a p-value above the 5% (Table 12).

**Table 6:** Contingency table showing the proportion of conversion to MCI conditioned to familial AD for subjects with all their 6 visits.

| Conversion | *FamilialAD* = 0 | *FamilialAD* = 1 |
| --- | --- | --- |
| No MCI | 265 | 135 |
| MCI | 20 | 23 |

Figure 30 shows the results of grouping by *conversion to MCI* relative to anthropometric features, i.e: *BMI, abdominal perimeter, weight* and *height*. We split the data into two groups (non converter vs converter to MCI) depending on the body mass index and related variables. The split is performed using the four quartiles of each variable as described in the figure’s legend. A one way ANOVA is performed for each variable separately, not observing any relevant conditioning on conversion to MCI. Although the p-value of BMI is below the 5% (only for subjects that came to at least 2 visits and not for those that came to all their visits), the effect is very small, *<* 0.1). See Table 13.

**Figure 30:**
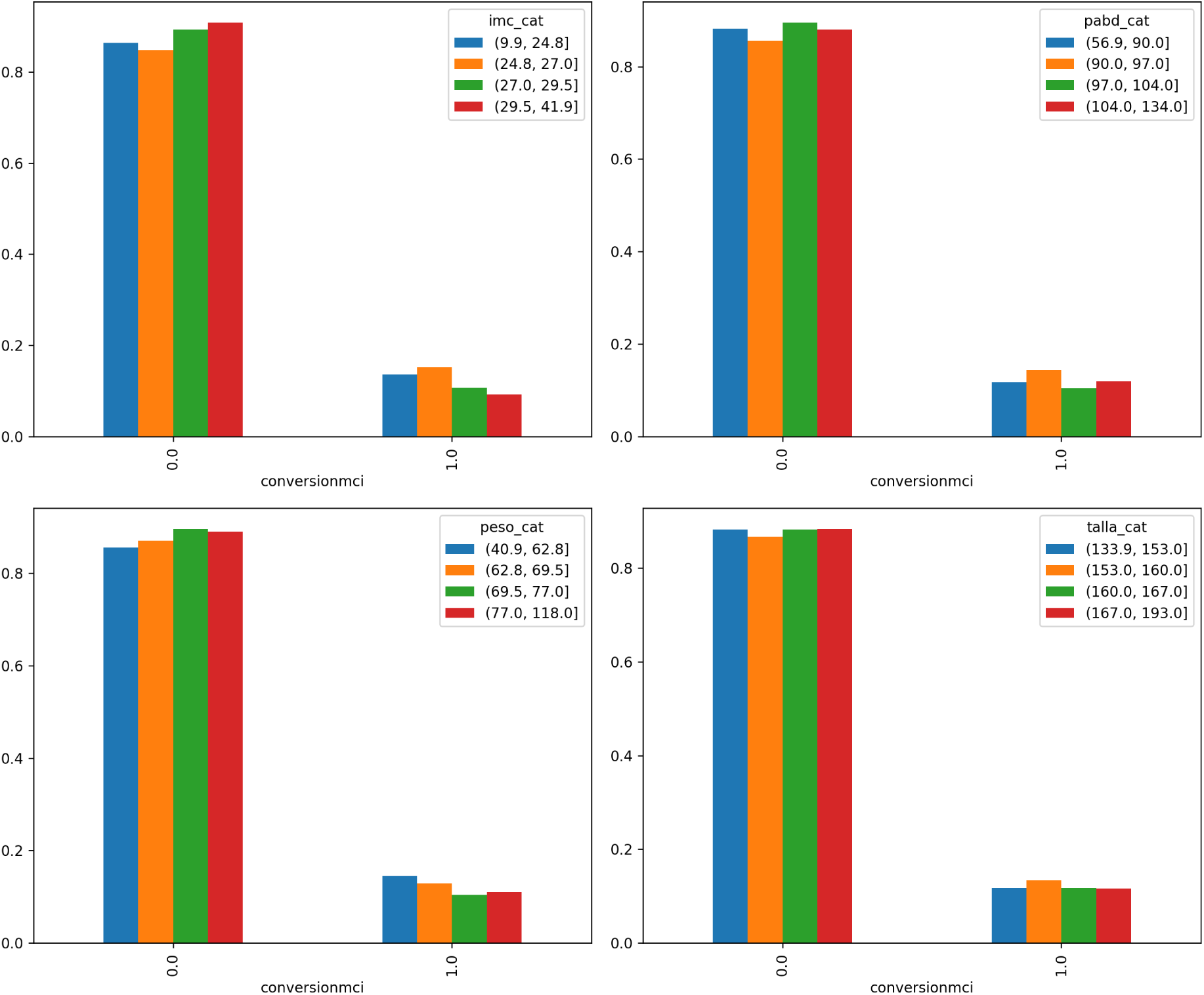
The figure shows the grouping by conversion to MCI relative to the body mass index, abdominal perimeter, weight and height (quartiles), all in relative numbers to acknowledge the fact that the dataset is unbalanced (the ratio non converters converters is 9 to 1).

Figure 31 shows the results of grouping by *conversion to MCI* relative to demographics information such as education level, family size, residence, marital status, self perceived socioeconomic status and work experience.

**Figure 31:**
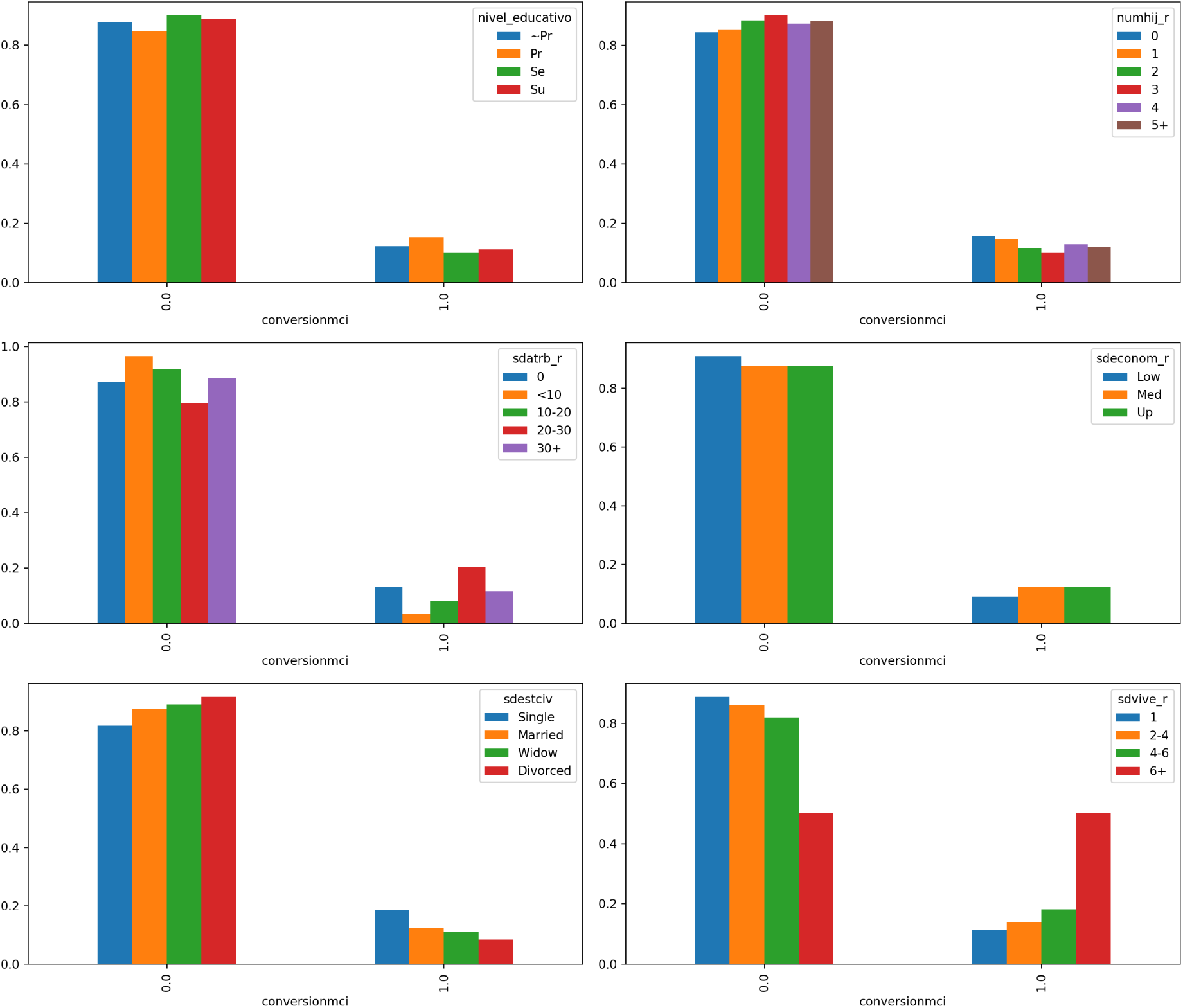
The figure shows the grouping by conversion to MCI relative to demographics information(for subjects with at least 2 visits). From up left to bottom right: conversion conditioned to educational level, number of sons, years of work as an employee, self perceived socioeconomic status, marital status, number of people living at home

The contingency tables displaying the frequency distribution of MCI and marital status is shown in Table 7. The significance of the difference between the proportion levels of marital status (single, married, widowed, divorced) is assessed using the Fisher’s exact test. We do not observe relevant conditioning of marital status on conversion to MCI (Table 12).

**Table 7:** Contingency table showing the proportion of conversion to MCI and marital status for subjects with all their 6 visits.

| Conversion | *Single* | *Married* | *Widowed* | *Divorced* |
| --- | --- | --- | --- | --- |
| No MCI | 35 | 237 | 87 | 32 |
| MCI | 4 | 25 | 11 | 3 |

The contingency tables displaying the frequency distribution of education level and MCI and wealth relative to MCI are shown in Tables 8 and 9, respectively. The significance of the difference between the proportion levels of educational level is assessed using the Fisher’s exact test and 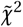 for wealth. We do not observe any relevant conditioning of educational level or wealth (residency) on conversion to MCI (Table 12).

**Table 8:** Contingency table proportion of conversion to MCI and educational level.

| Conversion | No | Prim | Sec | Uni |
| --- | --- | --- | --- | --- |
| No MCI | 112 | 126 | 57 | 96 |
| MCI | 11 | 12 | 5 | 13 |

**Table 9:** Contingency table proportion of conversion to MCI and wealth estimation based on residency.

| Conversion | Low | Med | High |
| --- | --- | --- | --- |
| No MCI | 82 | 243 | 66 |
| MCI | 5 | 31 | 7 |

### 4.2 Sleep

Figure 32 shows the results of grouping by *conversion to MCI* relative to information about sleep patterns self-reported by the subjects.

**Figure 32:**
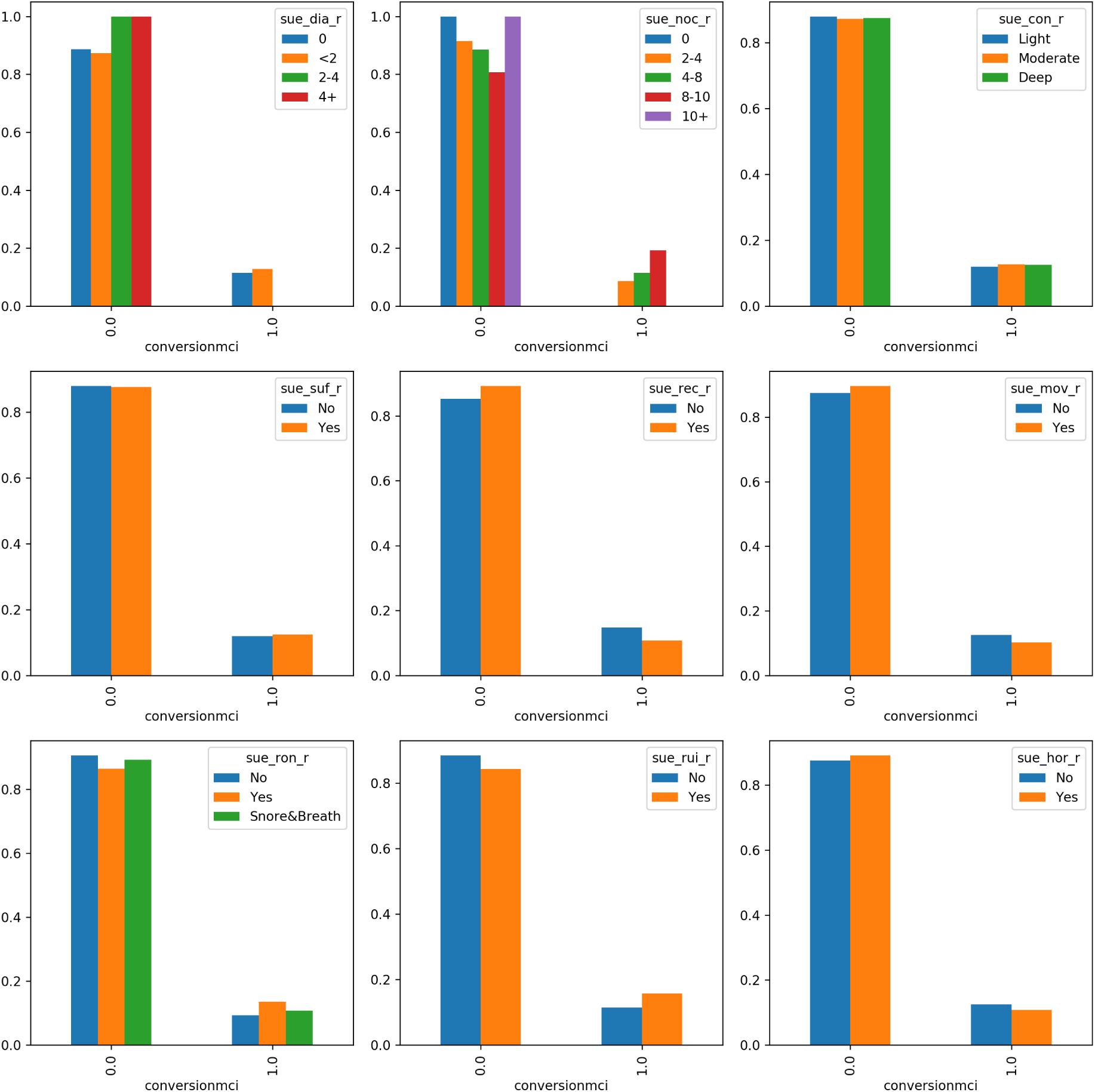
The figure shows the grouping by conversion to MCI relative to sleeping patterns. From up left to bottom right: conversion conditioned to number of hours of daily sleep, number of hours of sleep at night, loose of consciousness while sleep, sufficient sleep, remember dreams, movement while sleep, snoring, noises and tingling.

The contingency tables displaying the frequency distribution of *deep sleep* and MCI and *remember dreams* and MCI are shown in 10 and 11, respectively. We do not observe any relevant effect of sleep related variables on conversion to MCI (Table 12).

**Table 10:** Contingency table proportion of conversion to MCI and deep sleep (*Light, Moderate, Acute*).

| Conversion | Li | Mod | Acu |
| --- | --- | --- | --- |
| No MCI | 74 | 161 | 154 |
| MCI | 7 | 22 | 14 |

**Table 11:** Contingency table proportion of conversion to MCI and remember dreams (*No, Yes*).

| Conversion | N | Y |
| --- | --- | --- |
| No MCI | 115 | 276 |
| MCI | 42 | 68 |

### 4.3 Life style: food and diet, physical exercise and social bonding

Here we study the grouping by *Conversion to MCI* relative to information about life style patterns. Life style is a quite vague category where completeness is certainly unwarranted. Nevertheless, *The Vallecas Project* makes available a representative set of factors including, food consumption and diet, physical exercise and social bonding.

Figure 33 shows the results of grouping by *Conversion to MCI* relative to food consumption in weekly basis. We outline four group foods: fruits, vegetables, red meat and sweets. A visual inspection shows no effect of food consumption on conversion to MCI.

**Figure 33:**
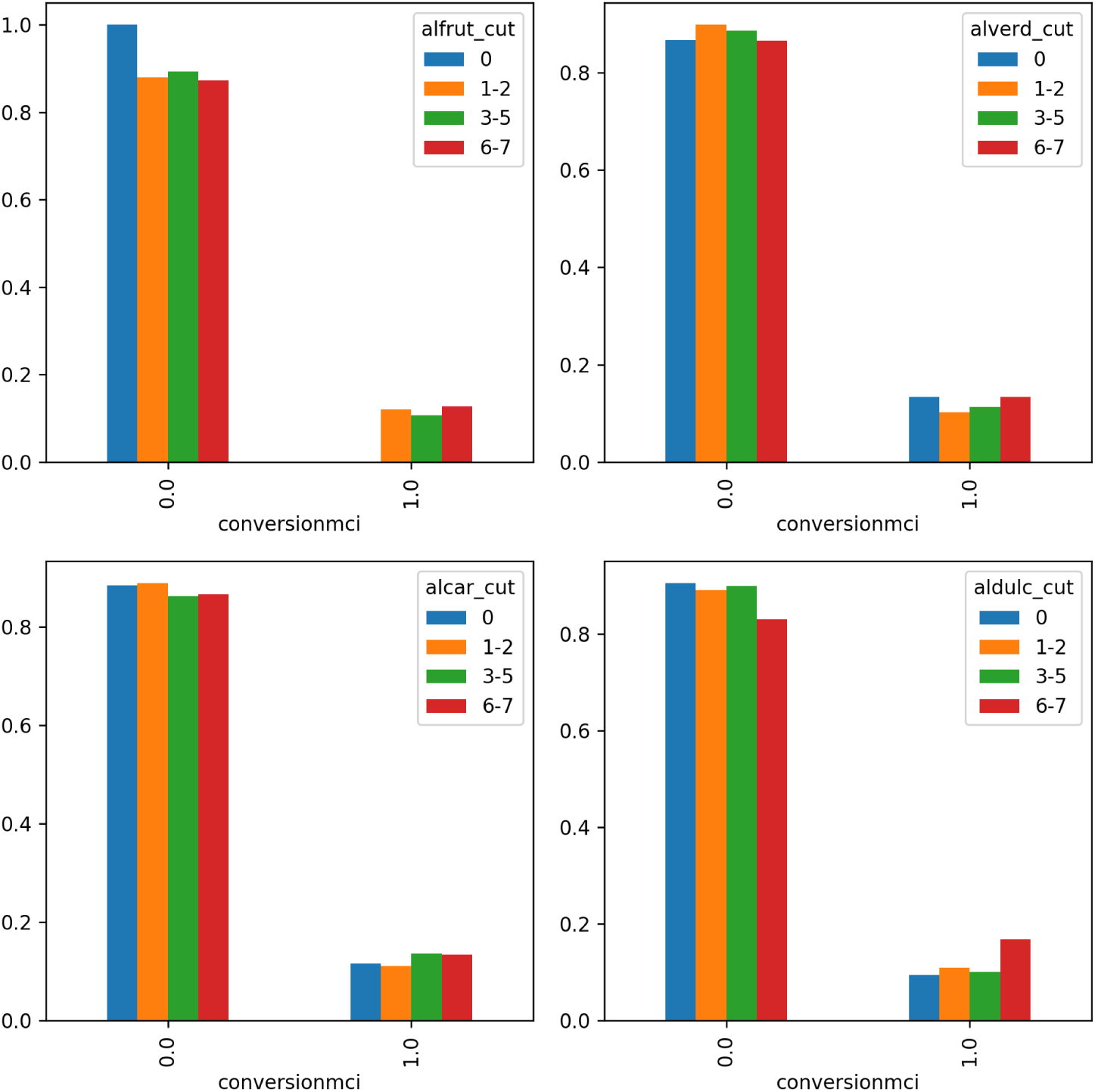
The figure shows the grouping by conversion to MCI relative to food consumption in weekly basis. From up left to bottom right: fruits, vegetables, red meat and sweets. The majority of subjects,63%, claims to eat fruit 6-7 days/week, 35% vegetables, only 4% red meat and 24% claim to eat sweets 6-7 days/week.

Figure 34 shows the results of grouping by *conversion to MCI* relative to diet. We distinguish between four types of diets based on food consumption: high in sugar, high in proteins, high in fruits and vegetables (Mediterranean) and high in fats. A visual inspection shows no effect of the type of diet on conversion to MCI, except for the bottom right plot (diet high in fats) which shows a very high rate of converters (50%). This appreciation is confirmed when we perform 1 way ANOVA tests(Table 13), we obtain values bellow 1%, however the effect size is too small (*<* 0.01).

**Figure 34:**
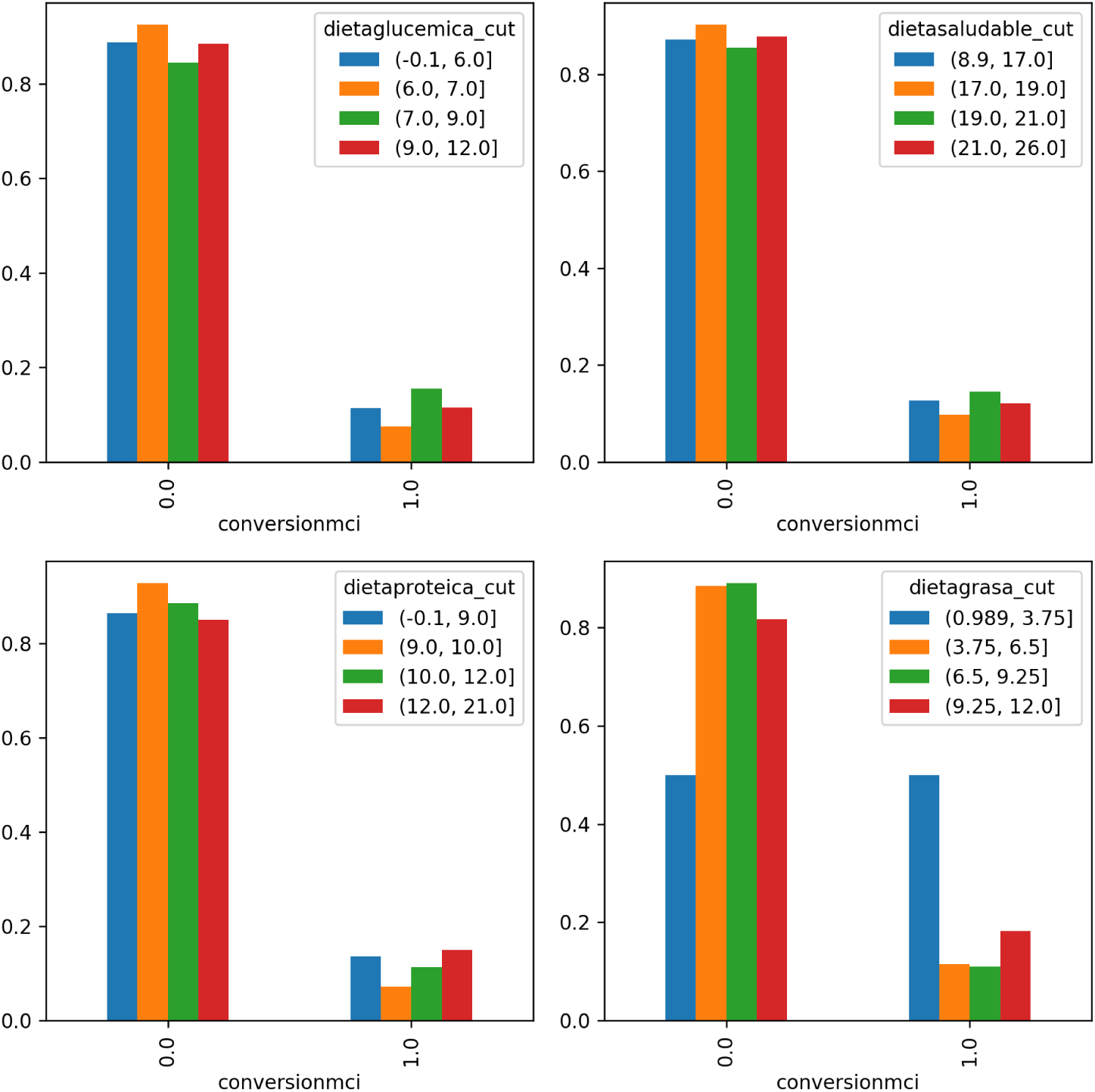
The figure shows the grouping by conversion to MCI relative to diet. From up left to bottom right: weekly consumption of fruits, vegetables, red meat and sweets.

Figure 35 shows the results of grouping by *Conversion to MCI* physical exercise. The physical activity is computed as the product of the number of sessions per week and the average minutes per session. We do not observe relevant conditioning of physical exercise on conversion to MCI (Table 13).

**Figure 35:**
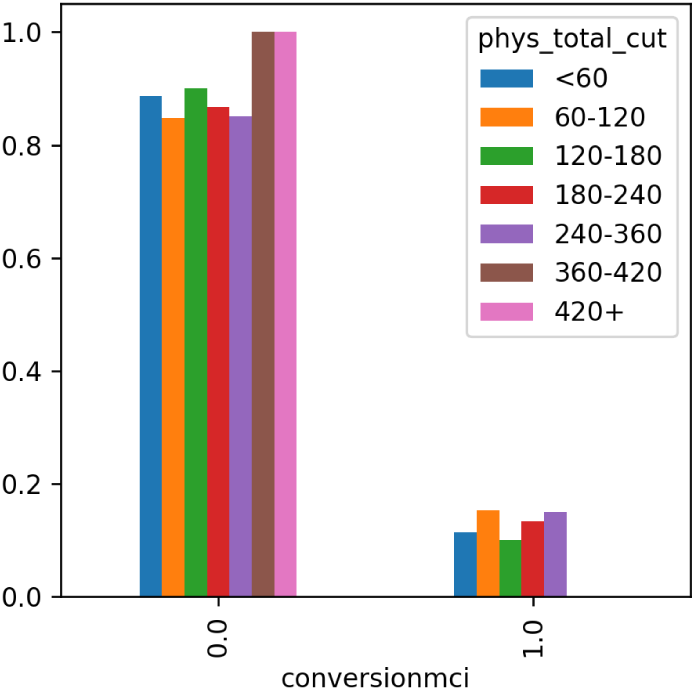
The figure shows the grouping by conversion to MCI relative to physical exercise (minutes of physical exercise per week).

Figure 36 shows the results of grouping by *conversion to MCI* relative to activities referred to social life and the subject’s engagement with the external world. A quick visual inspection seems to indicate a lesser conversion ratio for regular users of the Internet and new technologies and those that often go to the movie theater and to sport events, the opposite if observed for church goers.

**Figure 36:**
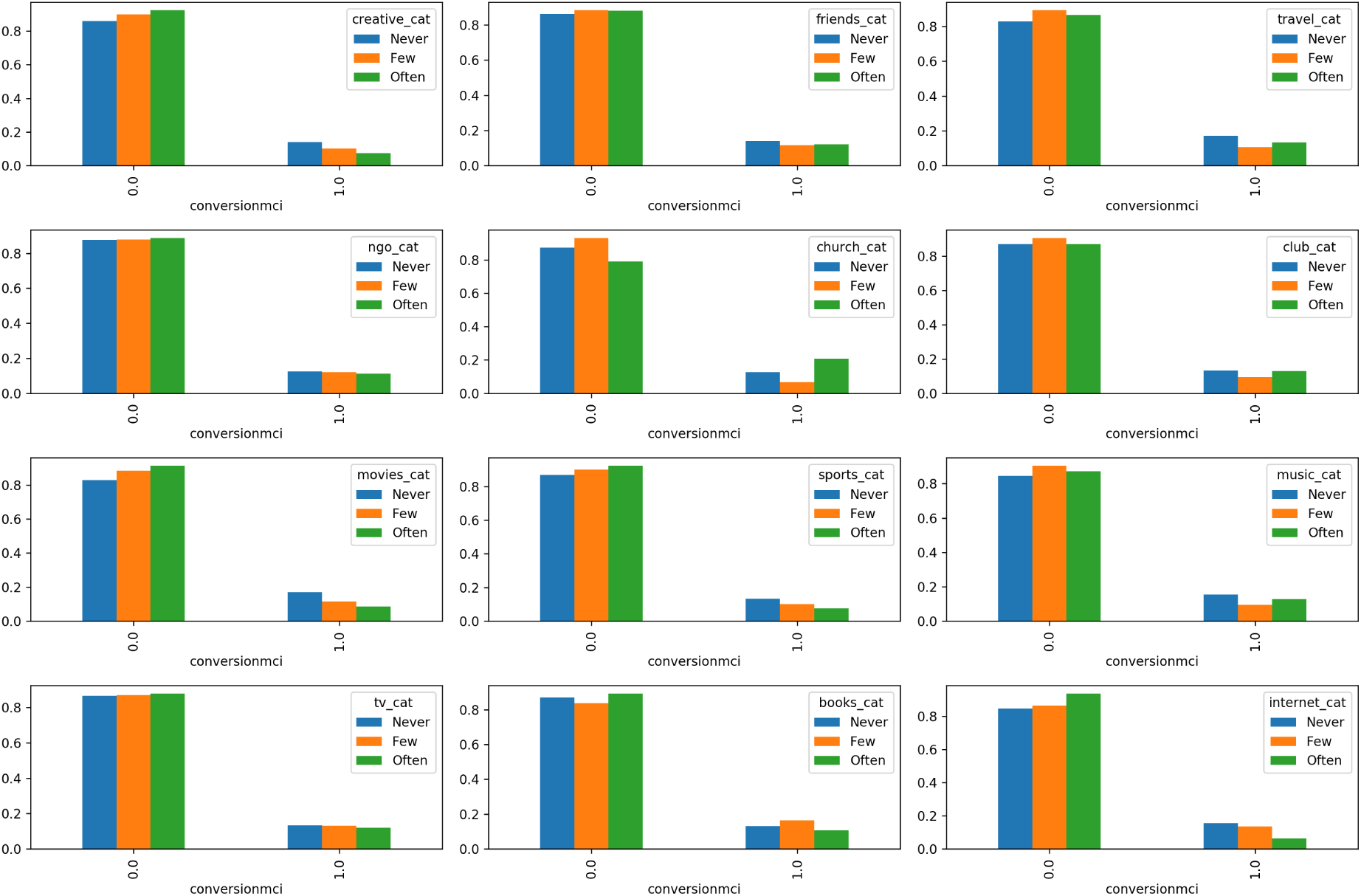
The figure shows the grouping by conversion to MCI relative to activities that reflect the subject’s social life and her engagement with the external world. From up left to bottom right the frequency (Never, Few, Often) of participating in: creative-artistic activities, go out with friends, traveling, participation in NGO and other non-profit bodies, go to church, social clubs, movie theater, sports events, music concerts, watching TV, reading (newspapers and books) and use the Internet.

The statistical tests for life style features are included in Table 12. We observe relevant conditioning on conversion to MCI for going to church and also for going to the movie theater and using the Internet and IT. Using the Internet is relevant for subjects with at least 2 visits (not when all the 6 visits are accounted for) and the reverse occurs for traveling, significant for visits coming to all their visits not for those that come to at least 2.

### 4.4 Cardiovascular and cerebrovascular pathologies

Figure 37 shows the results of grouping by *conversion to MCI* relative to the cardiovascular health and cerebrovascular of the subjects. A quick visual inspection seems to indicate a higher conversion ratio for subjects with heart problems (e.g. infarction) but not for overweight or smokers. This observation is not confirmed when we perform statistical tests (Table 12). The statistical tests for cardiovascular and cerebrovascular features are included in Table 12. We do not observe relevant conditioning of cardiovascular and cerebrovascular pathologies on conversion to MCI with the exemption of thyroid disorders (Table 13).

**Figure 37:**
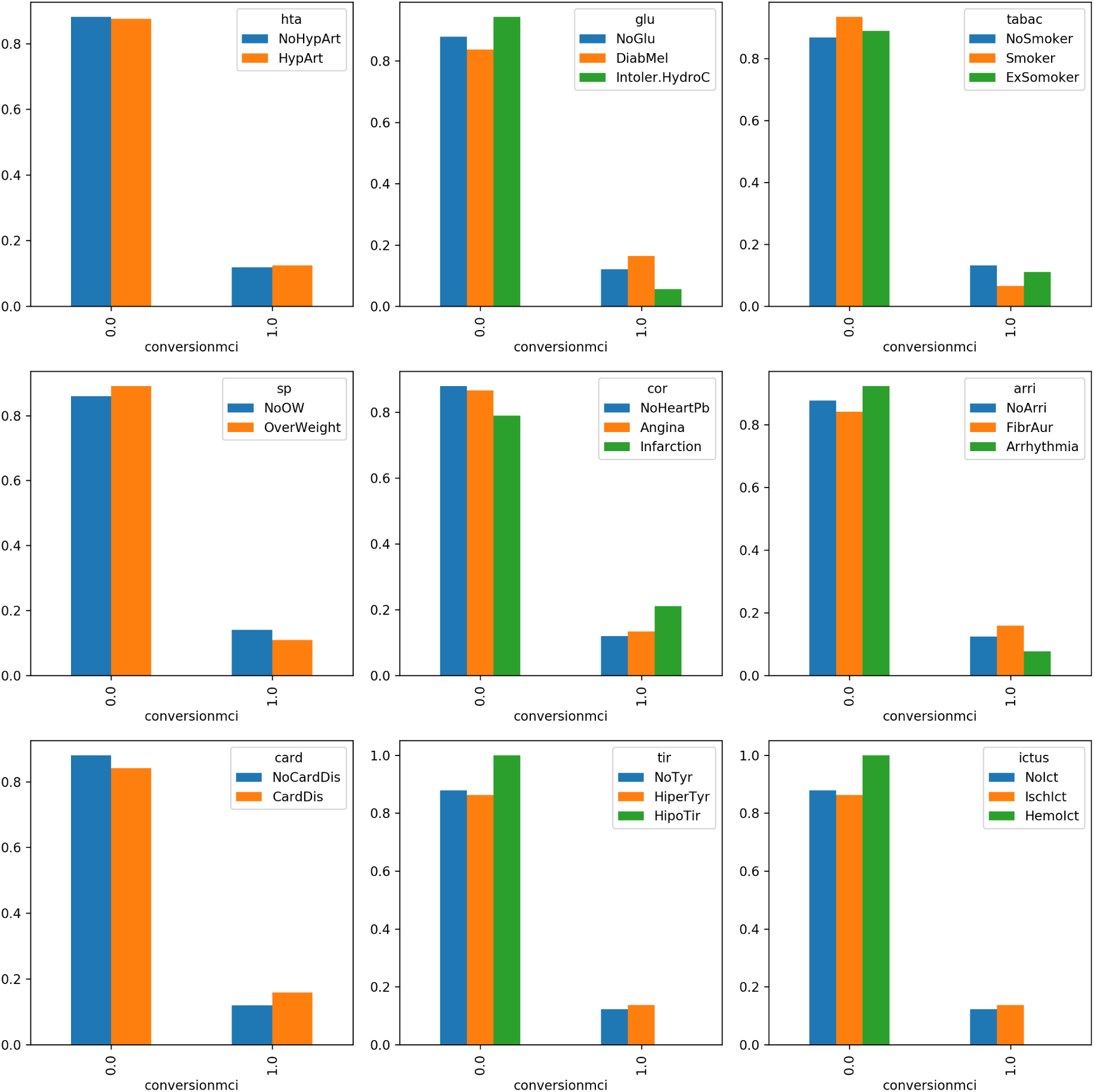
The figure shows the grouping by conversion to MCI relative to the cardiovascular health of the subjects. From up left to bottom right: blood pressure (*Normal blood preassure, hypertension*), diabetes (*No diabetic, diabetes mellitus, intolerance to carbs*), smoke (*Non smoker, Smoker, Ex-smoker*), body mass index (*No overweight, overweight*), cholesterol (*No cholesterol, hyper cholesterol, Hyper triglycerides, both hyper cholesterol and triglycerides*),arrythmias (*No Arrhythmia, Atrial fibrillation, Arrhythmia*), heart stroke (*No past strokes, Angina, Stroke*), thyroid problems (*No thyroiditis, Hyper thyroiditis, hipo thyroiditis*), ictus (*No, Ischemic, Hemorrhagic*).

Figure 38 shows the results of grouping *Conversion to MCI* by episodes of traumatic brain injury. A quick visual inspection seems to indicate a higher conversion ratio for subjects a history of TBI incidents which is not confirmed when we perform statistical tests (Table 12).

**Table 12:** Statistical tests to study whether a hypothetical observed imbalance of conversion to MCI is statistically significant. The first column shows the variable used to compute its effect upon conversion, second column shows the p-value for the test performed for those subjects that came at all their 6 visits (*N =* 471), the third column contains the p-value for those subjects that came at least to 2 visits (*N =* 960) and fourth column indicates the type of test performed (Fisher’s when contingency matrix is 2 *×* 2 and 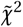 otherwise.)

| Feature | All 6 visits | At least 2 visits | Test |
| --- | --- | --- | --- |
| APOE | ** | ** | $\tilde{\chi}^2$ |
| Familial AD | * | 0.372 | F |
| Sex | 0.4163 | 0.7545 | F |
| Hand Lat | 0.0543 | 0.7420 | $\tilde{\chi}^2$ |
| Marital status | 0.9590 | 0.2769 | $\tilde{\chi}^2$ |
| Education | 0.5303 | 0.2950 | $\tilde{\chi}^2$ |
| Wealth(residency) | 0.3162 | 0.6742 | $\tilde{\chi}^2$ |
| Rem dreams | 0.5908 | 0.0988 | F |
| Suff. sleep | 0.4543 | 0.9082 | F |
| Snoring | 0.2944 | 0.3675 | F |
| Noises sleep | 0.0172 | 0.2059 | F |
| Moves sleep | 1 | 0.5688 | F |
| Tingling sleep | 1 | 0.7573 | F |
| Deep sleep | 0.4674 | 0.8898 | $\tilde{\chi}^2$ |
| Creative act. | 0.9083 | 0.4076 | F |
| Friends | 0.8306 | 0.6810 | F |
| Travel | * | 0.0726 | $\tilde{\chi}^2$ |
| NGO | 0.8234 | 0.9089 | $\tilde{\chi}^2$ |
| Church | ** | ** | F |
| Social club | 0.5873 | 0.3236 | F |
| Movie theater. | * | * | $\tilde{\chi}^2$ |
| Sports events. | 0.0983 | 0.28294 | F |
| Music | * | 0.1031 | F |
| TV radio | 0.8223 | 0.9435 | F |
| Books | 0.7887 | 0.0878 | F |
| Internet | 0.2191 | ** | F |
| Arrhythmia | 0.4234 | 0.5224 | $\tilde{\chi}^2$ |
| Blood press. | 0.6340 | 0.7631 | F |
| Diabetes | 0.8399 | 0.14987 | $\tilde{\chi}^2$ |
| Ictus | 0.7351 | 0.6641 | $\tilde{\chi}^2$ |
| Cholesterol | 0.6156 | 0.9667 | $\tilde{\chi}^2$ |
| Smoke | 0.1766 | 0.3150 | $\tilde{\chi}^2$ |
| Thyroid. | * | * | $\tilde{\chi}^2$ |
| Glucose metab.dis. | 0.8399 | 0.1498 | $\tilde{\chi}^2$ |
| Stroke | 0.5698 | 0.48274 | $\tilde{\chi}^2$ |
| Cholesterol | 0.6156 | 0.9667 | $\tilde{\chi}^2$ |
| Card. dis. |  | 0.32429 | F |

**Table 13:** Statistical tests (ANOVA) for non categorical variables. The first column shows the variable used to compute its effect upon conversion, second, third and fourth columns show the p-value, the coefficient regression (*β*) and the effect size(*ω*^2^) for those subjects that came at all their 6 visits (*N =* 471), similarly for the last three columns, now for those subjects that came at least to 2 visits (*N =* 960).

| 1-ANOVA | All 6 visits |  |  | At least 2 visits |  |  |
| --- | --- | --- | --- | --- | --- | --- |
| Feature | p-val | $\beta$ | $\omega^2$ | p-val | $\beta$ | $\omega^2$ |
| BMI | 0.7687 | 0.0013 | -0.0021 | * | -0.0066 | 0.0041 |
| Weight | 0.8271 | 0.0003 | -0.0021 | 0.123 | -0.0015 | 0.0015 |
| Height | 0.899 | -0.0002 | -0.0022 | 0.639 | 0.0002 | -0.0010 |
| Perim.abd. | 0.308 | 0.0015 | 0.00009 | 0.143 | -0.0003 | -0.0009 |
| Diet Gluc. | * | -0.0054 | 0.00132 | 0.065 | 0.0065 | 0.00053 |
| Diet Fats | ** | -0.0113 | 0.0011 | ** | -0.0054 | -0.0004 |
| Diet Med. | 0.8689 | -0.0009 | -0.0022 | 0.086 | -0.0009 | -0.00103 |
| Diet Prot. | * | -0.0030 | -0.0017 | * | 0.0017 | 0.00016 |
| Phys. Exercise. | 0.160 | -0.0002 | 0.00224 | 0.680 | 0.0000 | -0.0009 |

**Figure 38:**
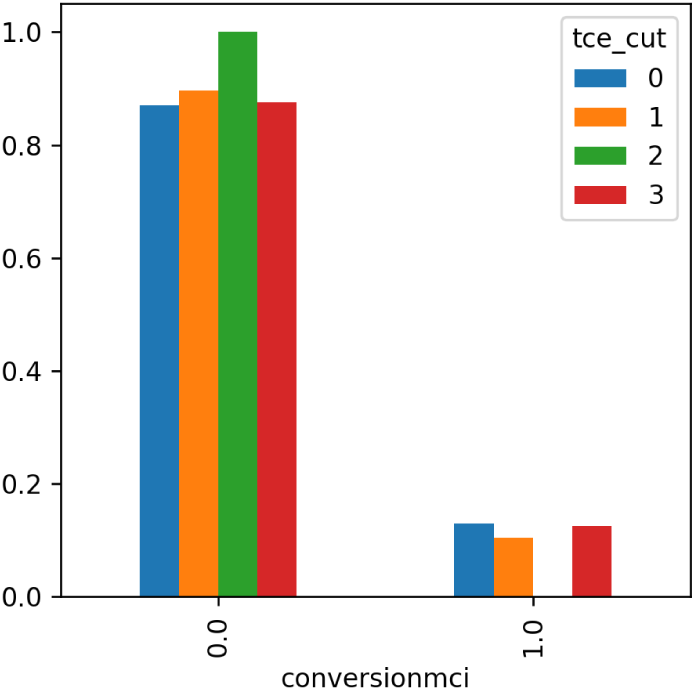
The figure shows the grouping by conversion to MCI relative episodes of traumatic brain injury.

#### Fisher and 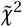 **tests**

Table 12 with statistical tests for both a population of the subjects with 6 visits and also in the second column for subjects that came to at least 2 visits.

## 5 Brain atrophy

In this section we show the analyze the volume estimates for brain extraction, tissue segmentation (CSF, white and gray matter) and subcortical segmentation (thalamus, putamen, hippocampus, accumbens, pallidum, caudate, amygdala) for a total of 314 subjects for both year 1 and year 6, (total 314 *×* 2 *=* 628).

### Brain extraction

Research shows that the growth rate of the brain slowly declines from the first year of birth (it increases dramatically during), brain volume reaches its maximum at the age of 40 years to then decline by 5% per decade, the decline is thought to accelerate at around 70 years [Peters, 2006].

Figure 39 shows the results of the brain extraction algorithm for years 1 and year 6. The brain’s size is expressed in cm^3^ in native space. Left side, brain volume density estimate for year 1 and on the right, for year 6.

**Figure 39:**
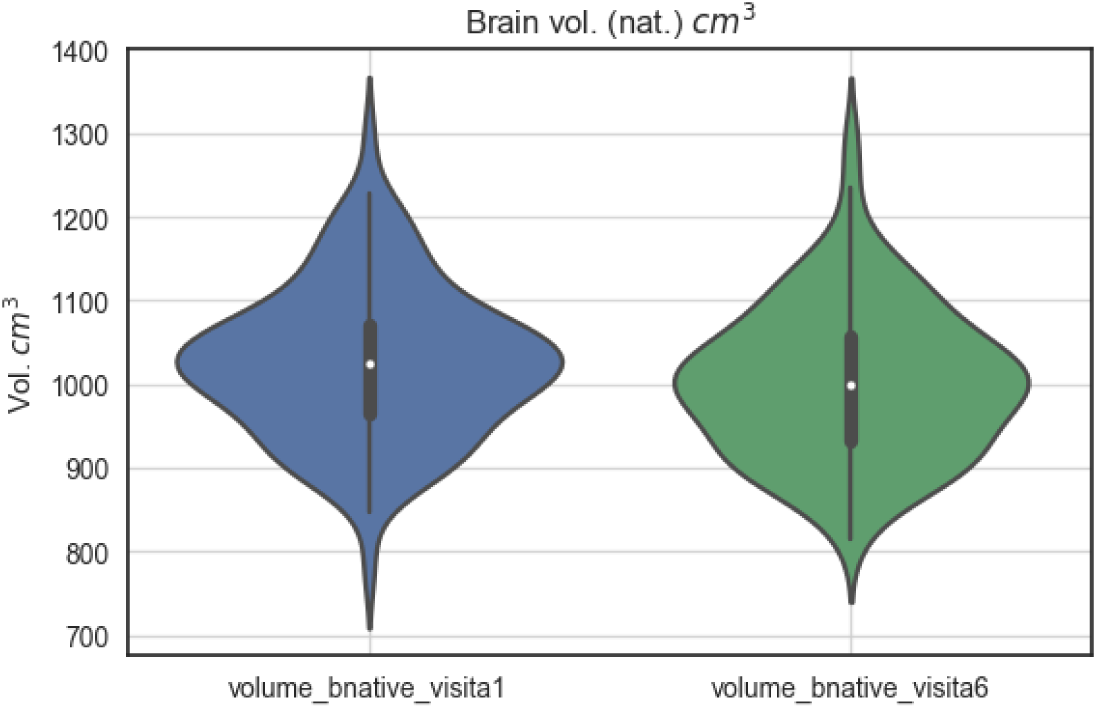
The violin plot shows the brain volume in native space (*cm*^3^) for year 1 (left) and year 6 (right). As expected, the mean volume decreases from year 1 to year 6. The average brain size is around 1,000 cm^3^ which reflects the age and the gender unbalance of the sample.

Since we have the brain volumes in two time points, we can calculate brain atrophy as the difference between the estimated volume at year 1 and the estimated volume at year 6. The atrophy between year 1 and year N for a volume *V* is calculated with the expression:

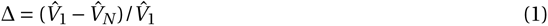

Figure 40 shows the atrophy for MCI converters and non MCI converters, grouped by sex and familial AD.

**Figure 40:**
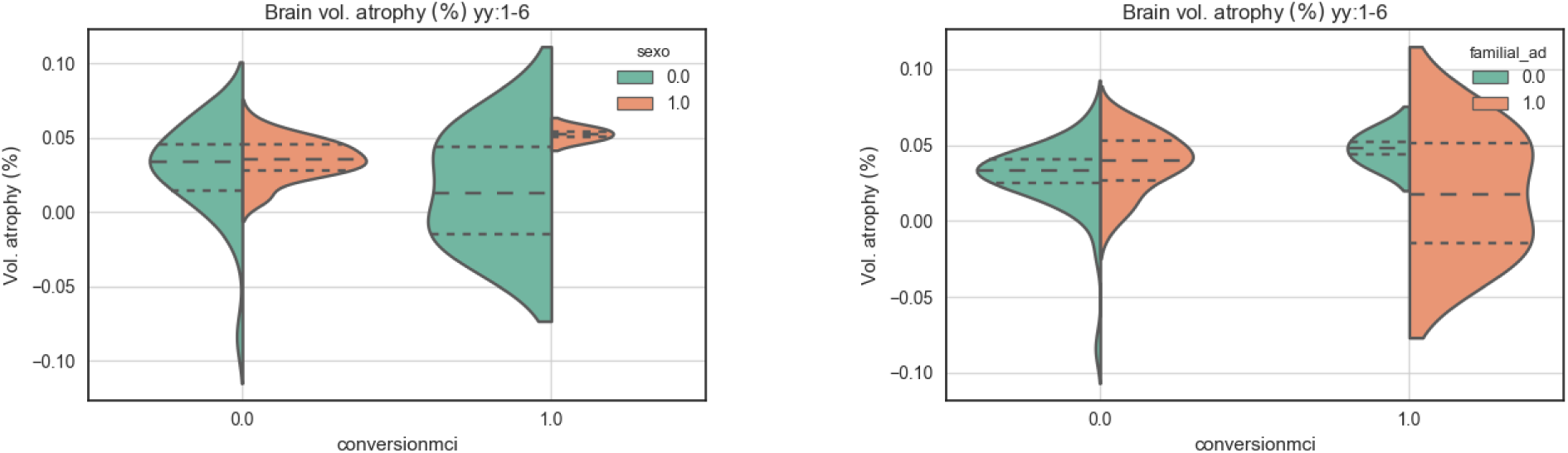
On the left, brain atrophy for converters and non converters grouped by familial sex (0:Male, 1:Female) and on the right grouped by familial AD.

### Segmentation of brain tissue

Figure 41 shows the segmentation based on tissue (CSF (green), gray matter(blue); white matter (red-yellow)) of the same subject of the *Vallecas Project*, in year 1 (left) and in year 6 (right).

**Figure 41:**
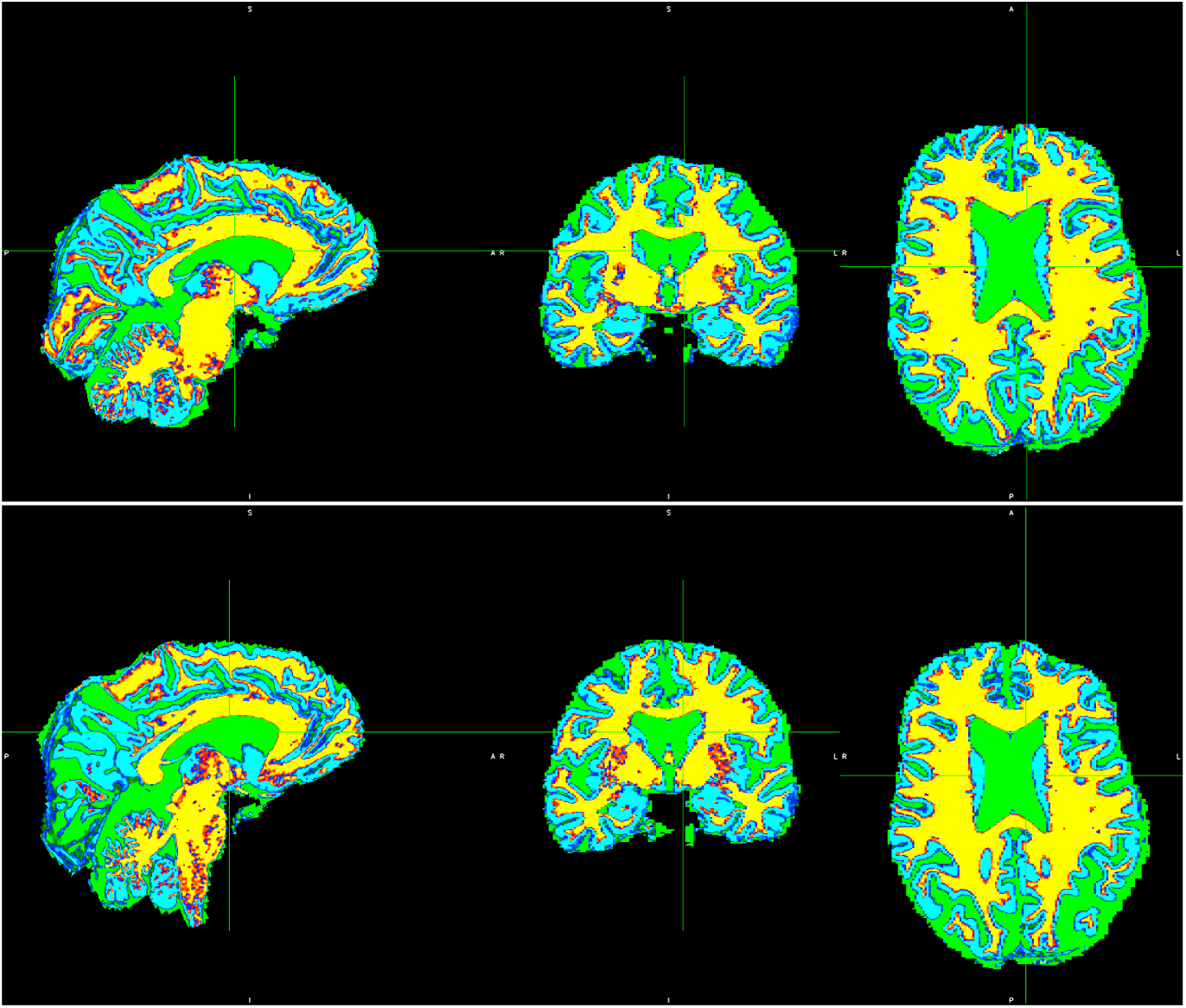
Tissue segmentation of one subject, on the left first year visit and on the right sixth year visit.

Figure 42 shows the estimated volume for tissue based segmentation across two time points, year 1 and year 6. The volume distribution of CSF (left), gray matter (middle) and white matter(right) are shown. Interestingly, the figure shows an increase in the volume of the CSF and at the same time a decrease in the brain parenchyma (white and gray matter).

**Figure 42:**
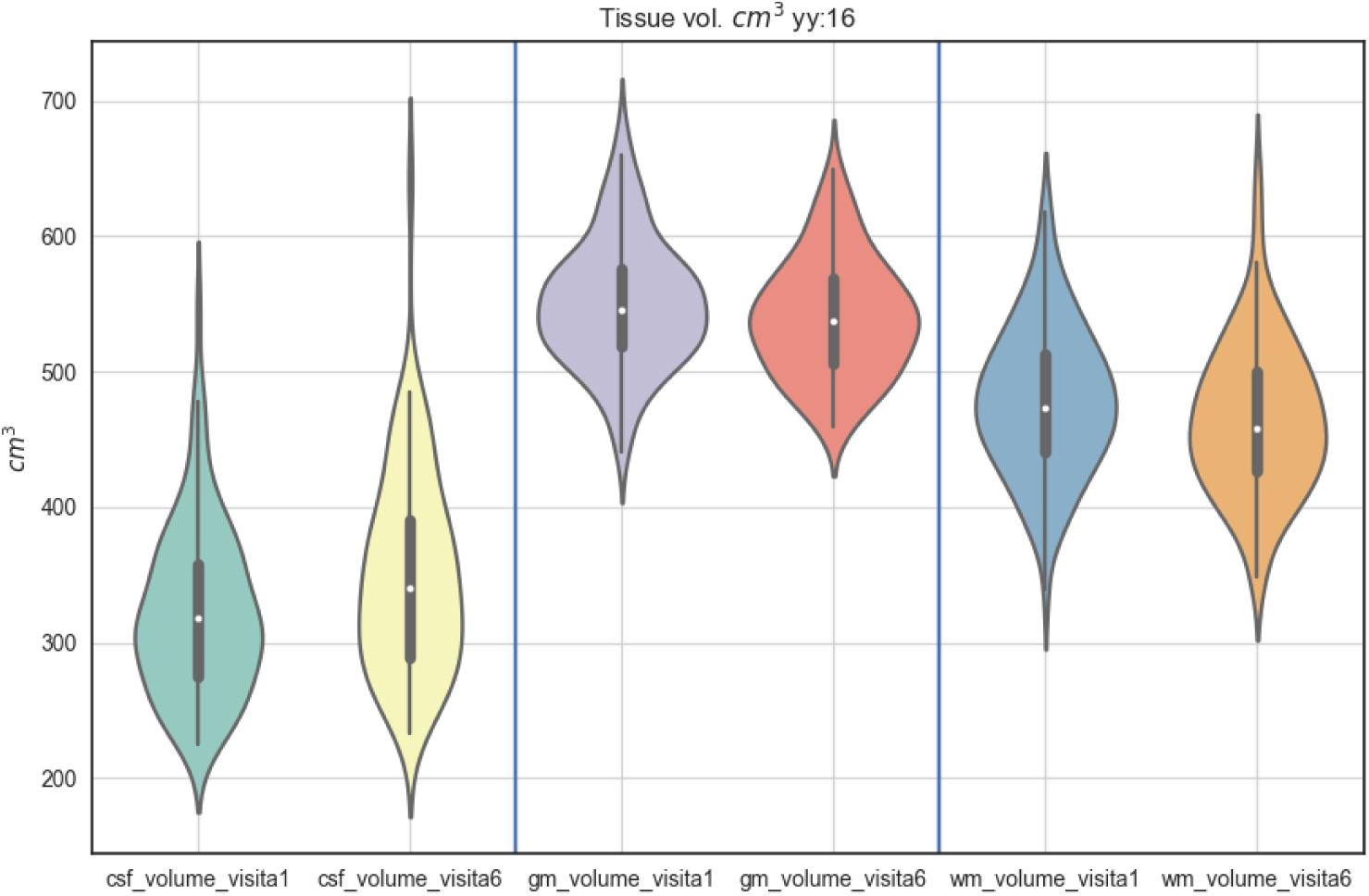
The plot shows the tissue based segmentation results. For each type of tissue we plot the estimated volume in year 1 and year 6. Left CSF volume in first and last years, center gray matter volume in first and last year and right white matter volume for first and last year.

As the brain ages, it increases its elasticity while keeping constant its porosity [Lasheras, 2007], the concentration of proteins also decreases with age which can be explained by the increment in volume observed here [Chen et al., 2012]. While it has been suggested that cerebrospinal fluid production is reduced in healthy aging [May et al., 1990], in a more recent study [Gideon et al., 1994] researchers argue that differences found in human CSF production rates may be caused by interindividual factors other than age. More uncontroversial is the decrease in cerebrospinal fluid pressure with older age [Fleischman et al., 2012] this would explain the enlargement of the brain ventricles that our estimates show.

Figure 43 shows the relationship between the volume of each tissue in year 1 and year 6. As expected, there is a strong linear relationship between the estimated volumes at two different time points. This means that healthy aging progresses monotonically and not for example, “catastrophically” in the mathematical interpretation of René Thom [Rosen, 1979], [Zeeman, 1979], [Thom, 2018]. Along these lines it would be of interest to investigate the points that fall outside the contour areas shown in the figure.

**Figure 43:**
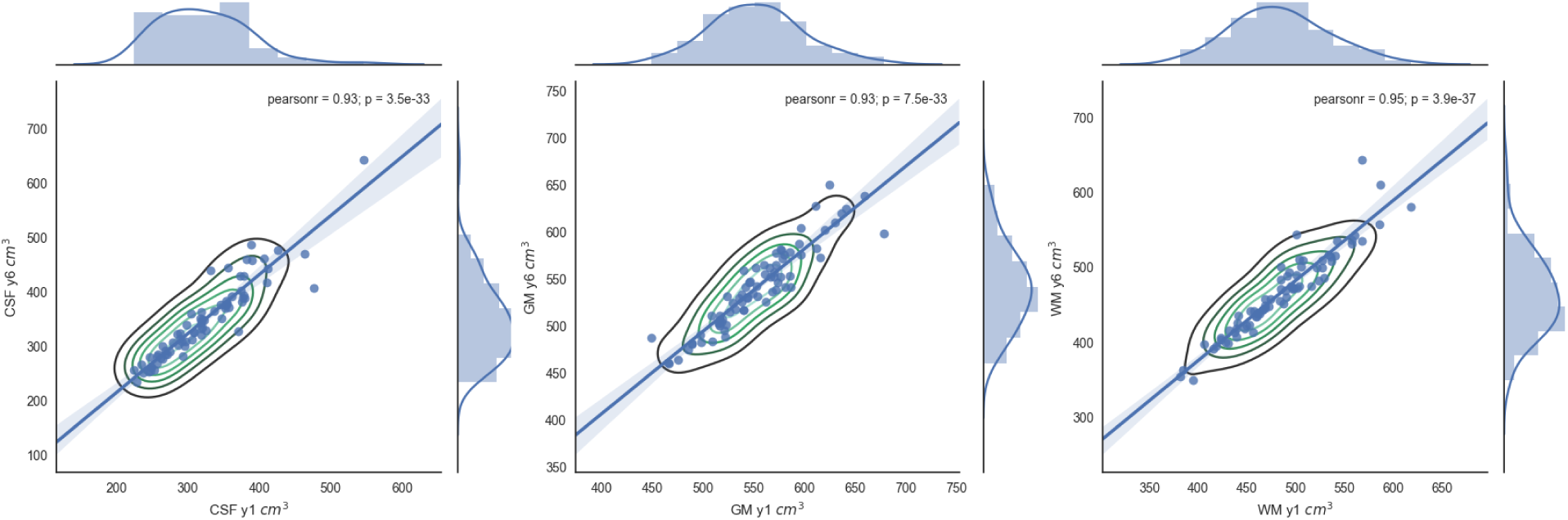
Left, relationship between the CSF volume in year 1 (x-axis) and in year 6 (y-axis). Center, relationship between the gray matter volume in year 1 (x-axis) and in year 6 (y-axis). Right, relationship between the white matter volume in year 1 (x-axis) and in year 6 (y-axis). The linear correlation is very strong in all cases (0.93, 0.93 and 0.95).

Figure 44 shows the relationship between the atrophy of tissues as well as the profiles of each tissue atrophy in the margins. Thus, on the left, the joint plot of gray matter atrophy and white matter atrophy, center, between gray matter atrophy and CSF atrophy and on the right between white matter and CSF. The figure indicates that there is no any particularly strong linear correlation between the volume atrophy of the brain tissues.

**Figure 44:**
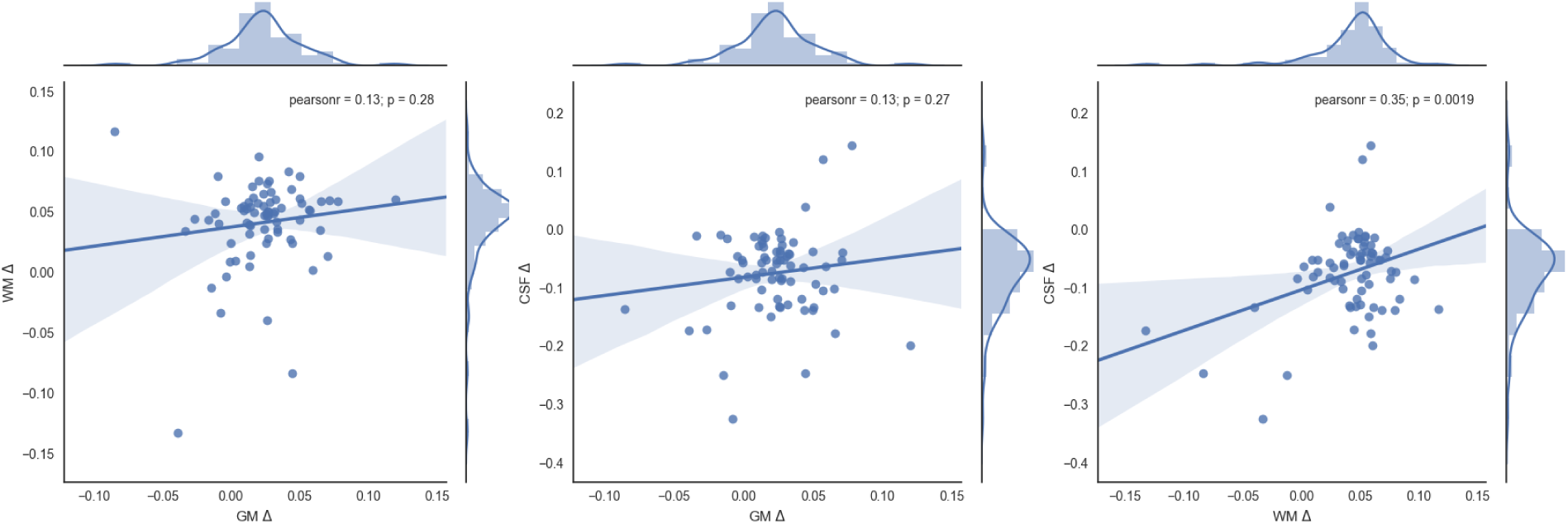
On the left, relationship between the gray matter atrophy and white matter atrophy. Center, relationship between gray matter and CSF atrophy. Right, relationship between white matter and CSF atrophy.

### Segmentation of brain subcortical structures

Figure 45 shows the segmentation of the subcortical structures: Thalamus, Putamen, Hippocampus, Accumbens, Pallidum, Caudate and Amygdala for one subject of the *Vallecas Project*.

**Figure 45:**
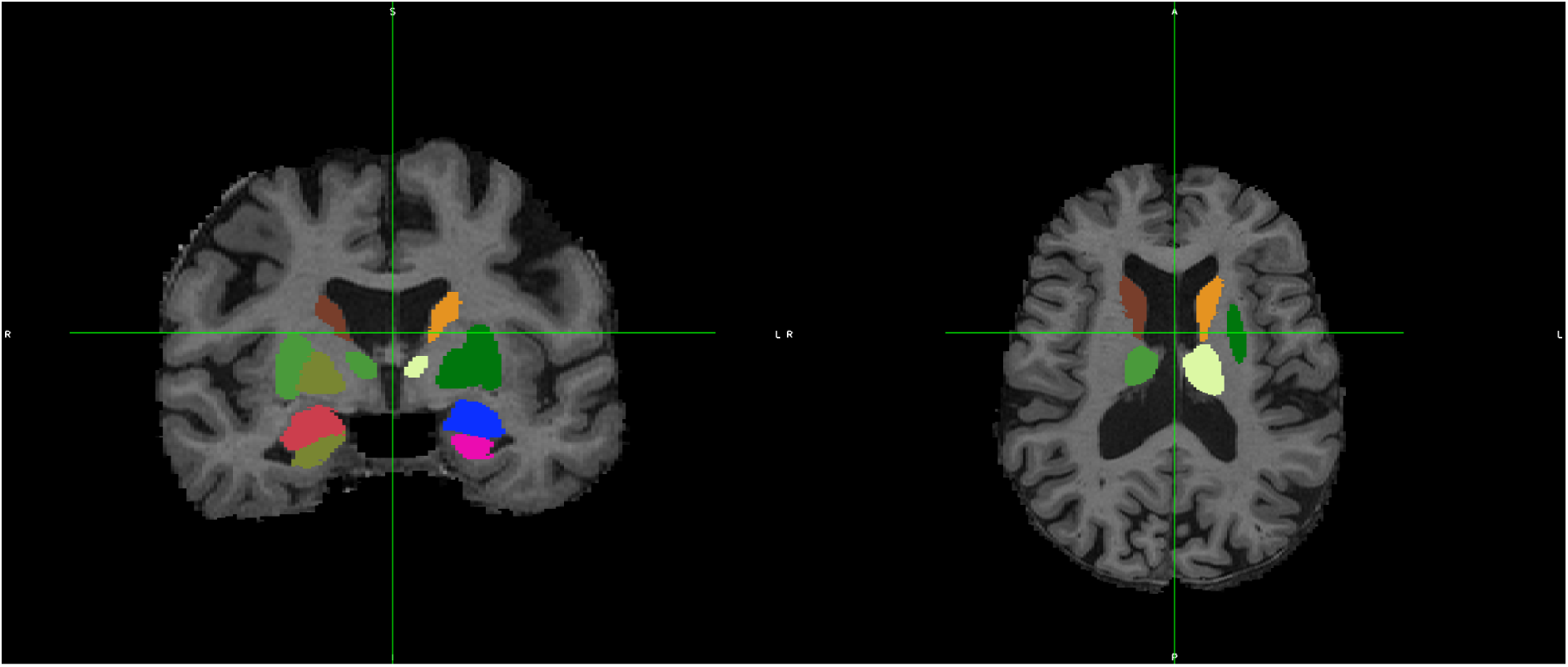
Segmentation of subcortical structures Thalamus, Putamen, Hippocampus, Accumbens, Pallidum, Caudate, Amygdala of one subject of the *Vallecas Project*.

Figure 46 shows the atrophy calculated as the standardized difference in volume between year 1 and year 6 for the different subcortical structures (Equation 1).

**Figure 46:**
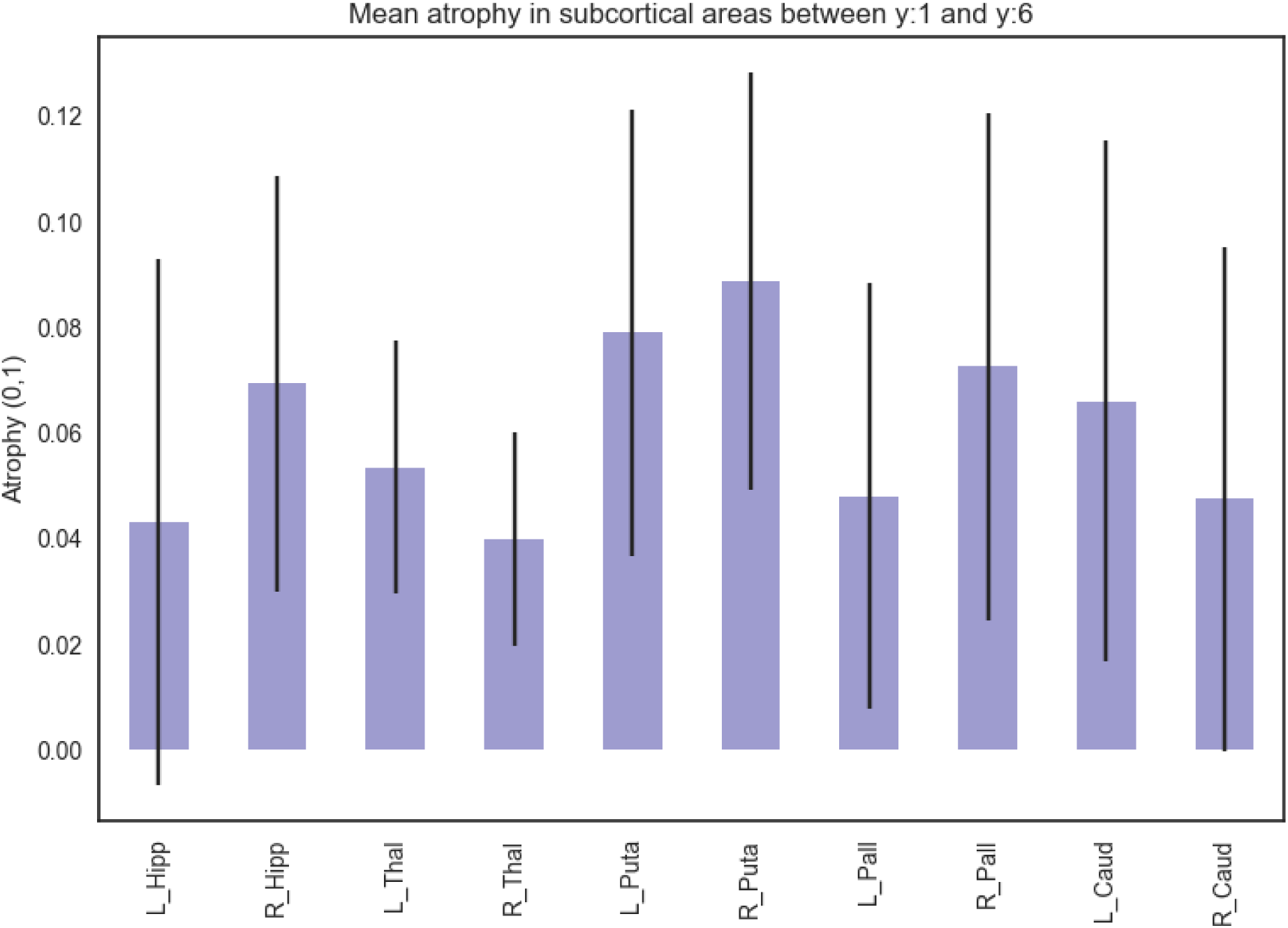
Mean atrophy estimated between year 1 and year 6 for the subcortical areas bilaterally (Thalamus, Putamen, Hippocampus, Accumbens, Pallidum, Caudate, Amygdala). Interestingly, the left hippocampus undergoes the least atrophy on average, the largest atrophy occurs in the Putamen. The mean atrophy has been calculated over a total of 78 subjects, it may be possible to extend this number by using a more lenient policy of outliers removal and of course with alternative segmentation methods.

Figures 47, 48, 49, 50, 51, 52, 53 show the correlation between the volume in year 1 and volume in year 6 for the Caudate, Putamen, Pallidum, Thalamus, Accumbens, Amygdala and Hippocampus, respectively.

**Figure 47:**
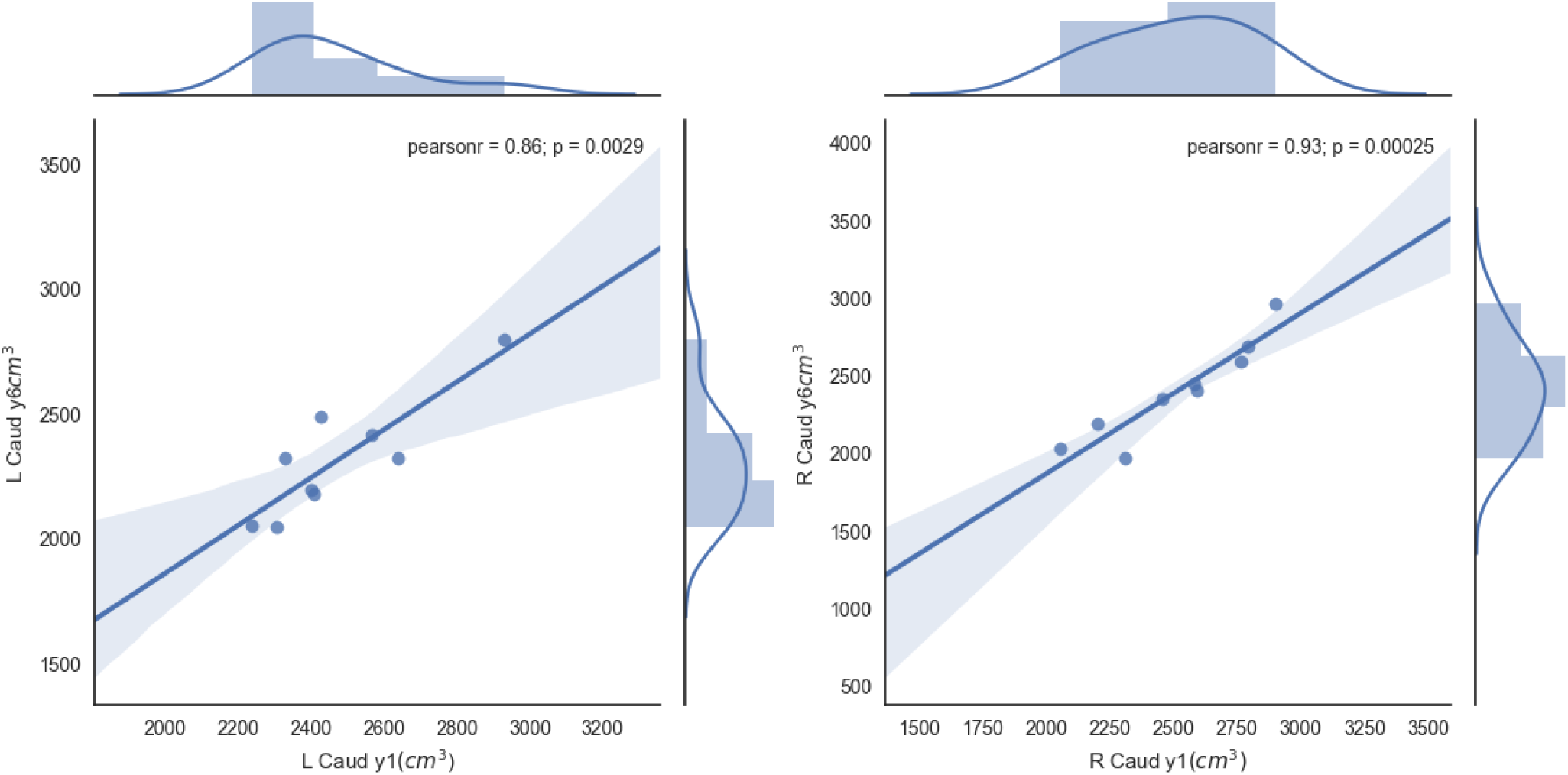
Linear correlation for the caudate (both sides) volume change in 6 years time.

**Figure 48:**
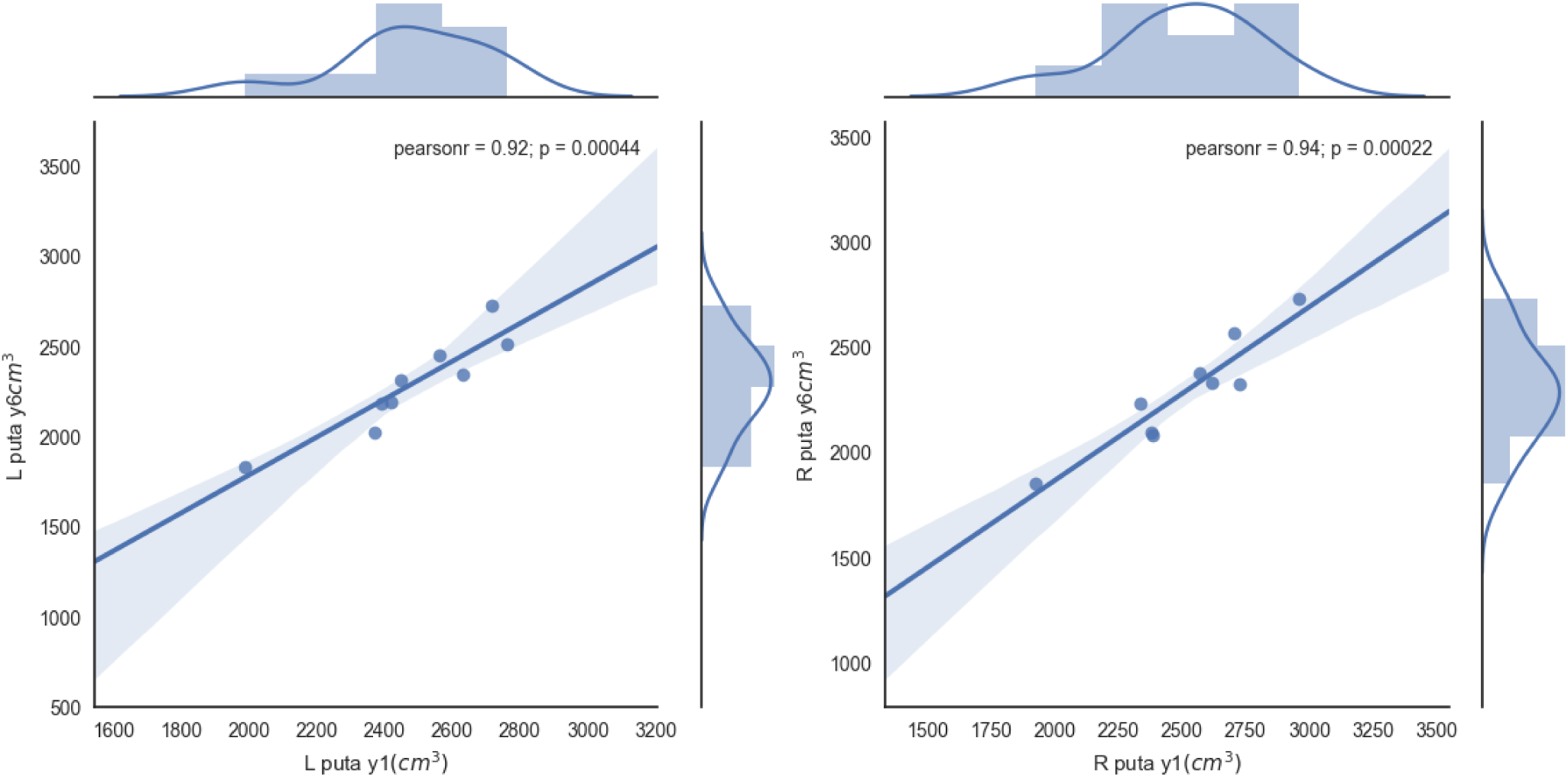
Linear correlation for the putamen (both sides) volume change in 6 years time.

**Figure 49:**
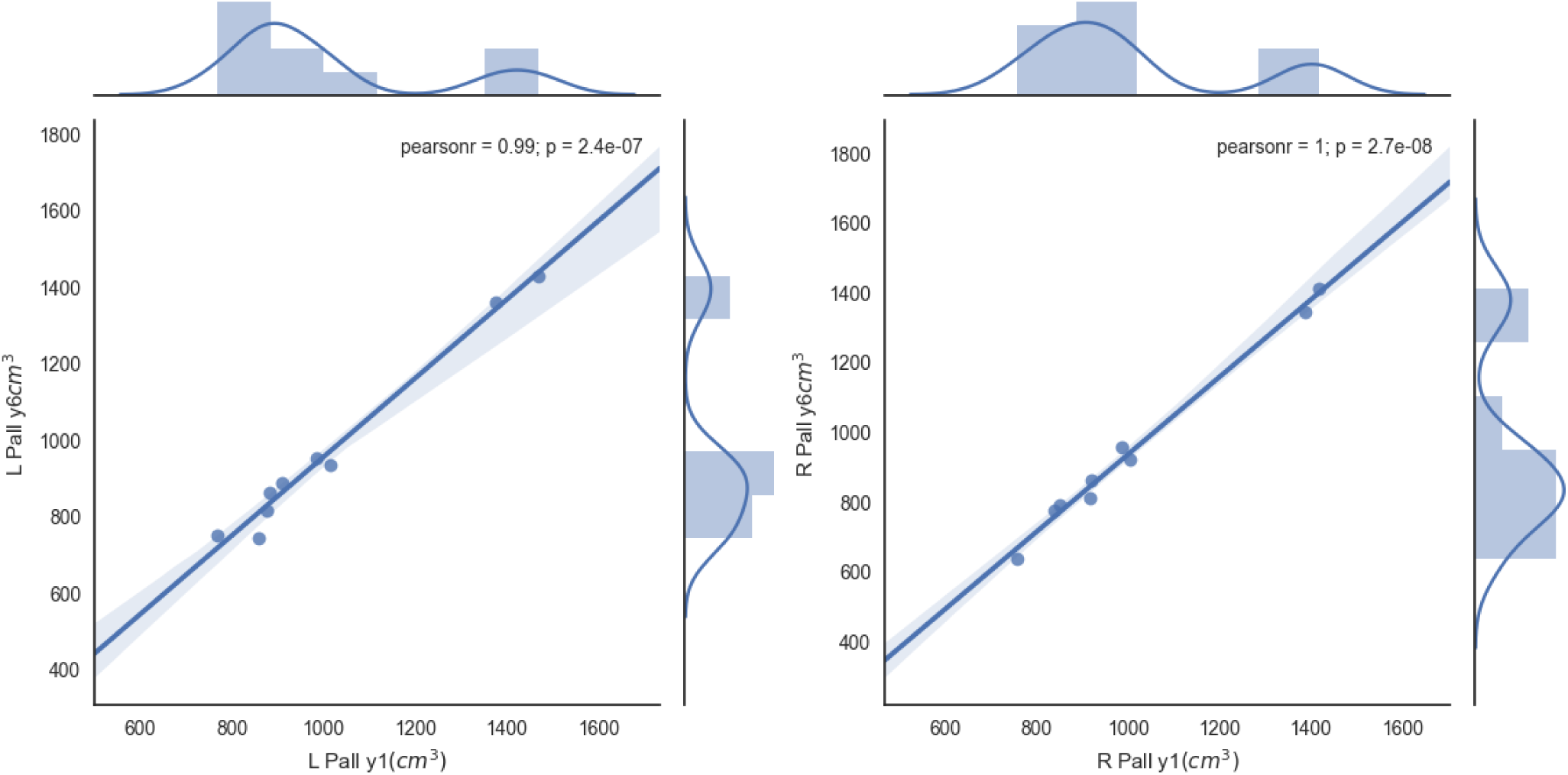
Linear correlation for the pallidum (both sides) volume change in 6 years time.

**Figure 50:**
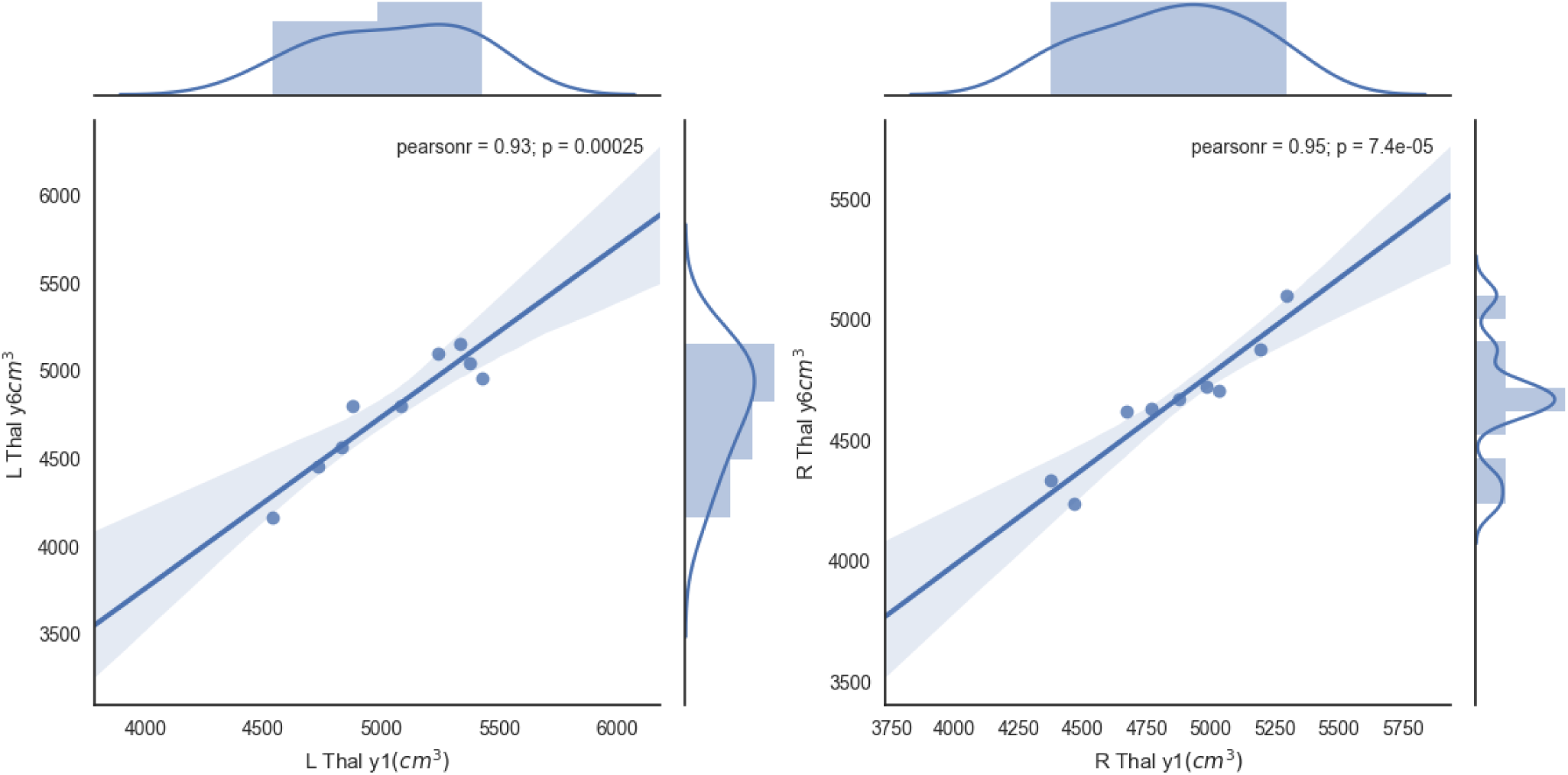
Linear correlation for the thalamus (both sides) volume change in 6 years time.

**Figure 51:**
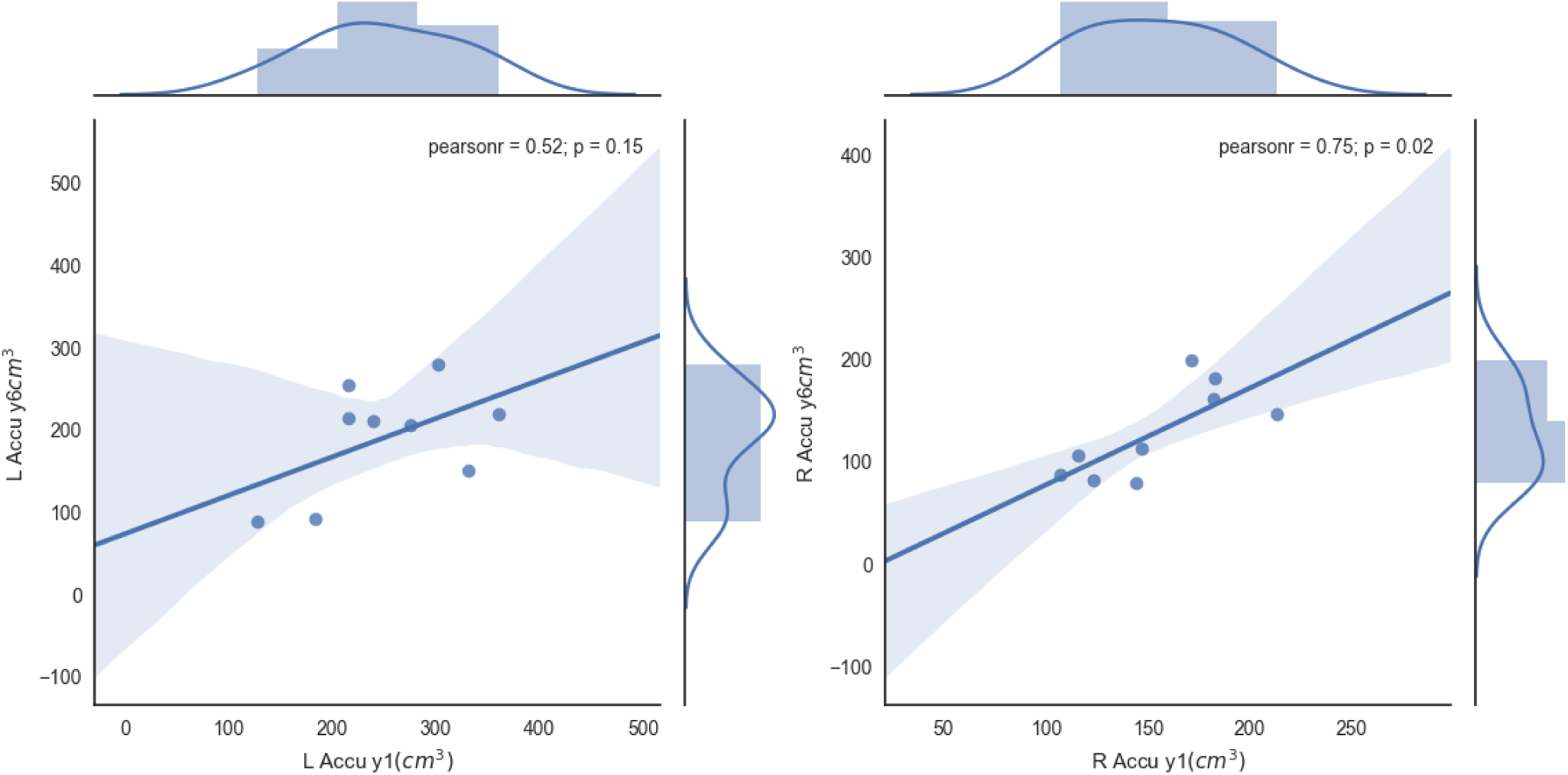
Linear correlation for the accumbens (both sides) volume change in 6 years time.

**Figure 52:**
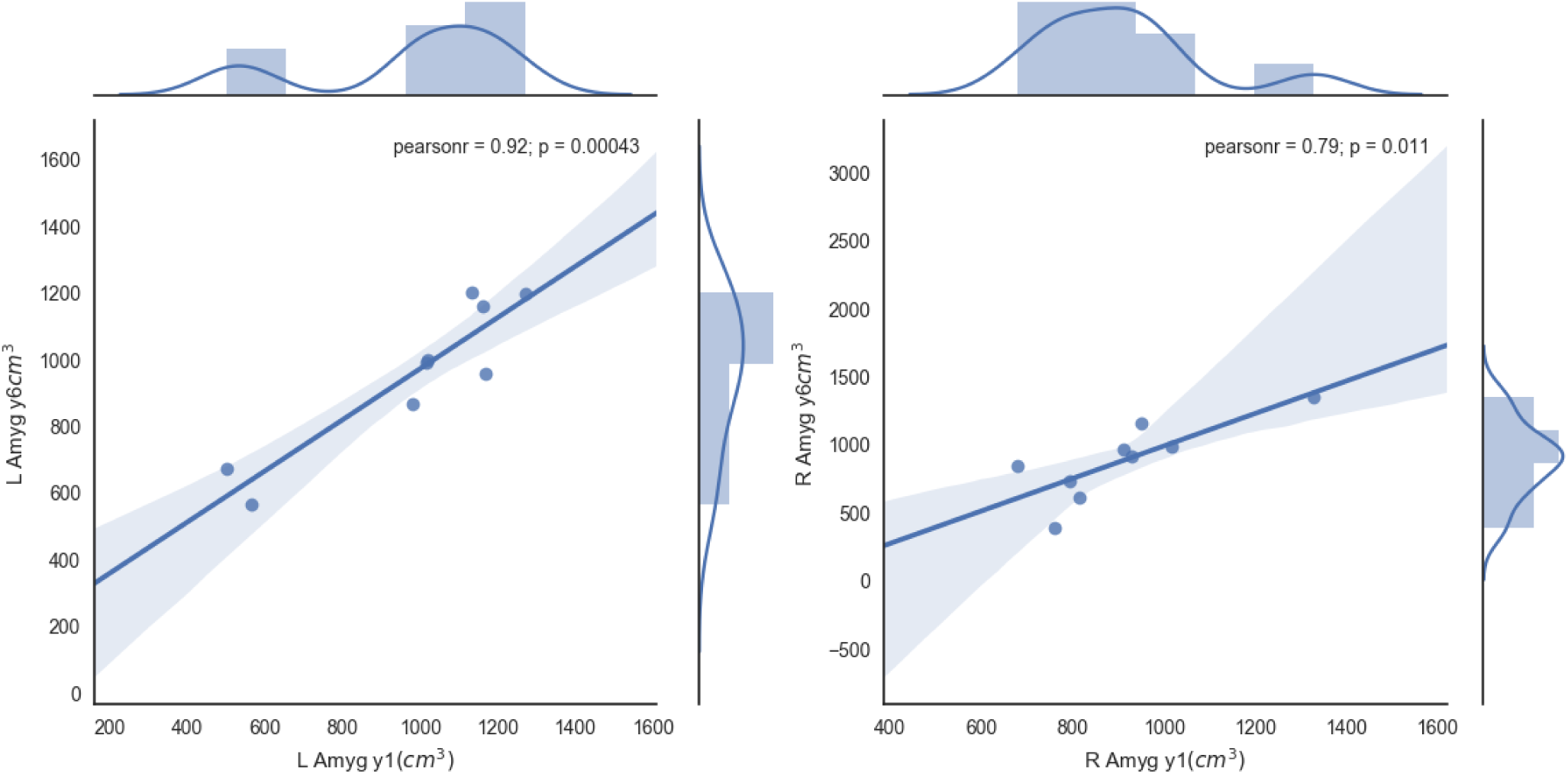
Linear correlation for the amygdala (both sides) volume change in 6 years time.

**Figure 53:**
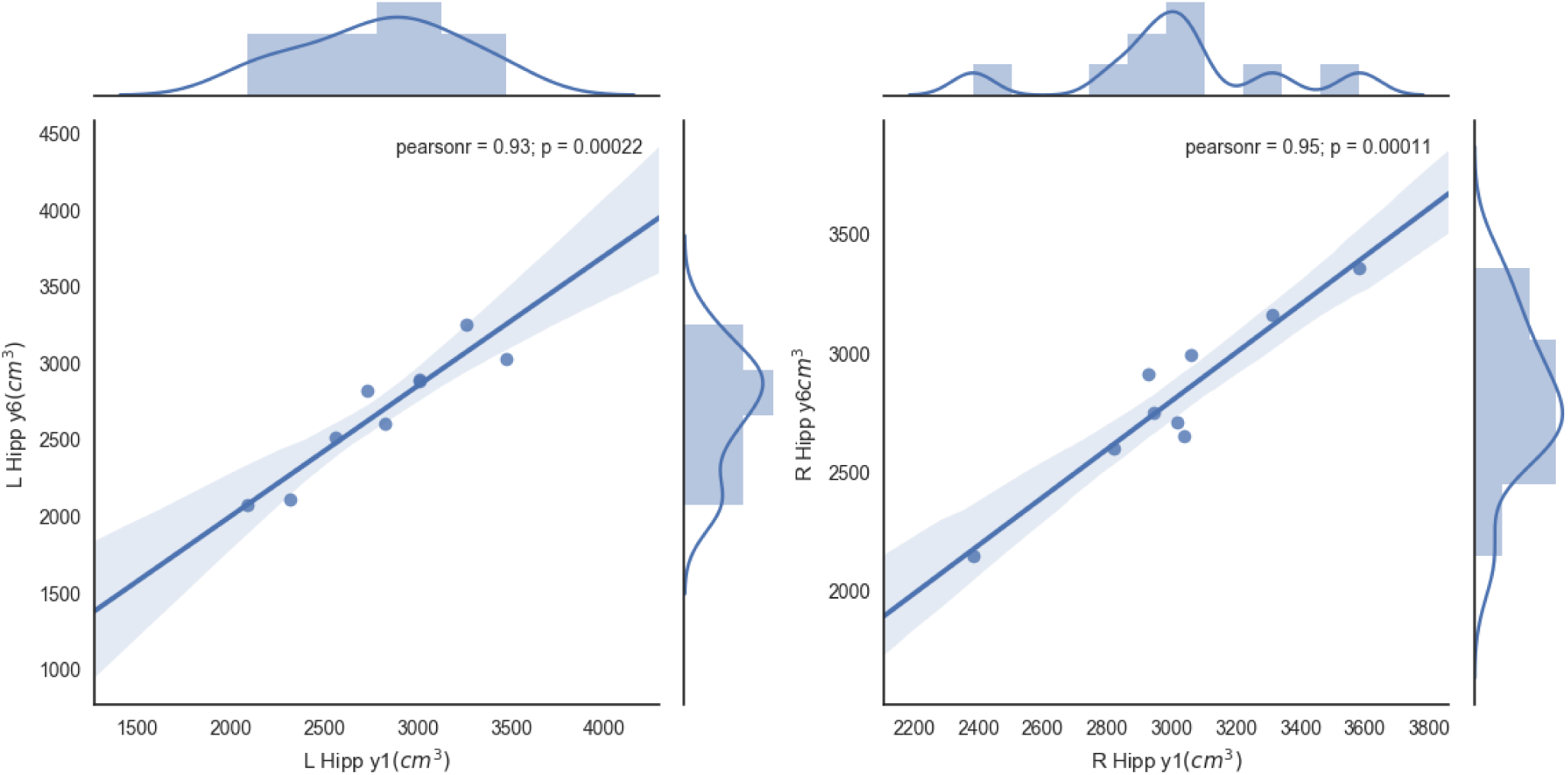
Linear correlation for the hippocampus (both sides) volume change in 6 years time.

Figure 54 shows the Pearson correlation between the atrophy produced between years 1 and 6 in the different subcortical areas.

**Figure 54:**
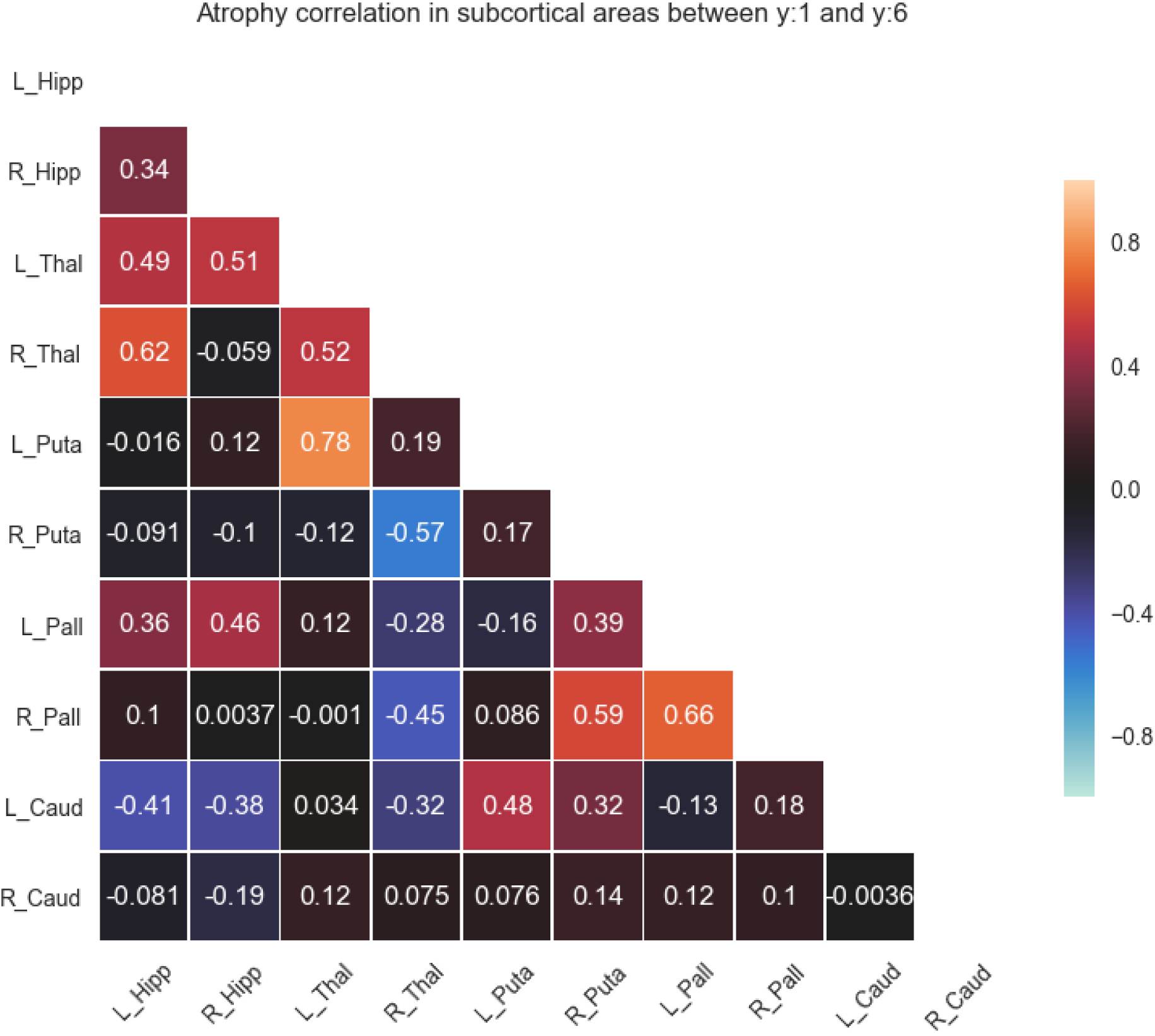
Pearson correlation of the estimated atrophy between year 1 and year 6 for all the studied subcortical areas bilaterally (Thalamus, Putamen, Hippocampus, Accumbens, Pallidum, Caudate, Amygdala).

## 6 Results

The study presented here compiles a longitudinal analysis of *The Vallecas Project*. Although the idea was to perform an exploratory analysis, it is still possible to draw conclusions and recommendations. We will focus on the results shown in Table 12 and Table 13 and in the insights produced by the brain segmentation procedure discussed in Section 5.

Bivariate analysis of cross sectional variables and conversion to MCI using statistical significance tests (Fisher and 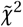) show a small set of variables with statistical relevance: *APOE, thyroid disorders* and variables coding for social activities such as the use of new technologies, going to church and participation in cultural events and public entertainment. The ANOVA tests for non categorical variables in Table 13 show statistical relevance for MCI conversion in dietary practices, in particular diet with a strong fat component and less so for protein based diets.

Aside from considerations of the profusely documented problems with statistical significance tests [Ioannidis, 2005], [Simmons et al., 2011], [Veresoglou, 2015],[Greenland et al., 2016], [Stupple et al., 2019],[Hurlbert et al., 2019], there are at least two important caveats. First, the classification problem of converters to MCI versus non converters is unbalanced, that is, the class of converters is underrepresented. Second, we are only studying two dimensional relationships, on the other hand, cognitive decline which is the process that conversion to MCI is trying to map is a complex phenomenon involving more complicated patterns than bivariate mapping.

In a forthcoming study we specifically tackle the feature selection problem using machine learning. Specifically we build a random forest classifier [Breiman, 2001] from which we will identify the most informative features, we call to this set features the *Vallecas Index*.

It is also possible to draw some interesting conclusions from the brain segmentation results shown in Section 5. First, as expected the volume of the brain decreases with aging. Figure 39 shows a decrease in the mean volume of the brain in year 6 compared to year 1. Furthermore, the kernel density estimation included in the figure indicates a marked difference in the shape of the distributions of each year. The figure suggests that aging makes the brain volume distribution more entropic or more flat. (See the shape of the violin plot in year 6 compared to year 1).

The tissue-based segmentation obtained the volume estimates of cerebrospinal fluid (CSF), gray matter and white matter for both years 1 and 6 in healthy subjects (diagnosed as healthy in the last year) shown in Figure 42, confirms that aging increases brain cavities which are filled with cerebrospinal fluid. As Figure 42 shows, the volume of CSF increases going from year 1 to year 6. On the other hand, the volume of both white and gray matter goes down as aging progresses. We also plot in Figure 43 the relationship between the tissue volume in year 1 and year 6. In all cases there is strong linear correlation, this can be interpreted as healthy aging progressing monotonically, that is, without big leaps. The more it deviates from linearity the less deterministic or entropic aging would be. This discussion will be expanded and made more clear in a forthcoming work that uses the full time series (*t =* 1, 2, 3, 4, 5 and 6) rather than the first and last time points (*t =* 1, 6). The atrophy between tissues is calculated as the standardize difference between the volume in year 1 and the volume year 6 (Equation 1) is shown in Figure 44. The Pearson correlation gives us 0.13 for both Δ(*GM*) *∼* Δ(*W M*) and Δ(*GM*) *∼* Δ(*CSF*) and 0.35 for Δ(*W M*) *∼* Δ(*CSF*).

Finally, the segmentation of brain subcortical structures including Thalamus, Putamen, Hippocampus, Accumbens, Pallidum, Caudate and Amygdala [Patenaude et al., 2011] throws some important information to be consider and replicated using alternative segmentation techniques [Despotović et al., 2015], [Zhang et al., 2018]. First, the mean atrophy per subcortical structure for subjects diagnosed as healthy in year 6 is shown in Figure 45. The left hippocampus is the structure that has on average the least atrophy (4%). The Putamen, both left an right, has the largest atrophy (8%). According to this result the left hippocampus would be a subcortical structure more stable, in terms of volume loss, than others subcortical areas during healthy aging. Nevertheless, this result must be interpreted with caution, a different outliers removal procedure and different segmentation algorithm could produce modifications in the results. In the future we will test and extend on this result to study the volumetric progression of the hippocampus with alternative segmentation procedures that are able to identify the hippocampal subfields as well.

The linear correlation for each substructure is also obtained (Figures 47 to 53). We find a strong linear dependency between subcortical structure volumes between year 1 and year 6. The rationale behind this finding has been already discussed above -aging is a monotonic process. It might be mentioned that the Pearson correlation of the volume changes in the nucleus accumbens (both sides) and the right amygdala is significantly lower than for the rest of the structures (*p >* 0.01). Whether this finding is relevant to brain senescence or a side effect of the segmentation algorithm used, will be investigated using alternative segmentation techniques that take advantage of the scanner spatial resolution, producing amygdala nuclei volume estimates [Iglesias et al., 2015].

A cogent understanding of how senescence affects brain function will be incomplete without brain functional data. We explore changes in brain connectivity using fMRI collected in *The Vallecas Project* in an upcoming work.

## Footnotes

1 We will use indistinctly *Vallecas Project* and its vernacular *Proyecto Vallecas*.

2 We use the term “Caucasian” as it is used in the referred work, however this labeling can be meaningless due to the large heterogeneity and non specificity in the population that the label is supposed to name [Bhopal and Donaldson, 1998]

